# The genetic history of Portugal over the past 5,000 years

**DOI:** 10.1101/2024.09.12.612544

**Authors:** Xavier Roca-Rada, Roberta Davidson, Matthew P. Williams, Shyamsundar Ravishankar, Evelyn Collen, Christian Haarkötter, Leonard Taufik, António Faustino Carvalho, Vanessa Villalba-Mouco, Daniel R. Cuesta-Aguirre, Catarina Tente, Álvaro M. Monge Calleja, Rebecca Anne MacRoberts, Linda Melo, Gludhug A. Purnomo, Yassine Souilmi, Raymond Tobler, Eugénia Cunha, Sofia Tereso, Vítor M. J. Matos, Teresa Matos Fernandes, Anne-France Mauer, Ana Maria Silva, Pedro C. Carvalho, Bastien Llamas, João C. Teixeira

**Affiliations:** Australian Centre for Ancient DNA, The Environment Institute, School of Biological Sciences, The University of Adelaide, Adelaide, South Australia, Australia; Faculty of Arts and Humanities, University of Coimbra, Largo da Porta Férrea, 3004-530 Coimbra, Portugal; Department of Biology, Pennsylvania State University, University Park, PA 16802, USA; Technology Advancement Unit, Genetics and Molecular Pathology, SA Pathology, Adelaide, South Australia, Australia; Laboratory of Genetic Identification & Human Rights (LABIGEN-UGR), Department of Legal Medicine, Faculty of Medicine, University of Granada, Av. Investigación 11 – PTS – 18016, Granada, Spain; CEAACP - Centro de Estudos de Arqueologia, Artes e Ciências do Património, UAlg Universidade do Algarve, F.C.H.S., Campus de Gambelas, 8000-117 Faro, Portugal; Institute of Evolutionary Biology, CSIC-Universitat Pompeu Fabra, Barcelona, Spain; Research Group in Biological Anthropology (GREAB), Biological Anthropology Unit, Department of Animal Biology, Vegetal Biology and Ecology, Universitat Autònoma de Barcelona, Bellaterra, Spain; Instituto for Medieval Studies, Nova University of Lisbon, Portugal; Australian Research Council Centre of Excellence for Australian Biodiversity and Heritage (CABAH), School of Biological Sciences, The University of Adelaide, South Australia, Australia; Mochtar Riady Institute for Nanotechnology, Universitas Pelita Harapan, Tangerang, Indonesia; Research Centre for Anthropology and Health (CIAS), Department of Life Sciences, University of Coimbra, Calçada Martim de Freitas, 3000-456 Coimbra, Portugal; HERCULES Laboratory and IN2PAST, University of Évora, Évora, Portugal; National Centre for Indigenous Genomics, John Curtin School of Medical Research, Australian National University, Canberra, Australian Capital Territory, Australia; Indigenous Genomics, The Kids Research Institute Australia, Adelaide, South Australia, Australia; Evolution of Cultural Diversity Initiative, School of Culture, History and Language, The Australian National University, Canberra, ACT, Australia; Australian Research Council Centre of Excellence for Indigenous and Environmental Histories and Futures, Australian National University, Canberra, ACT, Australia; Centre for Functional Ecology, Laboratory of Forensic Anthropology Department of Life Sciences, University of Coimbra, Calçada Martim de Freitas, 3000-456 Coimbra, Portugal; National Institute of legal Medicine, Lisbon, Portugal; Instituto de Estudos Medievais, FCSH-NOVA, Lisboa, Portugal; School of Sciences and Technology, Department of Biology, University of Évora, Portugal; Centre for Archaeology, University of Lisbon, Portugal; Centre for Interdisciplinary Studies of the University of Coimbra, Coimbra, Portugal

**Keywords:** ancient DNA, Portuguese populations, Iberia, paleogenomics, population genetics, archaeology, molecular anthropology

## Abstract

**Background:** Recent ancient DNA studies uncovering large-scale demographic events in Iberia have focused primarily on Spain, with limited reports for Portugal, a country located at the westernmost edge of continental Eurasia. Here, we introduce the largest collection of ancient Portuguese genomic datasets (n = 68) to date, spanning 5,000 years, from the Neolithic to the 19^th^ century.

**Results:** We found evidence of patrilocality in Neolithic Portugal, with admixture from local hunter-gatherers and Anatolian farmers, and persistence of Upper Paleolithic Magdalenian ancestry. This genetic profile persists into the Chalcolithic, reflecting diverse local hunter-gatherer contributions. During the Bronze Age, local genetic ancestry persisted, particularly in southern Iberia, despite influences from the North Pontic Steppe and early Mediterranean contacts. The Roman period highlights Idanha-a-Velha as a hub of migration and interaction, with a notably diverse genetic profile. The Early Medieval period is marked by Central European ancestry linked to Suebi/Visigoth migrations, adding to coeval local, African, and Mediterranean influences. The Islamic and Christian Conquest periods show strong genetic continuity in northern Portugal and significant African admixture in the south, with persistent Jewish and Islamic ancestries suggesting enduring influences in the post-Islamic period.

**Conclusions:** This study represents the first attempt to reconstruct the genetic history of Portugal from the analysis of ancient individuals. We reveal dynamic patterns of migration and cultural exchange across millennia, but also the persistence of local ancestries. Our findings integrate genetic information with historical and archaeological data, enhancing our understanding of Iberia’s ancient heritage.

## 1. Background

The Iberian Peninsula, located at the westernmost edge of continental Europe, is a geographically *quasi*-isolated region that offers a unique perspective on ancient patterns of human migration into a European *cul-de-sac*. Specifically, this geographical setting provides a valuable opportunity to investigate how relative local isolation and continental patterns of human movement influenced the persistence and arrival of genetic ancestries to the region, as well as the overall structure and stability of local population networks.

The widespread and archaeologically confirmed presence of anatomically modern humans in Portugal dates to the Lower Paleolithic, with the oldest human fossils, found in the Aroeira cave, dated to approximately 400,000 years ago (1). However, the importance of Iberia for European human evolution is underscored by its role as a refugium during the Last Glacial Maximum (LGM), as harsh climatic conditions led to the contraction of human habitable areas across Europe (2–7). As the climate warmed and the glacial ice receded around 14,000 years ago, human populations began expanding across Europe (2–6,8–12). This period marked the gradual transition from a hunter-gatherer lifestyle to more settled forms of subsistence, as evidenced by the presence of Mesolithic necropolises associated with shell middens (i.e., *concheiros*) in present-day Portugal (13).

It was not until the Neolithic period (∼5,700/5,600 BCE) that significant changes occurred in Iberia with the introduction of agriculture, herding, and permanent settlements. This shift from hunting and gathering to farming was primarily driven by population movements from western Anatolia into Europe (14–18). Specifically, the maritime spread of human groups along the western Mediterranean carrying *impressed* potteries, such as the Cardial group (19,20), reached central-southern Portugal by around 5,500 BCE (21). Recent genetic studies revealed dynamic population movements and a notable persistence of hunter-gatherer ancestry in Neolithic Iberian communities, higher than in Central Europe due to admixture along the migration route and additional local contributions (22–24).

Advancements in technology (e.g., copper and gold metallurgy) along with economic and trade intensification resulted in the emergence of more complex forms of social organization during the Chalcolithic period (around 3,000-2,000 BCE) (25,26). These societies were characterized by diverse regional cultures that, despite interactions, preserved unique cultural traits, likely due to the Peninsula’s geographical isolation. However, changes observed in the Bronze Age (beginning ∼2,000 BCE)—such as the development of bronze metallurgy and the emergence of social stratification and complex societies—were accompanied by Steppe-related changes in the genetic ancestry of North and Central Europeans and, to some extent, Iberians (23,27–33). While the existence of Bronze Age trade networks connecting the Iberian Peninsula with the Mediterranean and Atlantic regions, including North Africa, have been described (34,35), large-scale genetic evidence of such interactions remains limited, with only sporadic cases observed (23,32).

During the Iron Age, starting around 800 BCE, the introduction of iron technology significantly advanced agricultural and warfare practices (36). While material culture associated with the Celts in northern, western, and central Iberia, as well as with Eastern Mediterraneans, particularly the Phoenicians, has been found in southwestern Iberia, there is no evidence of demographic changes resulting from admixture with local populations (37,38).

The Roman conquest of the Iberian Peninsula, starting in 218 BCE, integrated present-day Portugal into the Roman Empire, initially divided into Lusitania and Tarraconense, later adding Gallaecia in the north (39). This period saw urbanization, infrastructure development, and economic growth driven by Mediterranean trade and large-scale mineral exploitation (40). Roman culture, law, and the Latin language spread throughout the region, gradually integrating indigenous populations, while settlers from other parts of the Empire, including Europe, North Africa, and the Eastern Mediterranean, arrived in Western Iberia (40).

In the 5^th^ century, the arrival of Germanic tribes led to the collapse of Roman power in the Iberian Peninsula (41,42). The Suebi dominated much of the northwest, corresponding largely to modern-day Portugal, until the Visigoths politically unified the Peninsula in 585 CE (41,42).

In the early 8^th^ century, Islamic tribes from North Africa conquered most of the Iberian Peninsula, incorporating it into the Umayyad Caliphate (43,44). The gradual Christian Conquest from the north, began in the 9^th^ century (45) and culminated in the formation of the County of Portugal and the establishment of the Kingdom of Portugal in 1143 (46). During the late Middle Ages, the Kingdom of Portugal expanded and consolidated its borders when the Algarve (southern Portugal) was conquered by 1249 (46).

During the early Modern Ages (15^th^-18^th^ centuries), Portugal expanded its global empire with colonies in Africa, Asia, and South America, and Lisbon emerged as a major global trade hub (47). The 19^th^ century was marked by political instability, including the Napoleonic occupation (1807–1811), the Liberal Revolution of 1820 and a subsequent civil war that lasted until 1834 (47). The 20^th^ century saw the monarchy overthrown in 1910, leading to the establishment of the Portuguese Republic, which was followed by António Salazar’s authoritarian regime until the peaceful Carnation Revolution restored democracy in 1974 (47). Portugal has since become a stable democratic republic, joining the European Union in 1986 and experiencing steady economic and social development (47).

Over the last few years, several human ancient DNA studies have aimed to uncover large-scale demographic events in the history of the Iberian Peninsula (17,22–24,30–33,48–58), primarily focusing on Spanish populations (n=471 individuals). To date, however, only 51 individual ancient genomes dating from pre-Visigoth times are published for Portugal, which necessarily limits demographic inferences for such a historically rich region (23,24,31,33,53,57).

Here, we present the largest collection of ancient Portuguese genomes to date. We screened 94 samples spanning 5,000 years of human history, including 68 newly generated ancient genomes, from the Neolithic to the 19^th^ century (Fig.1). Overall, we analyzed 590 Iberian ancient genomes (17,22–24,30–33,48–58) from key periods of cultural transition in Portugal and Spain—Prehistory, Antiquity, Middle Ages and Modern Age—to explore population integration and historical movements, thereby enhancing our understanding of the genetic history of Portugal and providing a more complete picture of Iberia’s genetic heritage.

**Figure 1.**
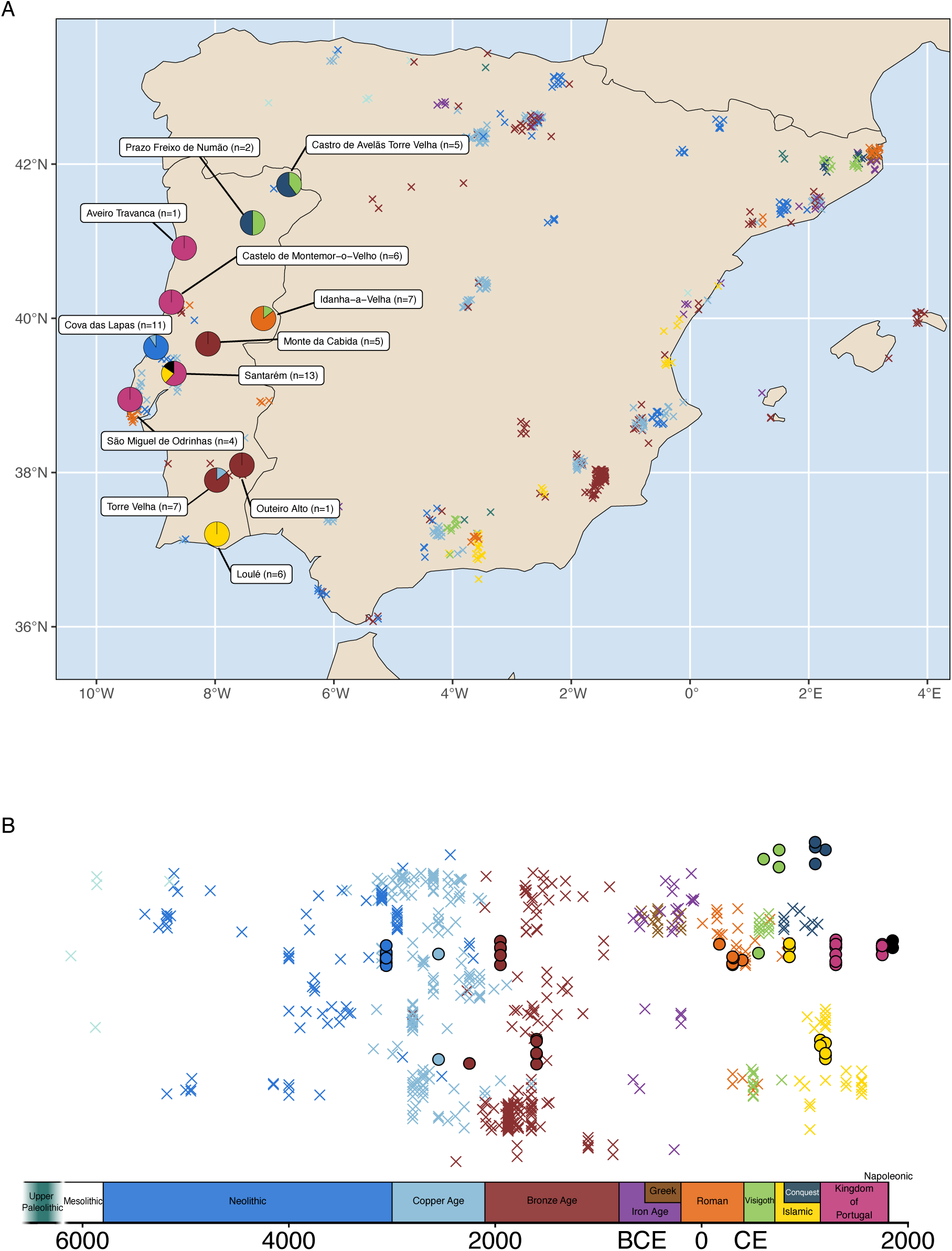
(A) Geographic and (B) chronological distribution of newly generated paleogenomic datasets (n=68; circles) and previously reported paleogenomic datasets (crosses) along a north-south (top to bottom) gradient. Pie charts indicate the archaeological sites analyzed in this study, with colors corresponding to different temporal periods. Random jitter has been applied to other sites for clarity.

## 2. Results

### Neolithic

We analyzed a total of 10 newly sequenced ancient genomes from Neolithic Portugal, including 8 genetic males and 2 genetic females (Table S2) from the archaeological site of Cova_das_Lapas_N (central-western Portugal). A first-degree relationship was identified between two male individuals using whole-genome data (Fig.S1), with both individuals also sharing the Y-chromosome haplogroup I2a1a2a (Table S6) and the mitochondrial DNA (mtDNA) haplogroup U8a1b (Table S5), which indicates they were very likely full brothers. Additionally, a second-degree relationship (Fig.S1) was found between two other male individuals carrying the Y-chromosome haplogroup I2a1a2a1a (Table S6) and the mtDNA haplogroups T2b3 and K1a4a1 (Table S5), suggesting they were either paternal uncle and nephew, paternal grandfather and grandson or half-brothers sharing the same father. In the Multidimensional Scaling (MDS) plot based on 1- *f*3(Mbuti; Ind1, Ind2) (Fig.S2), these related individuals cluster together and separate from the other individuals of Cova_das_Lapas_N indicating a closer genetic relationship. Interestingly, we detected a high diversity of mtDNA haplogroups associated with the LGM Iberian refugium (U5b1c, U5b2b3a, U5b2b, and U8a1b) and the spread of Neolithic farmers into Europe from Southwestern Asia (K1a4a1, J1c1c, T2b3, and T2c1d) (Table S5) compared to the presence of only two distinct Y-chromosome sub-haplogroups from the same clade, representing different levels of resolution (I2a1a2a1a and I2a1a2a as one sub-haplogroup, and I2a1a1a1a1 as another) (Table S6).

Our genome-wide Principal Component Analysis (PCA) computed with present-day and ancient West Eurasians and North Africans revealed that the genetic profile of Cova_das_Lapas_N clusters closely with the broader Iberian Peninsula (Fig.2A). Neolithic Iberians fall between Mesolithic hunter-gatherers and Anatolia_N, and we observed that Cova_das_Lapas_N harbors an admixed genetic profile, comprising both a pre-Neolithic hunter-gatherer component and an Anatolia_N ancestry component (Fig. 2A; Fig. S3B; Fig. S4). This mixed Anatolian-Iberian ancestry was further confirmed through *qpAdm* modeling, which showed that Cova_das_Lapas_N could be modeled as approximately 68% Anatolia_N and 32% Iberian_HG (Fig. 2E; Table S7).

**Figure 2.**
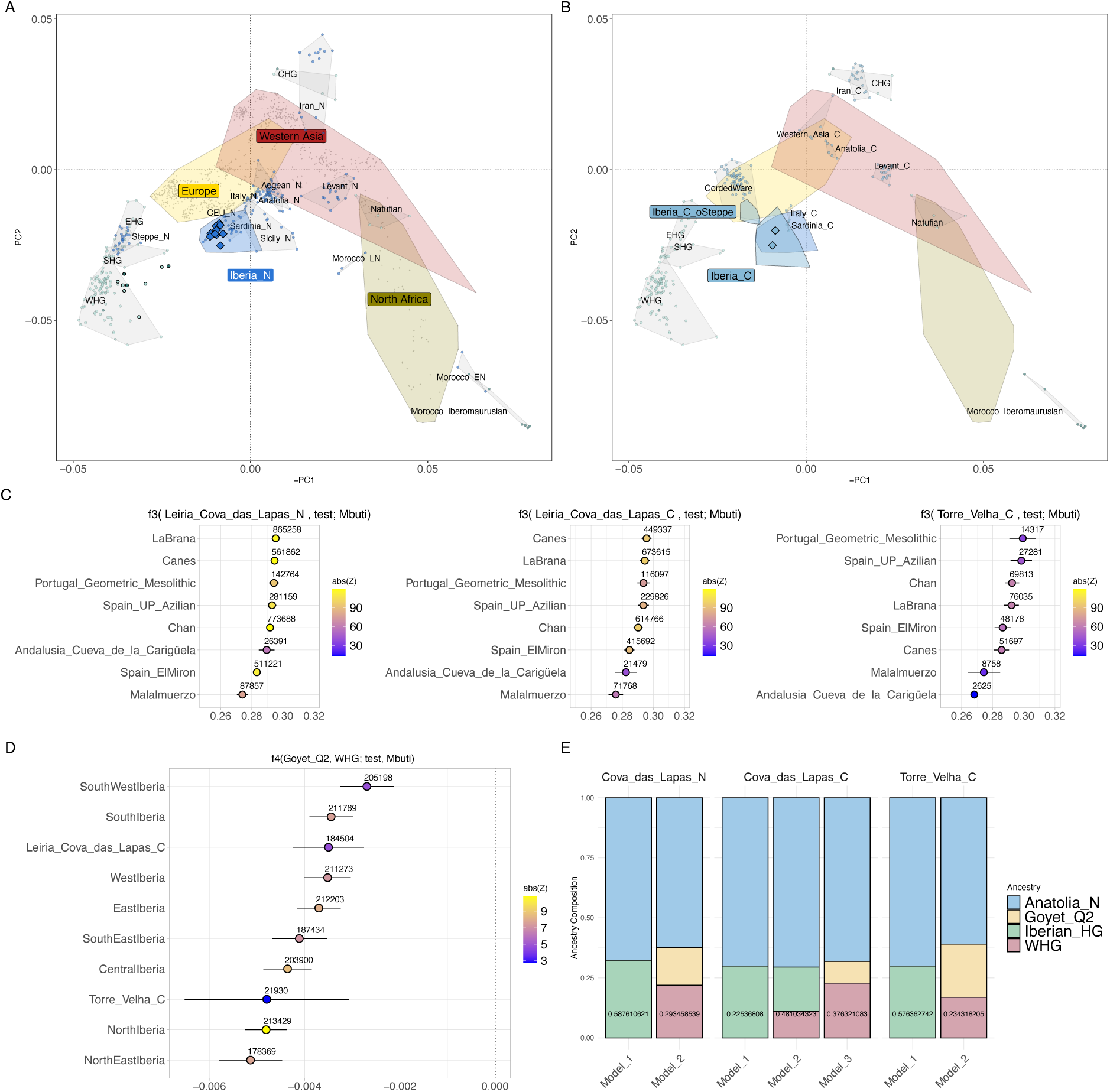
PCA of present-day West Eurasians and North Africans (gray points; overlaid colored polygons represent geographical clusters) with ancient individuals from Iberia and other regions projected onto the first two principal components. Individuals from this study are represented as colored diamonds. Colors indicate different temporal periods as shown in 1B, with a focus on (A) the Neolithic and (B) the Chalcolithic. (C) Outgroup *f3*-statistics of the form *f3*(X, Test; Mbuti), where *X* represents the archaeological site, and *Test* includes Iberian and European Upper Paleolithic and Mesolithic hunter-gatherers. The x-axis shows the *f3*- statistic values, with results displayed as the mean ± 1-SD and colors representing the Z-scores. The numbers above each dot indicate the number of SNPs used for each calculation. (D) *f4*- statistics of the form *f4*(Goyet_Q2, WHG; Test, Mbuti), where *Test* includes various Chalcolithic Iberian geographical regions, Cova_das_Lapas_C, and Torre_Velha_C. The x-axis shows the *f4*-statistic values, with results displayed as the mean ± 1-SD and colors representing the Z-scores. The numbers above each dot indicate the number of SNPs used for each calculation. (E) Ancestry proportions for Cova_das_Lapas_N/C and Torre_Velha_C (y-axis) using different admixture modeling frameworks (x-axis). The p-values are provided inside each column.

In order to determine the closest genetic affinity between Cova_das_Lapas_N and different geographically defined Neolithic Iberian groups, we computed an outgroup *f3*-statistics of the form *f3* (Cova_das_Lapas_N, Test; Mbuti), where the *Test* population rotates through different geographically defined groups and found no geographic correlation (Fig.S3A).

Previous studies have shown that northwestern, southeastern and southwestern Iberian Mesolithic hunter-gatherers retained more ancestry from the LGM period compared to Northern Iberian Mesolithic hunter-gatherers (23,24,59). The LGM ancestry is related to individuals associated with the Magdalenian genetic profile (Goyet_Q2 cluster), whereas most of the ancestry found in northern Iberian Mesolithic hunter-gatherers is associated with post-LGM expansions (Villabruna/Oberkassel cluster) (6). To investigate the hunter-gatherer ancestry of Cova_das_Lapas_N, we computed an outgroup *f3*-statistics of the form *f3*(Cova_das_Lapas_N, Test; Mbuti), where *Test* rotates through different pre-Neolithic hunter-gatherers. In this analysis, we observed that the closest genetic relative is the northwestern Mesolithic individual from Chan do Lindero (Pedrafita do Courel, Lugo, Spain) (Fig.2C). Additionally, *qpAdm* modeling using WHG (Luxembourg_Loschbour and Hungary_EN_HG_Koros) instead of Iberian_HG as a source, required Goyet_Q2 as an additional third component (∼16%) (Fig.2E; Table S7). Accordingly, the Goyet_Q2 ancestry in Cova_das_Lapas_N is higher, and the WHG ancestry is lower, than that observed in other Neolithic Iberian populations (Fig.S3C) despite the lack of detectable geographic clines in these ancestries (Fig.S3D).

When integrating Cova_das_Lapas_N with previously published Neolithic Iberians and modeling ancestry proportions for different geographically defined Neolithic Iberian groups using the frameworks previously described in *qpAdm* (Fig.S5; Table S7), the results generally align with previous findings. However, in some cases, Iberian_HG provides a better model fit than Goyet_Q2 when using WHG, suggesting admixture from both European hunter-gatherers, likely carried west by Anatolian farmers, as well as local hunter-gatherers. Specifically, Goyet_Q2 ancestry is lowest in Northern and Northeastern Iberia, particularly near the Pyrenees. The highest and most significant Goyet_Q2 ancestry (∼12%) is observed in Northwestern Iberia, followed by Central Iberia. This highest proportion is lower than that found in Cova_das_Lapas_N, indicating that local admixture in Cova_das_Lapas_N could be due to a longer period of coexistence, possibly related to earlier Neolithisation, or the prolonged survival of hunter-gatherer populations in relative isolation. Additionally, since Iran_GanjDareh_N is required as a fourth ancestry source in Western Iberia, where Cova_das_Lapas_N is located, may suggest a deeper ancestry carried by Anatolia_N.

### Chalcolithic

To investigate the genetic composition of Chalcolithic Portugal, we combined previously published data (Table S19) with two new individuals, one from Cova_das_Lapas_C (PT_22197; genetically male) and another from Torre_Velha_C (PT_23206; genetically female; southern Portugal). We found that both individuals cluster together in the PCA with previously published Chalcolithic and Neolithic individuals from Iberia (Fig.2B), despite the absence of a geographic correlation as measured by an outgroup *f3*-statistics of the form *f3*(Cova_das_Lapas_C/Torre_Velha_C, Test; Mbuti), where *Test* rotates through different geographically defined Chalcolithic Iberian groups (Fig.S6A-B). Both individuals carried the local U5b mtDNA haplogroup (U5b1i and U5b1 sub-haplogroups, respectively) (Table S5), and Cova_das_Lapas_C carried the same Y-chromosome I2a1a1a1a sub-haplogroup as found in Cova_das_Lapas_N (Table S6). In addition, the MDS plot showed a close genetic affinity between the Neolithic and Chalcolithic individuals from Cova_das_Lapas (Fig.S2).

Whilst Cova_das_Lapas_C and Torre_Velha_C are shown to carry highly similar proportions of Iberian_HG and Anatolia_N in *qpAdm* outgroup-based modeling, ADMIXTURE and *f4*- statistics (Fig.2E; Table S8; Fig.S4; Fig.S6C), the results obtained through outgroup *f3*- statistics indicate Cova_das_Lapas_C shares a closer affinity to the northwestern Iberian Mesolithic hunter-gatherer Chan do Lindero (as observed in Neolithic individuals), while Torre_Velha_C is more closely related to central Portuguese Late Mesolithic hunter-gatherer (CMS001) (Fig.2C). Notably, neither individual shows traces of Steppe ancestry, as indicated by *f4*-statistics of the form *f4*(Anatolia_N, Cova_das_Lapas_C/Torre_Velha_C; Russia_Samara_EBA_Yamnaya, Mbuti) (Fig.3B) and *f4*(Portugal_C, Portugal_N/BA; Russia_Samara_EBA_Yamnaya, Mbuti) (Fig. S6F), which yield extremely low positive values. This finding provides further support to the genetic continuity observed between Neolithic and Chalcolithic populations in present-day Portugal, and to the persistence of discrete genetic ancestry from different hunter-gatherer groups across time.

**Figure 3.**
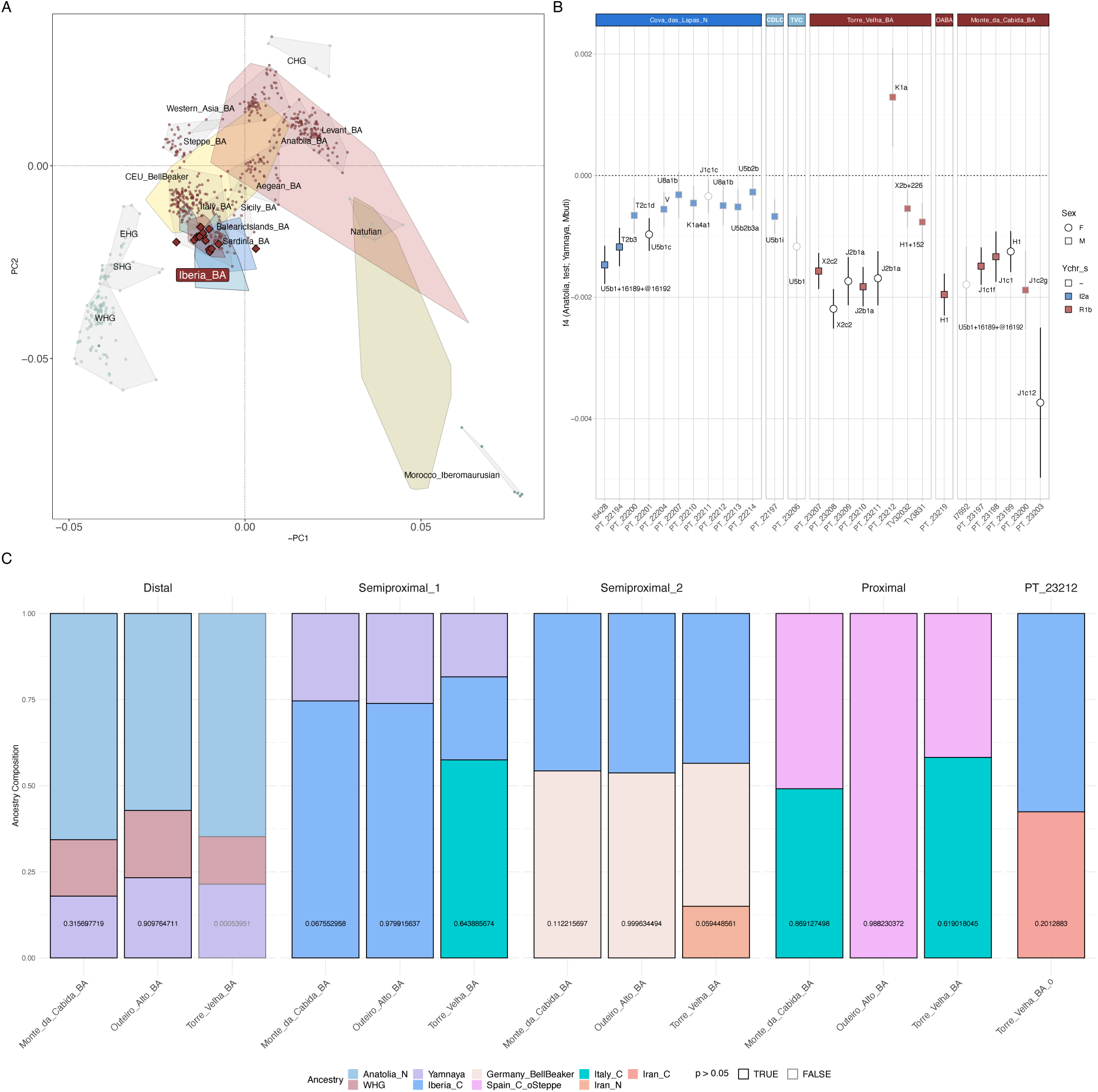
(A) PCA of present-day West Eurasians and North Africans (overlaid colored polygons represent geographical clusters) focusing on the Bronze Age. The ancient individuals from Iberia and other regions were projected onto the first two principal components. Individuals from this study are represented as colored diamonds. Colors correspond to different temporal periods, as shown in 1B. (B) *f4*-statistics of the form *f4* (Anatolia_N, Test; Russia_Samara_EBA_Yamnaya, Mbuti), where *Test* includes Cova_das_Lapas_N, Cova_das_Lapas_C (CDLC), Torre_Velha_C (TVC), Torre_Velha_BA, Outeiro_Alto_BA (OABA), and Monte_da_Cabida_BA. The y-axis shows the *f4*-statistic values, with results displayed as the mean ± 1-SD. Markers represent the genetic sex, colors indicate Y-chromosome haplogroup lineages, and the black strokes denote Z-scores > 3. mtDNA haplogroups are indicated next to the dots. (C) Ancestry proportions (y-axis) for the aforementioned archaeological sites based on different admixture modeling frameworks (x-axis). The p-values are provided inside each column.

Under the *qpAdm* model using Anatolia_N and WHG instead of Anatolia_N and Iberian_HG, the inclusion of Goyet_Q2 was required for both Cova_das_Lapas_C and Torre_Velha_C, as observed in Neolithic individuals. However, for Cova_das_Lapas_C we found the best model fit when both WHG and Iberian_HG were included (Fig.2E; Table S8). Interestingly, *f4*- statistics revealed southern Iberian individuals possess an increased affinity towards Goyet_Q2 compared to WHG (Fig.2D). While outgroup *f3*-statistics suggest that Cova_das_Lapas_C shares both more genetic drift with Goyet_Q2 (Fig.S6D) and less drift with WHG (Fig.S6C; Fig.S6E) than Torre_Velha_C does, *qpAdm* modeling indicates a higher Goyet_Q2 component in the latter (Fig.2E; Table S8). This pattern is consistent with a north-south geographical ancestry gradient of increasing Goyet_Q2 affinity – although ancestry proportions and model fit in *qpAdm* when calculated from single individuals can be highly variable and should thus be interpreted with caution.

### Bronze Age

A total of twelve ancient individuals (genetically identified as 7 males and 5 females) were newly sequenced from three southern Portuguese Bronze Age archaeological sites, including Monte_da_Cabida_BA (n=5), Torre_Velha_BA (n=6) and Outeiro_Alto_BA (n=1). Of note, two individuals from Monte_da_Cabida_BA were previously published by Olalde et al 2019, but in the present study new sequencing data using the Twist Bioscience “Twist Ancient DNA” reagent kit were generated (Fig.S7; SI Text) (60).

First-degree relationships were identified both in Torre_Velha_BA (Fig.S8) and Monte_da_Cabida_BA (Fig.S7). In the former case, a male-female pair shared the mtDNA haplogroup X2c2 (Table S5) and could thus represent cases of mother-son or sibling kinship relationships, whereas in the latter case, a male-female pair carrying different mtDNA haplogroups—J1c2g and J1c12 (Table S5)—likely represented a father-daughter relationship. Interestingly, both individuals cluster separately from the rest of the population in the MDS plot based on 1- *f3*(Mbuti; Ind1, Ind2) (Fig.S9-10). Furthermore, this analysis showed that the Torre_Velha individuals from the Chalcolithic and Bronze Age cluster together.

Whole-genome analysis demonstrated that the Torre_Velha_BA, Outeiro_Alto_BA, and Monte_da_Cabida_BA archaeological sites closely align and cluster together with previously published Bronze Age Iberian sites (Fig.S11). We observe a Steppe-related ancestry presence across the three studied sites (Fig.S4) coinciding with a shift from the previous Iberian individuals in the PCA (Fig.3A) and a turnover in Y-chromosome R1b lineages (Fig.3B; Table S6). However, we also observe a persistence of mtDNA haplogroups associated with the LGM Iberian refugium (H1, H1+152 and U5b1+16189@16192) and the spread of Neolithic farmers into Europe from Southwestern Asia (X2b+226, X2c2, K1a, J1c1, J1c1f, J1c12, and J1c2g), but none of the most frequent mtDNA haplogroups of Eastern Europe (Table S5). Interestingly, three individuals from Torre_Velha_BA carried the same mitochondrial sub-haplogroup, J2b1a, which is relatively rare in Iberia—found only seven times before or during the Bronze Age (23,30,32)—and most frequent in present-day Southeastern Europe and Southwestern Asia (e.g., Greece, Turkey, or the Caucasus) (Table S5).

The Steppe-related ancestry is higher in the newly analyzed individuals from Torre_Velha_BA and Monte_da_Cabida_BA compared to individuals from the same archaeological sites previously published (23,31) (Fig.3B). While this Steppe-related ancestry is consistent across the Iberian Peninsula, as evidenced by ADMIXTURE analyses (Fig.S4), we found a slight decrease towards the south and southwest of Iberia (Fig.S12). Moreover, compared to broader Eurasian populations, the Iberian cluster exhibits lower Steppe-related ancestry overall, with individuals from Portugal among those with the lowest Steppe-related ancestry (Fig.S13). These results suggest a diffusion route of Steppe-related ancestry across Iberia, with the south and southwest of Peninsula corresponding to the latest point of arrival.

The modeling of a Distal framework with WHG and Anatolia_N as sources, including Russia_Samara_EBA_Yamnaya, improves the fit for Outeiro_Alto and is required as a third component to explain the genetic ancestry of Monte_da_Cabida_BA. However, this model does not fit Torre_Velha_BA (Fig. 3B; Table S9).

While semiproximal frameworks that include Iberia_C along with Russia_Samara_EBA_Yamnaya (Semiproximal_1) or Germany_BellBeaker (Semiproximal_2) can explain the genetic ancestry of Outeiro_Alto_BA and Monte_da_Cabida_BA, these models still fail for Torre_Velha_BA (Fig. 3B; Table S10). Instead, Torre_Velha_BA is best fit by a three-source model incorporating Iberia_C, Russia_Samara_EBA_Yamnaya, and Italy_C in Semiproximal_1 (Fig. 3B; Table S10). Alternatively, in Semiproximal_2, Torre_Velha_BA is best explained by a three-source model incorporating Iberia_C, Germany_BellBeaker, and several ancient Central Mediterranean Punic populations, along with ancient and present-day North African populations (Fig. 3B; Table S10). Despite these observations, we found no evidence of excess African ancestry in these populations compared to other Iberian regions (Fig. S14), suggesting either an equal distribution of African ancestry across the Peninsula or the persistence of deep genetic variation. Moreover, Central Mediterranean and North African Punic populations post-date Torre_Velha_BA, and this pattern was not confirmed by *f4*-statistics (Fig. S15C-F). In fact, Torre_Velha_BA shows higher affinities with Central Mediterranean islands, as indicated by outgroup *f3*-statistics (Fig. S15A-B). Therefore, the best-fit model in a Semiproximal_2 framework includes Iberia_C, Germany_BellBeaker, and Iran_N (Fig. 3B; Table S10).

The modeling of a Proximal framework provided a means to decipher the deep genetic variation found in Torre_Velha_BA, as the best-fitting model included Iberia_C_oSteppe and Italy_C ancestries which were also required in Monte_da_Cabida_BA, while Outeiro_Alto_BA was explained solely by Iberia_C_oSteppe (Fig. 3B; Table S11).

An outlier (PT_23212) in the Torre_Velha_BA group was identified in PCA (Fig. 3A), MDS plots based on 1- *f3*(Mbuti; Ind1, Ind2) (Fig. S8), and *f4*-statistics (Fig. 3C), suggesting a distinct genetic background. Despite this, no genetic ancestry from outside Iberia was detected (Fig. S16A-B, D-G), being the best-fitting model in *qpAdm* Iberia_C and Iran_C (Fig. 3B; Fig. S16C; Table S10).

### Roman Period

Since no new samples from the Iron Age period (Fig.S17) were available for study due to the widespread practice of cremation during that time (61), we next analyzed six unrelated individuals (genetically assigned as 1 male and 5 females) from the historical village of Idanha_a_Velha_Roman (central-eastern Portugal) (Fig.S18; Table S2). Among these, PT_24182 and ID_25538 exhibit significant North African ancestry as suggested in the PCA (Fig.4A; Fig.S19), which is further supported by positive *f4*-statistics values (Z > 2) of the form *f4*(PT_24182/ID_25538, Iberia_Roman_oLocal; Test, Chimp), where *Test* includes present-day African populations as proxies (Fig.S20E-F). These individuals appear to be admixed, with all *qpAdm* models requiring both Iberian (Spain_IA or Iberian_Roman_oLocal) and African (Saharawi or Morocco_Iberomaurusian) ancestries as proxies (Fig.4C; Table S12). These results are mirrored by observations of uniparentally-inherited markers, which showed both individuals carried mtDNA haplogroups most frequently found in present-day North Africans (X1c and T1a6)—but probably commonly found across Bronze Age and Iron Age Mediterranean—, while PT_24182 had a local R1b1a1b1a1a (R1b-P312) Y-chromosome haplogroup (Table S5-6).

**Figure 4.**
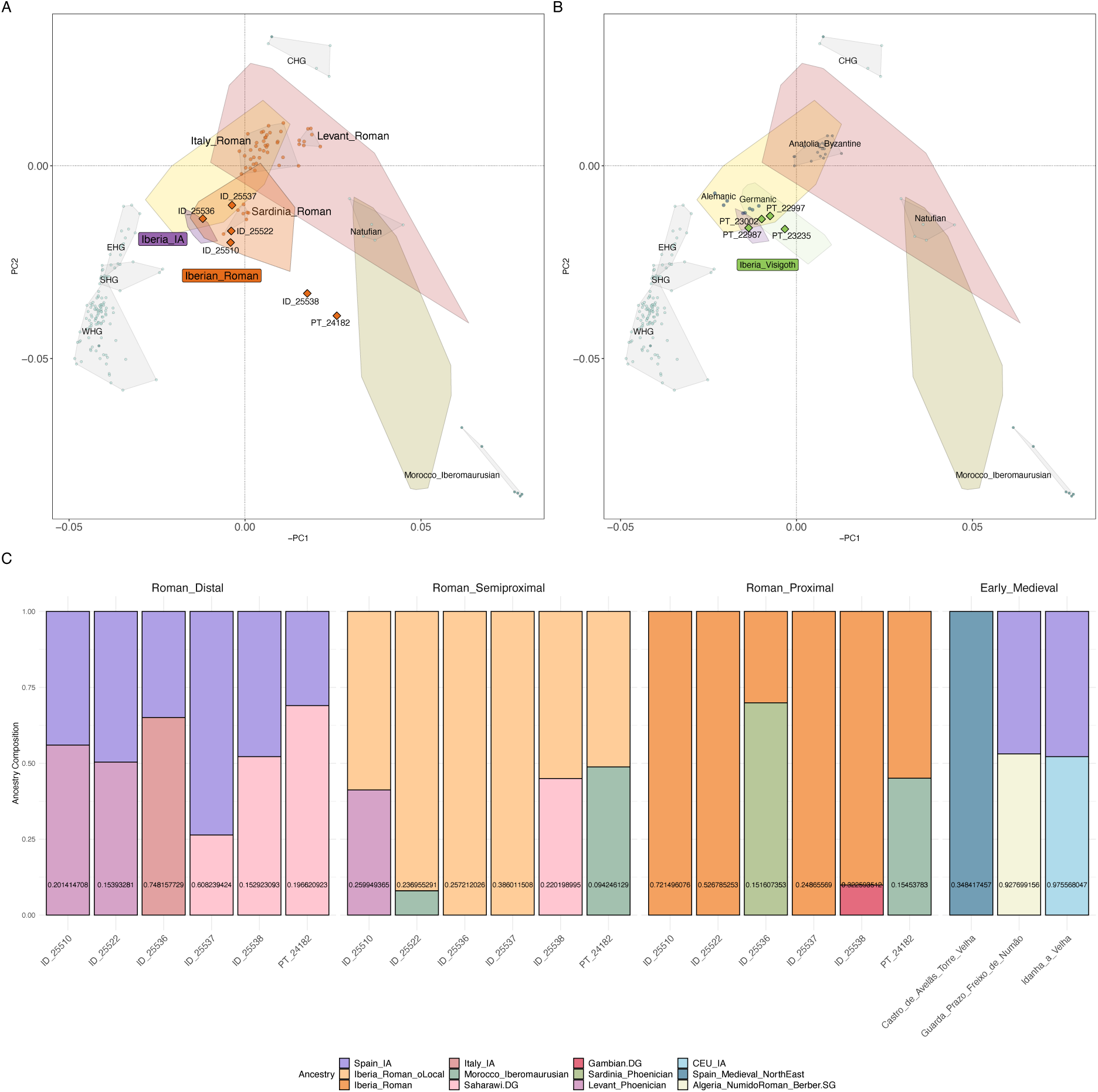
PCA of present-day West Eurasians and North Africans (overlaid colored polygons represent geographical clusters). The ancient individuals from Iberia and other regions were projected onto the first two principal components. Individuals from this study are represented as colored diamonds. Colors indicate different temporal periods as shown in 1B, with a focus on (A) the Roman and (B) Early Medieval periods. (C) Ancestry proportions using different admixture modeling frameworks for Idanha_a_Velha_Roman, and (D) Early Medieval archaeological sites.

Individual ID_25536 carried the local H1e1c mtDNA haplogroup and clusters with individuals carrying local Iberian ancestry from the Iron Age and Roman periods (Fig.4A). However, this individual shows a high shared genetic drift with Mediterranean populations (Fig.S20C), with the best fitting model in *qpAdm* requiring Spain_IA and Italy_IA as ancestry sources (Fig.4C; Table S12).

The remaining three female individuals, ID_25510, ID_25522, and ID_25537, cluster together with other Iberian individuals from the Roman period (Fig.4A), but also showed admixed ancestry. While the best fitting model for ID_25537, carrying the local Iberian mtDNA haplogroup K1b1a2, is a mixed genetic ancestry consisting of Spain_IA and North African, ID_25510 and ID_25522 are best modelled as local individuals with genetic admixture from the Eastern and African Mediterranean (Fig.4C; Table S12), respectively. These results align with their mtDNA haplogroups (H1ah1 and M1a3a, respectively) and the results from *f4*- statistics of the form *f4*(ID_25510/ID_25522, Iberia_Roman_oLocal; Test, Chimp), where *Test* includes Eurasian and African populations from the Roman period or their proxies (Fig. S20A- B).

### Early Medieval Period

We sequenced four unrelated ancient individuals (genetically identified as 1 male and 3 females) (Table S2; Fig.S21) from Idanha_a_Velha_EarlyMedieval (n=1), Guarda_Prazo_Freixo_de_Numão_EarlyMedieval (central Portugal; n=1), and Castro_de_Avelãs_Torre_Velha_EarlyMedieval (northeastern Portugal; n=2), all dating to the Early Medieval period, a time marked by the collapse of the Roman Empire and the recorded migrations of Visigoth and Suebi peoples from Central Europe. Their genetic profiles clustered closely with those of other Early Medieval individuals from the Iberian Peninsula in the PCA (Fig.4B).

Specifically, Idanha_a_Velha_EarlyMedieval (PT_22987) showed a clear genetic affinity with CEU_IA and German_Saxon populations as computed with outgroup *f3*-statistics of the form *f3*(Idanha_a_Velha_EarlyMedieval, Test; Chimp) and *f4*(Idanha_a_Velha_EarlyMedieval, X; Test, Mbuti) with *Test* including Eurasian and African populations from the Early Medieval period or close proxies, and *X* representing different Iberian and Central European Iron Age, Roman and Early Medieval populations (Fig.S22A; FigS23). While this individual carried the local mtDNA haplogroup T2b3 (Table S5), we were able to obtain the best fitting models of their ancestry as Spain_IA and CEU_IA or Iberia_Roman_oLocal and Alemanic (suitable proxy for the observed genetic influences) (Fig.4D; Table S13).

Guarda_Prazo_Freixo_de_Numão_EarlyMedieval (PT_23235) also showed an admixed genetic ancestry, which we were able to model as with Spain_IA and North African (Fig.4D; Table S13). Even though f-statistics computed as previously described in Idanha_a_Velha_EarlyMedieval indicate no significant gene flow from other populations occurred (Fig.S22B; FigS24), this individual harbored a W5 mtDNA haplogroup that is mostly found in present-day Germany, the Benelux, the British Isles, Norway and Poland (Table S5).

Finally, two individuals (one male - PT_23002; and one female - PT_22997) from Castro_de_Avelãs_Torre_Velha_EarlyMedieval revealed a genetic ancestry closely related to Spain_Medieval_NorthEast (a more recent proxy) (Fig.4D; Table S13). This suggests a local ancestry for this individual, despite f-statistics computed, as previously described for Idanha_a_Velha_EarlyMedieval, indicated possible gene flow from Eastern Mediterranean populations (Fig.S22C; FigS25), as previously found in Roman and Early Medieval Iberians (23,57). PT_22997 carried a local mtDNA haplogroup H1, while PT_23002 carried a local R1b1a1b1a1a (R1b-P312) Y-chromosome sub-haplogroup and an extremely rare mtDNA haplogroup (H44b) not previously found in ancient populations (Table S5-6), and only found in present-day people from the Western Black Sea.

### Islamic and Christian Conquest periods

We analyzed a total of 9 individuals from the Islamic Period (5 genetic males and 4 genetic females) (Table S2), including Santarém_Islamic (central Portugal; n=3) and two archaeological sites from Loulé_Islamic (southern Portugal), respectively Quinta_da_Boavista_Islamic (n=4) and Hospital_da_Misericórdia_Islamic (n=2). We observed that these individuals, along with previously published Islamic individuals, form a distinct cluster in the PCA, shifting towards North Africans when compared to other contemporary Iberian individuals (Fig.5A). Particularly, one outlier from Hospital_da_Misericórdia_Islamic (PT_24179) shifts towards present-day Sub-Saharan populations (Fig.S26).

**Figure 5.**
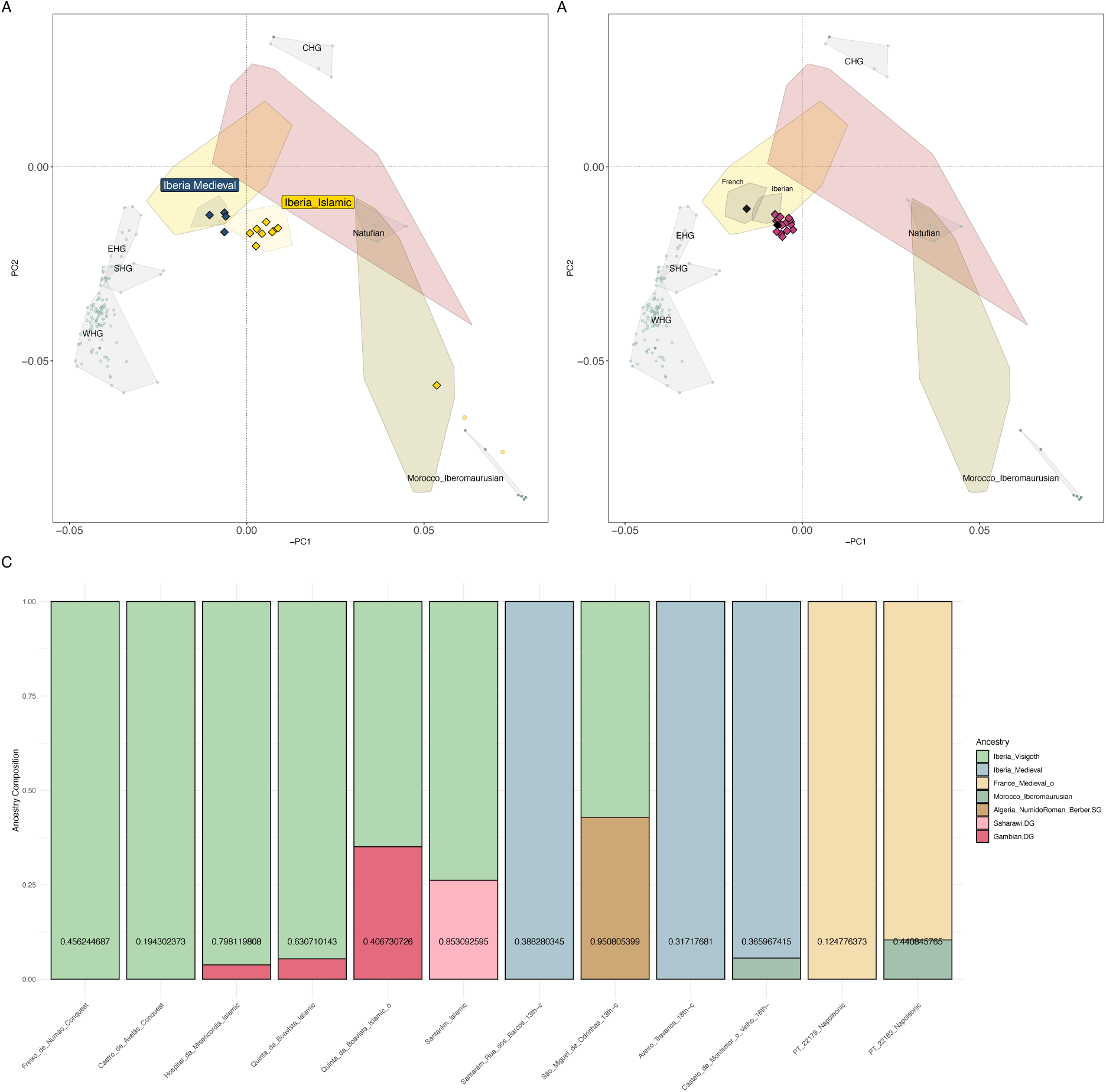
PCA of present-day West Eurasians and North Africans (overlaid colored polygons represent geographical clusters) with ancient individuals from Iberia and other regions projected onto the first two principal components. Individuals from this study are represented as colored diamonds. Colors indicate different temporal periods as shown in 1B, with a focus on the (A) Islamic (yellow) and Christian Conquest (dark blue) periods, and (B) 13^th^-18^th^ century (pink) and 19^th^ century (black) Portugal. (C) Ancestry proportions for Portuguese individuals from the Islamic and Christian Medieval period, 13th century, 18^th^ century, and 19^th^ century.

To determine which archaeological site exhibited the highest levels of genetic drift with African populations, we computed outgroup *f3*-statistics of the form *f3*(Test, Mbuti; Chimp), where *Test* rotates through the Portuguese Islamic archaeological sites. We observed that Santarém_Islamic showed the highest shared genetic drift with Mbuti, followed by Quinta_da_Boavista_Islamic_o (PT_24179), Hospital_da_Misericórdia_Islamic, and lastly, Quinta_da_Boavista_Islamic (Fig.S27). African gene flow is also evident in *qpAdm* results (Fig.5C; Table S14) and *f4-*statistics of the form *f4*(Santarém_Islamic/Loulé_Islamic, Spain_IA/Iberia_Visigoth; Test, Chimp), with *Test* including Eurasian and African populations from the Islamic period or close proxies (Fig.S28-29). Admixture modeling using Gambian as proxy indicates that West African ancestry is present across Hospital_da_Misericórdia_Islamic, Quinta_da_Boavista_Islamic, and its outlier individual (Fig.5C; Table S14). While the first two sites exhibit low levels of West African ancestry, the outlier individual shows approximately 40% (Fig.5C; Table S14). Meanwhile, Santarém_Islamic displays significant North African ancestry (∼26%), using Saharawi as a proxy (Fig.5C; Table S15).

The analysis of uniparental markers further supports these findings, with widespread presence of mtDNA and Y-chromosome haplogroups commonly found in North Africa (Table S5-6). We note that the uniparentally-inherited markers of the individuals from Santarém_Islamic and Loulé_Islamic were previously reported (62) (AF Maurer, personal communication, 2024). However, our newly sequenced data for Santarém_Islamic offers improved haplogroup resolution for both mtDNA and the Y-chromosome.

One individual from Santarém_Islamic (PT_24170) shows a ROH profile (> 20 cM) suggestive of consanguinity, likely being the offspring of a sibling pair (Fig. S30). However, none of the other analyzed Islamic individuals were genetically related (Fig. S31).

We also analyzed a total of 4 samples (2 genetic males and 2 genetic females) (Table S2) from Guarda_Prazo_Freixo_de_Numão_Conquest (n=1) and Castro_de_Avelãs_Torre_Velha_Conquest (n=3), dating to the period of Christian Conquest. Notably, one individual from Castro_de_Avelãs_Torre_Velha_Conquest (PT_22994) was previously reported to carry an extra copy of the X-chromosome, consistent with Klinefelter’s Syndrome (Table S2) (63).

The Christian Conquest individuals formed a distinct cluster in the PCA compared to the Islamic individuals (Fig.5A). Additionally, *f4*-statistics of the form *f4*(X, Y; Test, Mbuti), where *X* represents Guarda_Prazo_Freixo_de_Numão_Conquest or Castro_de_Avelãs_Torre_Velha_Conquest, *Y* represents different Iberian Medieval populations, and *Test* rotates through various Eurasian and African populations from the Medieval period or close proxies, showed no significant gene flow from populations with different genetic ancestry (Figs.S32-33). Supporting this finding, both Portuguese Christian Conquest archaeological sites could be modeled as Iberian_Visigoth (Fig.5C; Table S15). All individuals carried local mtDNA (X2b, V, H4a1a, and H10f, the latter two being quite rare— Table S5) and Y-chromosome (R1b) haplogroups (Table S6), indicating a degree of genetic continuity from Early Medieval populations.

To test the genetic distances between the Early Medieval and Christian Conquest individuals— as some were sampled from the same archaeological sites—we computed an MDS plot based on 1- *f3*(Mbuti; Ind1, Ind2) (Fig.S34). We observed that all but one sample from Castro_de_Avelãs_Torre_Velha_Conquest (PT_23237) clustered closely together, indicating no major genetic differences between the two periods. Moreover, while Idanha_a_Velha_EarlyMedieval aligned closely with individuals from Castro_de_Avelãs_Torre_Velha across both time periods, we detected significant genetic differences between the two individuals from different periods at Guarda_Prazo_Freixo_de_Numão.

### Kingdom of Portugal

We analyzed a total of 18 ancient individuals dating from after the establishment of the Kingdom of Portugal in the middle of the 12^th^ century. These included 13^th^ century individuals from Santarém_Rua_dos_Barcos_13th-c (central Portugal; 6 genetic males and 3 genetic females) and São_Miguel_de_Odrinhas_13th-c (western Portugal; 2 genetic males and 2 genetic females); as well as 18^th^ century individuals from Castelo_de_Montemor_o_Velho_18th-c (central Portugal; 4 genetic females and 1 genetic male) and Aveiro_Travanca_18th-c (northwestern Portugal; 1 genetic male) (Table S2).

Our results indicate Portuguese populations from the 13^th^ to the 18^th^ centuries maintained genetic continuity with local inhabitants from the post-Islamic period (Fig.5B). However, *qpAdm* modelling required an additional proportion of African ancestry (Fig.5C; Table S16- 17). This is further supported by the analysis of uniparental markers, with the presence of mtDNA (U6d3a and U6a3b) and Y-chromosome (E1b1b1b1a1 and E1b1b1a1b1a10a1a3) haplogroups common in North Africa but present at low frequencies in present-day Iberian populations (Table S5-6). Additionally, we also found an individual (PT_23227 from Santarém_Rua_dos_Barcos_13th-c) who carried Q1b1a3a1, a Y-chromosome haplogroup commonly found among Ashkenazi Jews and in Central Asia, but which is rare in Iberia (Table S6).

Kinship analysis (Fig.S35-37) at São_Miguel_de_Odrinhas_13th-c revealed complex familial relationships among the sampled individuals. Specifically, one female (PT_24164) and one male (PT_24163) carrying the H1 mtDNA haplogroup shared a first-degree relationship, possibly as siblings or mother and son. The male individual also shares a third-degree relationship with another male (PT_24165) carrying a different mtDNA haplogroup (T2b3) but the same J2b1b1 Y-chromosome haplogroup. PT_24165, however, is not related to PT_24164, suggesting PT_24163 and PT_24165 could be paternal cousins and PT_24164 and PT_24163 could be mother and son.

### Napoleonic period

Finally, we analyzed two genetically male individuals (Table S2) buried at Castelo_de_Abrantes_19th-c (Santarém, central Portugal), dating from the occupation of Napoleon’s forces in Portugal. Interestingly, we found one individual (PT_22183) with a genetic affinity to present-day French (Fig.5B; Fig.S38) that could be modelled as French_Medieval in *qpAdm* (Fig.5C; Table S18). Likewise, the other individual (PT_22183) was of French_Medieval and African ancestry and carried a mtDNA haplogroup (L2b3a, Table S5) of West/Central Africa origin, also commonly found in Europe due to historical migrations. PT_22183 carries a Y-chromosome haplogroup I2a1b1a1b1a1a1a (Table S6) which is more commonly found in Eastern and Southeastern Europe.

## 3. Discussion

The genetic profile of Cova_das_Lapas_N aligns with previous research on the Iberian Peninsula, indicating admixture between local hunter-gatherers and Anatolian farmers (23,24,31,49). This conclusion is further supported by analysis of uniparentally-inherited markers, which reveals a persistence of mtDNA haplogroups from Upper Paleolithic and Mesolithic Iberia linked to LGM refugia and postglacial re-expansions, as well as haplogroups associated with Early Neolithic migrations from Southwestern Asia (64,65).

Our study identified the Y-chromosome haplogroup I2a exclusively among the male individuals from Cova_das_Lapas_N. This lineage is connected to Central European hunter-gatherers who mixed with East Mediterranean Neolithic farmers associated with the westward expansion of the Cardial culture (49). These farmers carried Y-chromosome haplogroup G2a (absent in Cova_das_Lapas_N) and mtDNA haplogroup K1a (present in Cova_das_Lapas_N). Although this admixture occurred around 2,000 years earlier in Central Europe, the genetic traces in Cova_das_Lapas_N reflect a broader pattern of genetic diffusion over time. This suggests that the admixture likely happened earlier and persisted locally, as the site dates back to the Middle/Late Neolithic, postdating the Cardial culture period in Portugal (5,500-5,000 BCE).

Furthermore, the presence of two Y-chromosome haplogroups from the same clade, alongside high mtDNA diversity and exclusively paternal first- and second-degree relationships, suggests a pattern of patrilocality. This is consistent with findings from other Neolithic European archaeological sites (66–68). Additionally, radiocarbon dates for Cova_das_Lapas_N, combined with votive artifacts and preliminary anthropological analyses, point to a complex management of the burial site, reinforcing the notion that cultural exchange, rather than mere admixture, shaped these Neolithic communities. These interactions likely involved the exchange of goods, information, acculturation, and female exogamy, which could explain patrilocality and the continued presence of hunter-gatherer mtDNA in later Neolithic populations.

The persistence of Upper Paleolithic Magdalenian ancestry in Cova_das_Lapas_N, compared to continental Europe and Northern and Northeastern Iberia, underscores the Iberian Peninsula’s role as a refugium for Upper Paleolithic populations. The relative isolation of the western Atlantic coast in response to environmental changes likely contributed to the distinct genetic profile observed in this region. However, further sampling of Upper Paleolithic, Mesolithic, and Neolithic Iberian populations is necessary to fully discern the genetic landscape of Iberian hunter-gatherers over time, particularly as these populations likely remained in relative isolation and contained heterogeneous genetic profiles that were carried through time.

We found similar patterns of population persistence in Chalcolithic Portugal, with genetic analyses of Cova_das_Lapas_C and Torre_Velha_C revealing notable regional genetic continuity. Importantly, Cova_das_Lapas_C clusters with the earlier Cova_das_Lapas_N population, sharing the paternal Y-chromosome lineage I2a1a1a1a and the local U5b mtDNA sub-haplogroup. The admixture proportions between hunter-gatherer and Neolithic-related ancestry are comparable to those observed during the Neolithic, including contributions from local hunter-gatherers and Anatolia_N. Additionally, we detected a significant Upper Paleolithic Magdalenian ancestry component that increases southwards across the peninsula but dissipates over time. This pattern may suggest: (i) a more pronounced population isolation at the southern and southwestern edges of the Iberian Peninsula, with loss over time due to genetic drift; (ii) dilution of this ancestry over time with the arrival of other Neolithic groups; or (iii) limited resolution resulting from insufficient sampling.

The genetic analysis of the Bronze Age archaeological sites of Torre_Velha_BA, Monte_da_Cabida_BA and Outeiro_Alto_BA also demonstrates strong regional ties and continuity, as evidenced by the persistence of local mtDNA haplogroups. However, this is accompanied by a notable presence of Steppe-related ancestry associated with migrations from the North Pontic Steppe into Central Europe that eventually spread into the Iberian Peninsula, which is further supported by the arrival and replacement of distinct R1b Y-chromosome lineages (23,27,31,33,69). Interestingly, we observe a nuanced decline in Steppe-related ancestry towards the south and southwest of the Iberian Peninsula, indicating that this migration had a relatively lower impact in Portugal compared to other regions of the peninsula. This diminishing trend in Steppe-related ancestry forms a gradient that likely reflects dilution through admixture with local Chalcolithic populations as the migrants moved southwestward from the Northeast of the Peninsula.

Interestingly, the new ancient genome-wide data also revealed connections to the Central Mediterranean during the Bronze Age, which had only been sporadically reported in Southern Spain (32). These findings indicate the possibility of early contacts between western Iberia and Mediterranean cultures, potentially linked to maritime trade and cultural exchanges that peaked during the Iron Age but predate the foundation of the Phoenician city of Cádiz(37). In fact, this genetic signal could be representing the earliest signatures of contacts with Phoenicians, which followed the African coast of the Mediterranean and later influenced southwestern Iberia(37). These contacts could have also played a role in the development of the Tartessian culture in southwestern Iberia, reflecting a broader pattern of cultural and genetic exchange across the Mediterranean and further west to the Atlantic coast that occurred as early as the Bronze Age.

Due to the scarcity of archaeological remains from the Portuguese Iron Age, which is largely attributed to prevalent funerary cremation practices (61), we were unable to analyze samples from this period. However, our analysis of previously published Spanish populations reveals a genetic continuity through the Iron Age, with notable connections extending to the Eastern Mediterranean, a pattern that was previously suggested (23).

The genetic analysis of individuals from the Roman period sampled at the historical village of Idanha_a_Velha_Roman revealed significant diversity, supporting the hypothesis that the town functioned as a key crossroads within Lusitania, linking the capital cities Emerita Augusta, Bracara Augusta, and Lucus Augusti (70). The presence of distinct African and Mediterranean ancestries highlights the impact of extensive trade, migratory networks, and cultural exchanges characteristic of Roman Iberia (71). Migrants that often were Roman citizens and part of the social elite were buried in mausoleums of Italic tradition, as shown by architectural elements found in the area (72). Conversely, the presence of enslaved individuals and freedmen, some with Greek names and potentially of Eastern Mediterranean or North African origin, reflects a more complex social dynamic with varying levels of integration (72). Epigraphic evidence also points to military personnel involved in public works and gold mining, contributing to the population’s diversity (72). Hence, African ancestry may be linked to trade, as suggested by the discovery of amphorae and fine tableware from Roman Africa, reflecting the city’s role in broader Mediterranean trade networks. Additionally, the Latin name *Niger*, found in local inscriptions, may refer to dark-skinned individuals, offering further insight into the town’s diverse population (72).

Recent paleogenomic studies have documented a genetic shift towards Near Eastern and Anatolian genetic influences in Imperial Roman populations compared to earlier Iron Age populations in Italy, the Southern Arch and the Balkans (29,57,73,74). Some individuals from Idanha_a_Velha_Roman show a similar shift, though others exhibit a notable shift towards North Africa. While direct comparisons with Iron Age samples are lacking, this shift suggests a pattern akin to that observed in Italy (74), though perhaps to a lesser extent. This lower affinity may indicate a degree of higher genetic continuity from the Iron Age populations within the Roman period in Iberia compared to Italy.

In the aftermath of the fall of the Western Roman Empire, historical records document the arrival of Germanic tribes from Eastern/Central Europe, in particular Suebi and Visigoth peoples, to the two provinces overlapping with contemporary Portuguese territory: Gallaecia and Lusitania (41,42). Our analysis reveals an ancient female individual from Idanha_a_Velha_Roman with clear genetic affinities to Central European populations, an observation that is consistent with a genetic link to migrant Suebi- or Visigoth-related communities from Eastern and Central Europe.

In contrast, the individual from Guarda_Prazo_Freixo_de_Numão_EarlyMedieval had a predominant Iberian ancestry and North African influences, along with maternal Central European ties. Similarly, the two individuals from Castro_de_Avelãs_Torre_Velha_EarlyMedieval were also local but displayed Eastern Mediterranean influences that are likely associated with population connections lasting from the Roman period. These individuals also share a small proportion of Central European ancestry, suggesting possible admixture some generations further in the past, or instead recent intermingling with admixed individuals. Overall, our study highlights complex population dynamics in the Iberian Peninsula during the Early Medieval period that can only be resolved through further sampling and analyses.

The genetic analysis of individuals from the Islamic and Christian Conquest Medieval contexts at various sites across Portugal reveals a complex and nuanced picture of population dynamics during this historical time. The individuals from Islamic contexts exhibit a more diverse genetic landscape, characterized by an evident admixture with Sub-Saharan and North African populations. The presence of African ancestry, particularly among individuals from sites like Santarém_Islamic (southern Portugal), is supported by both autosomal DNA analyses and uniparental markers (62). The admixture modeling suggests that while some individuals carried only low levels of African ancestry, others displayed substantial admixture, indicating a complex and heterogeneous population during the Islamic period in Portugal, being more pronounced in the southern regions, probably due to the extended time of Islamic presence in Southern Iberia.

While cousin marriages were common in the Islamic world to preserve family wealth and strengthen alliances (75), the ROH profile of PT_24170 from Santarém_Islamic indicates that this individual was likely the offspring of siblings—the first such case identified in ancient Iberia. Although Islamic law explicitly prohibits sibling unions (Quran 4:23), this finding may point to rare deviations from religious norms, potentially influenced by local practices or limited social and geographic marriage pools. Alternatively, it is possible that this individual’s parents were unaware they were siblings, or that this individual could be the result of multiple generations of cousin marriages.

In contrast, the genetic continuity observed among individuals from the Christian Conquest reflects a strong preservation of local ancestry with minimal external gene flow. Specifically, individuals from Castro_de_Avelãs_Torre_Velha_Conquest (northern Portugal) exhibit limited influence from the brief Islamic expansion into the region. Situated strategically near the Douro River, Castro_de_Avelãs_Torre_Velha_Conquest was crucial during the Christian Conquest, as this area, previously described as the “Desierto del Duero” by Sánchez-Albornoz, was contested and sparsely populated and was central to both Christian and Islamic military efforts. Despite the brief Islamic control under Almazor’s campaigns at the end of the Caliphate of Córdoba (47), our findings reveal local genetic ancestry persisted.

The genetic analysis of Portuguese populations from the 13^th^ to 18^th^ centuries reveals a continuity with the post-Islamic period, highlighting the persistence of local genetic signatures over several centuries. However, the small yet notable proportion of African ancestry detected in admixture modelling, further supported by North African mtDNA and Y-chromosome haplogroups, underscores the lasting genetic impact of the earlier Islamic presence in the region. This suggests that the genetic influence of Islamic populations persisted long after their political power declined. Alternatively, due to frequent mobility between Europe and North Africa during the Medieval and Modern periods (47), the detected African ancestry may instead reflect the contributions of subsequent waves of African migrants.

One particularly intriguing finding is the identification of the Y-chromosome haplogroup Q1b1a3a1 in an individual from Santarém_Rua_dos_Barcos_13th-c (PT_23227). This haplogroup is rare in Iberia but more commonly associated with Ashkenazi Jewish populations and Central Asia. During the 13^th^ century, Jewish communities in Portugal were well integrated, contributing significantly to the economy, culture, and intellectual life. Throughout the Medieval period, they maintained their autonomy and identity while coexisting peacefully with the broader society (47). However, this harmony began to erode in the late 15^th^ century, as increasing socioeconomic and religious restrictions, including the Expulsion Decree of 1496, led many Jews to convert to Catholicism and become known as New Christians (47). While this led to the migration of thousands of Sephardic Jewish populations to other parts of Europe and the New World, the genetic signature of these populations persists in present-day people from Portugal (76–79).

The genetic analysis of two soldiers from the Napoleonic forces sampled at Castelo_de_Abrantes_19th-c provided insights into the origins of some of the French troops that occupied Portugal. The first individual, who exhibits a clear genetic affinity with present-day French populations, can be modeled as French_Medieval. This result is consistent with the historical context, as Napoleon’s occupation force primarily consisted of soldiers from France(80). The second individual (PT_22183) presents a more complex genetic background, requiring both French_Medieval and an African proxy in admixture modelling. This individual carries a mtDNA haplogroup L2b3a, which is of African origin, particularly prevalent in West and Central Africa, and likely introduced in Europe through historical migrations, including the transatlantic slave trade and earlier movements across the Mediterranean. The presence of this haplogroup in a Napoleonic soldier could suggest the inclusion of soldiers of African descent in the ranks, either through direct recruitment or as descendants of earlier African immigrants to Europe. Additionally, PT_22183’s Y-chromosome haplogroup I2a1b1a1b1a1a1a is more commonly found in Eastern and Southeastern Europe. This haplogroup’s presence indicates possible migration or conscription of individuals from these regions into the Napoleonic forces. Napoleon’s armies were known for their diverse composition, often incorporating soldiers from various parts of Europe, including those from territories under French influence or control (80). This individual reflects the broader socio-political and demographic currents of the time, including colonialism, migration, and the movement of peoples across Europe and Africa. This genetic diversity within the invading force underscores the extensive reach of Napoleon’s military campaigns and the varied backgrounds of those who served within his armies.

## 4. Conclusions

In this study, we reconstructed the genetic history of Portugal from the Neolithic to the contemporary period, revealing a complex and dynamic population history.

During the Neolithic, paleogenomic evidence indicates a potential patrilocal society with admixture between local hunter-gatherer and Anatolian farmers, marked by the persistence of Upper Paleolithic Magdalenian ancestry. This genetic signature extended through the Chalcolithic, while the Bronze Age revealed influences from the Steppe and the Central Mediterranean Sea. Importantly, the latter might represent genetic evidence associated with cultural exchanges and maritime trade predating the arrival of Phoenicians in the Iberian Peninsula.

The genetic analysis of the Roman period highlighted the important role of migration, and the strengthening of trade networks as evidenced by a diverse genetic pool, with influences from Europe, Africa and the Mediterranean. The Early Medieval period further enriches this perspective, with connections to Central Europe probably associated with Suebi and Visigoth populations complementing earlier North African and Mediterranean ancestries already found in the local population.

During the periods of Islamic and Christian Conquest, northern Portugal exhibits strong genetic continuity to the Early Medieval period, while the south reveals significant African admixture. These observations are in agreement with historical records of greater cultural and political influences of Islam in the South of the country. The enduring presence of Jewish and Islamic genetic markers into the post-Medieval period reflects the lasting impact of these cultures on the genetic pool of the Portuguese.

Overall, our study highlights Portugal’s intricate genetic heritage, shaped by centuries of migration, cultural exchange and interaction, with significant contributions from diverse populations shaping the contemporary genetic diversity of the country.

## 5. Methods

### 5.1. Experimental design

#### Archaeological samples

A total of 94 individuals from Portugal dated to a time span from ∼3000 BCE to ∼1800 CE were analyzed for the presence of ancient DNA. A detailed description of the archaeological context, sites, and individuals is reported in Supplementary Information (SI) and summarized in Table S1.

#### Laboratory Facilities

Pre-amplification experiments took place at the ultra-clean laboratory facilities of the Australian Centre for Ancient DNA (ACAD). Stringent laboratory procedures were implemented to minimize contamination and uphold the genetic data’s high-quality standards (81,82). All post-amplification experiments were conducted in standard molecular biology laboratories at the University of Adelaide, followed by subsequent bioinformatics workflows executed on the University of Adelaide’s HPC.

#### DNA Extraction and Library Preparation

We preferentially sampled petrous bones and teeth. Details of sampled skeletal elements are listed in Table S1. Before DNA extraction, the skeletal samples underwent sterilization using UV, bleach, ethanol, and/or scraping the bone surface to minimize contamination. Approximately 0.1 g of bone powder was utilized for DNA extraction. A method optimized for degraded DNA (83) was employed to retrieve ancient DNA molecules, and subsequently, partially UDG-treated (84) double-indexed double-stranded DNA libraries were generated (85). Quality controls and DNA quantification were carried out using Qubit (Thermo Fisher) and TapeStation (Agilent). DNA-sequencing libraries were sent for sequencing on a NovaSeq 6000 platform at the Kinghorn Centre for Clinical Genomics (Sydney, NSW, Australia).

#### Data Processing

Raw data underwent processing using the aDNA analysis workflow package nf-core/eager version 2.4.6 (86). Merged read mates were aligned to the GRCh37d5 reference genome using *bwa aln* with parameters *-l 1024 -n 0.01 -o 2* (87). Trimming of two nt from the terminal ends of all retained reads was performed using the *trimBam* function of *bamUtil* (https://github.com/statgen/bamUtil). Standard quality filters (mapping quality ≥ q25 and base quality ≥ Q30) were applied, and reads were deduplicated using the *MarkDuplicates* function from Picard.

#### Authentication and Quality Control

DamageProfiler (88) was used to assess aDNA authenticity, endogenous DNA proportion, fragment size distribution, contamination estimates, and post-mortem damage rates at the read termini (Table S1).

#### Genetic Sex Determination

Genetic sex determination was performed using SexDetERRmine (https://github.com/nf-core/modules/tree/master/modules/nf-core/sexdeterrmine) with default quality cut-off values for -q30 and -Q30 (Table S2).

#### Library Enrichment

A total of 83 libraries met authenticity quality thresholds and underwent enrichment. Over-amplification was conducted to achieve the 1000 ng required for enrichment. For each library, the PCR reaction mix consisted of 5-10 µl of library, 25 µl of KAPA HiFi HotStart ReadyMix (Roche), 5µl each of 10 µM IS5 and IS6 primers (84), and ultrapure water in a total volume of 50 µl. PCR amplification was performed with an initial denaturation and polymerase activation at 98°C for 2 min, 15 cycles of 98°C for 20 sec, 56°C for 30 sec, 72°C for 45 sec, and a final extension at 72°C for 5 min. DNA purification was performed using 1.2x AmpureXP beads with two 80% ethanol washes. The DNA was eluted in 30 µl of water.

Pooling of libraries was determined from total DNA quantification of each library, considering the endogenous content, as described in (89). Enrichment was performed using the Twist Bioscience “Twist Ancient DNA” reagent, following the manufacturer’s protocol. The post-enrichment PCR amplification was performed using KAPA HiFi HotStart ReadyMix (Roche) and IS5 and IS6 primers as described above, with a 98°C initialisation for 24 sec, 7 cycles of 98°C for 15 sec, 60°C for 30 sec, 72°C for 30 sec, and a 72°C final extension for 60 sec. Some libraries underwent a reconditioning PCR to reduce heteroduplexes. Specifically, libraries were concentrated down to 5 µl and mixed with 10 µl of Herculase Buffer (Agilent), 5 µl of 2.5 nM dNTPs, 1 U of Herculase II Fusion (Agilent), 1 µl each of 10 µM IS5 and IS6 primers, and ultrapure water in a final volume of 50 µl. This solution was then reconditioned with one cycle of 95°C for 2 min, 58°C for 2 min, and 72°C for 5 min. DNA purification and library quality control and quantification were performed as described above. Enriched libraries were sent for sequencing on a NovaSeq 6000 platform at the Kinghorn Centre for Clinical Genomics (Sydney, NSW, Australia).

#### Genotyping

Sequencing raw data underwent processing as previously described (Table S3). Pseudohaploid variant calling was executed using the Twist Bioscience “Twist Ancient DNA” SNP panel (60) with *pileupCaller* (https://github.com/stschiff/sequenceTools).

As expected, the hybridization procedure ensured efficient enrichment for DNA targets, resulting in increased endogenous DNA proportions and a decrease in the complexity of the libraries (Table S3). After selecting individual samples that met the following criteria: (i) more than 15,000 SNPs covered by at least one read of the Twist SNPs panel; (ii) the presence of the misincorporation patterns characteristic of aDNA (>3%); and (iii) absence of contamination; a total of 68 samples were retained for genome-wide data analyses.

### 5.2. Uniparental markers

#### Mitochondrial DNA

The analysis of mitochondrial DNA involved merging the raw data obtained from two distinct sources: i) data generated using the Twist Bioscience “Twist Ancient DNA” reagent (60), and ii) data obtained with the myBaits Expert Human Affinities Prime Plus Kit by DAICEL Arbor Biosciences (Ann Harbor, MI, USA). These datasets were processed following the respective manufacturers’ protocols and utilizing the pooling technique outlined in (89). As previously validated (60,90), no allelic biases were present in the mitochondrial capture performed with the myBaits Expert Human Affinities Prime Plus Kit.

The raw data from both sources underwent processing using the aDNA analysis workflow package nf-core/eager version 2.4.6 (86). Merged reads were then mapped to the mitochondrial revised Cambridge Reference Sequence (rCRS) using CircularMapper (https://github.com/apeltzer/CircularMapper) and *bwa aln* with parameters *-l 1024 -n 0.01 -o 2 -k 2* (87). Read trimming and filtering followed the procedures outlined above (Table S4). The read pileups were visually inspected using Geneious v2022.1.1 (Biomatters; https://www.geneious.com) (Table S4). Mitochondrial haplogroup calling was performed using *haplocart* (91), and contrasted using mitoverse HaploCheck version 1.3.2 (92), which also enabled the estimation of contamination levels (Table S5).

#### Y-chromosome

The Y-chromosome genotype was determined using the *UnifiedGenotyper* tool from Picard v2.26.0 (Broad Institute, 2019), and inferred from the merged, trimmed BAM files obtained through the nf-core/eager pipeline. The analysis was conducted using the GRCh37d5 genome as a reference. Only the Y-chromosome SNPs identified from the Twist Bioscience “Twist Ancient DNA” SNP panel (60) were considered for analysis. The python script hGrpr2.py from *HaploGrouper* (https://gitlab.com/bio_anth_decode/haploGrouper) was used to obtain the Y-chromosome haplogroups with the default tree and the SNP file from the Ghr37 reference genome (Table S6).

### 5.3. Kinship estimation

BREADR was used to determine kinship between pairs of individuals from the same archaeological site using the default settings (93).

### 5.4. Population genetic analysis

#### Dataset

We merged our final dataset with previously published datasets of ancient and modern individuals reported by the Reich Lab (94)(https://reich.hms.harvard.edu/datasets; please see Table S19 for a detailed list of individuals and the new labels used) .

#### PCA

We computed PCA using the *smartpca* software from the EIGENSOFT package (v7.2.1) with the *lsqproject* and *SHRINKMODE* option *YES* and an extended list of modern and ancient populations from Eurasia, Africa and the Caucasus. The ancient individuals were projected onto PC1 and PC2.

#### F-statistics

If two individuals showed evidence for a kinship relation, we removed the one with the lowest number of SNPs for population genetics analyses, but not for archaeological-specific analyses. In addition, only individuals with >35,000 SNPs were retained. F-statistics were computed with ADMIXTOOLS v5.1 (https://github.com/DReichLab). For *f4*-statistics, we used qpDstat and the activated *f4*-mode. *f3*-statistics were calculated using qp3Pop; and *f4*-Ratios were calculated using D4RatioTest.

#### Admixture modelling

As described above, relatives with the lowest number of SNPs were discarded from the analysis, and only individuals with more than 35,000 SNPs were retained.

ADMIXTURE was used to define the main genetic cluster profiles. Data was pruned for linkage disequilibrium using PLINK with parameters *--indep-pairwise 50 10 0.1* and *--geno 0.999*. Four populations were selected as fixed source ancestry groups (k=4) implementing a supervised ADMIXTURE model with ten replicates.

To model the genetic ancestry of the newly sequenced ancient individuals, we used the *qpAdm* program from the ADMIXTOOLS v5.1 package (https://github.com/DReichLab), with the “*allsnps: YES*” option to maximize the number of SNPs used and subsequently increase the power to discriminate different models.

We also used *qpAdm* to quantify the proportion of genetic ancestry contributed by each source population (Table S19). The ancestry proportions in the target population were inferred on the basis of how the target population is differentially related to a set of reference/outgroups via the source populations.

For all the models applied here, we have used a set of 12 outgroups in total (Mbuti.DG, Ethiopia_4500BP.DG, Papuan.DG, Belgium_UP_GoyetQ116_1, Czech_Vestonice, Italy_North_Villabruna_HG, Russia_MA1_HG.SG, Russia_Ust_Ishim_HG.DG, Malalmuerzo).

### 5.5. Runs of homozygosity

We used hapROH VERSION 0.64 with default settings to identify runs of homozygosity within the genome (95). Only individuals with > 400,000 SNPs covered in the panel were used in this analysis.

## 6. Declarations

### 6.1. Ethics approval and consent to participate

Not applicable.

### 6.2. Consent for publication

Not applicable.

### 6.3. Availability of data and materials

The datasets generated and analyzed during the current study will become publicly available upon publication.

### 6.6. Authors’ contributions

Conceptualization: X.R-R., B.L., J.C.T.

Sample provision: R.A.M., L.M., E.C., S.T., V.M., T.M.F., A-F.M., A.M.S., P.C.

Lab work: X.R-R., M.P.W., E.C., C.H., L.T., J.C.T., B.L.

Data processing: X.R-R., S.R.

Data analysis: X.R-R., R.D., S.R., D.R.C-A.

Visualization: X.R-R., R.D. Interpretation: All authors Supervision: B.L., J.C.T. Funding: R.T., B.L., P.C., J.C.T.

Writing - original draft: X.R-R., J.C.T. Writing - review & editing: All authors

## Acknowledgements

Laboratory work was conducted at The University of Adelaide with support from technical officers Holly Heininger, Vilma Pérez, Corinne Mensforth, and Navdeep Kaur. Computational analyses were conducted using supercomputing resources provided by the Phoenix HPC service at the University of Adelaide.

