## Supplementary Information for "The genetic history of Portugal over the past 5,000 years"

### 1. Archaeological context

#### 1.1. Cova das Lapas (Middle Neolithic)

Cova das Lapas is a very small cave located in the Limestone Massif of Estremadura that was intensively used as a sepulchral space. The sequence of absolute dating and votive artifacts indicates that this necropolis was used in a short period of time, between 3245-3263 and 3036-2913 BCE, with only one exception, a cranium dated to the Chalcolithic. The detailed anthropological study is ongoing, but it was already possible to identify complex sequences of management of the sepulchral space. The votive artifacts include the characteristic elements of the magic-religious complex of the first and second phases of Megalithism: geometric armatures, blades, daggers, scarce ceramics and one engraved schist plaque.

The previously published individual (I5428) from this archaeological site by Olalde et al 2019 was wrongfully dated to Chalcolithic.

Samples (ACAD ID, Library ID, Archaeological ID, Bone):
Neolithic:

- PT_22194; LP128_2; Camp 2(86) Number 1 Obs L6A; Petrosal
- PT_22200; LP128_4; Camp 1(85) I23A; Petrosal
- PT_22204; LP116_11; Camp 2(86) number 9-12 Obs K9A; Petrosal
- PT_22207; LP116_12; Camp 2(86) Number 2-4 M9; Petrosal
- PT_22210; LP116_13; Skull CLCB33; Petrosal
- PT_22212; LP123_7; Camp 2(86) number 1-2 I9B; Petrosal
- PT_22213; LP116_15; Camp 2(86) number 1 I9A; Petrosal
- PT_22214; LP116_16; Camp 2(86) number 1-3 L10; Petrosal

Chalcolithic

- PT_22197; LP128_3; Camp 2(86) Number 1-3 M10; Petrosal

*Olalde I, Mallick S, Patterson N, Rohland N, Villalba-Mouco V, Silva M, et al. The genomic history of the Iberian Peninsula over the past 8000 years. Science. 2019;1234(March):1230–4.*

#### 1.2. Torre Velha 3 (Chalcolithic and Bronze Age)

Torre Velha 3 (Beja) is a multi-period site, dating from the Chalcolithic to Late Antiquity in the Alentejo region. Burial structures dated to the Chalcolithic and Bronze Age include pits and Hypogea. Most burials were found in a flexed position. In many of the Hypogea, burials were accompanied by grave goods, including pottery, metal artifacts and animal offerings.

We sampled a single individual from the Chalcolithic, found in an individual pit. According to anthropological and paleogenomic analysis, the skeleton belongs to an adult female.

Burials dated to the Bronze Age were recovered from pits and Hypogea, both with a variable number of individuals. The only pit analyzed in this study displayed a minimal number of three individuals. The first burial, found in a flexed position at the base of the pit, belonged to an adult of unknown sex (PT_23209), which our analysis confirmed to be genetically female. Above, two skeletons were apparently deposited simultaneously. The remaining skeletons from this period were unearthed from Hypogea formed by an antechamber and a chamber, the majority of which revealed two burials (one *in situ* and another resulting from a reduction of a previous burial). Here, we sampled one individual burial from an hypogeum located in the antechamber (PT_23210), consisting of an adult male according to anthropological analysis (and here confirmed by genetic sex determination). An additional two samples were obtained from individual burials at Hypogea (PT_23208 and PT_23211), consisting of adult individuals of both sexes according to anthropological analysis (and here confirmed by genetic data). One Hypogea revealed a double *in situ* inhumation of non-adults with associated reduction from a previous adult burial. We sampled one of the non-adults for genetic analysis (PT_23212).

Chalcolithic:
Samples (ACAD ID, Library ID, Archaeological ID, Bone):
- PT_23206; LP113_11; TV3 2015 39; Petrosal

Bronze Age:
Samples (ACAD ID, Library ID, Archaeological ID, Bone):
- PT_23210; LP112_10; TV3. 1361; Petrosal
- PT_23207; LP111_6; TV3 1789 Hipogeu; Petrosal
- PT_23208; LP111_5; TV3 1799 Hipogeu; Petrosal
- PT_23209; LP112_15; TV3. Dec. 2392. 24; Petrosal
- PT_23211; LP116_1; TV3 2051 Hipogeu; Petrosal
- PT_23212; LP123_14; TV3 2368 Hipogeu; Petrosal

#### 1.3. Monte da Cabida 3 (Bronze Age)

The Necropolis of Monte da Cabida 3 (São Manços, Évora, Portugal) was discovered in 2004, and excavated during the year of 2007. Human remains were unearthed from rectangular cists (sep.) and pit burials (fossa), all dated to the Bronze Age. From these burials, some were discovered with signs of perturbation, containing a minimum number of individuals between one and five, including non-adults and adults of both sexes.

One individual (I7692) was previously genetically analyzed by Olalde et al. 2019. Here, we sequenced an additional two ancient genomes using the Twist Bioscience “Twist Ancient DNA” kit reagent.

Samples (ACAD ID, Library ID, Archaeological ID, Bone):

- PT_23199; LP113_3; 3 Fossa 64; Petrosal **= I7689 Olalde et al. 2019**
- PT_23197; LP113_8; 3 Sep 8 872; Petrosal
- PT_23198; LP123_3 + LP113_5; Fossa 44 [945]; Petrosal **= I7691 Olalde et al. 2019**
- PT_23200; LP113_10; 3 Fossa 41 931; Petrosal
- PT_23203; LP116_6; 3 Fossa 41 932; Petrosal
- PT_23202; LP115_14; 3 Sep 10 875; Petrosal
- PT_23201; LP123_13; 3 Sep 9 905; Petrosal
- PT_23205; LP116_3; 3 Sep 9 904; Petrosal
- PT_23204; LP116_2; Sep7 700; Petrosal

*Olalde I, Mallick S, Patterson N, Rohland N, Villalba-Mouco V, Silva M, et al. The genomic history of the Iberian Peninsula over the past 8000 years. Science. 2019;1234(March):1230–4.*

**1.4. Outeiro Alto (Bronze Age)**

Outeiro Alto 2 (Brinches, Serpa, Portugal) is part of a wider range of archaeological sites excavated on the left bank of the Guadiana River, in Baixo Alentejo. The chronology of its occupation extends from the Final Neolithic to the Bronze Age. The individuals analyzed here belong to Núcleo-C and were sampled from the Hypogea necropolis dated to the Bronze Age.

Samples (ACAD ID, Library ID, Archaeological ID, Bone):
- PT_23219; LP123_11; 2608; Petrosal
- PT_23218; LP115_12; 505; Petrosal

**1.5. Idanha-a-Velha - Espírito Santo (2nd-7th century)**

Archeological assemblage recovered in two different campaigns, accounting for a total of eight individuals, two exhumed in 1995 at *Palheiros* (not considered here) and another six unearthed in 1997 at the *Espírito Santo* street (Idanha-a-Velha, Portugal). This location is placed outside the wall that has fortified Idanha, at least, since the end of the 1st century BC (Mantas, 2006; Carvalho, 2022) and to the right of the north gate of this fortification.

Sample (ACAD ID, Library ID, Archaeological ID, Bone, Age):
- PT_24182; LP115_2 & LP123_10; Espítio Santo nº34 Ent. 4 Cont 61; Petrosal; 2nd-3rd century (110-236 cal AD; 1840 - 1714 cal BP)
- PT_22987; LP111_4; Espírito Santo nº 34, Ent 5, Cont 62; Petrosal; 5th-7th century
- PT_22988; LP123_1; Espírito Santo nº 30, Ent 7, Cont 56; Petrosal; 4th-5th century

According to the Anthropological analysis:
- Espítio Santo nº34 Ent. 4 Cont 61: This non-adult individual has a reasonable bone preservation (29.7%), including well-preserved dentition that allowed to estimate the age-at-death between 10.5 postnatal months to 1.5 years (AlQahtani et al., 2010). The absence of the auricular surface of the ilium prevented the estimation of biological sex from anthropological analysis. In paleopathological terms, only occipital endocranial growths were recorded, as well as some bone reactive foci on the medial surfaces of both tibias, possibly related to children's physiological periostitis.
- Espírito Santo nº 34, Ent 5 is a well-preserved (41.2%) perinatal individual (femoral age: 38±1,5 fetal weeks; dental age: 1.5 postnatal months). Metrically, the auricular surface of the ilium indicated a biological male individual. No bone changes consistent with pathology were observed.
- Espírito Santo nº 30, Ent 7 is very poorly preserved (14.8%), except for the dentition (21/28 teeth) that allowed for an estimated age-at-death of 10.5-11.5 years.

*AlQahtani SJ, Hector MP, Liversidge HM. Brief communication: The London atlas of human tooth development and eruption. Am J Phys Anthropol. 2010 Jul;142(3):481-90. doi: 10.1002/ajpa.21258. PMID: 20310064.*

*Carvalho PC. Igaedis, a capital da civitas Igaeditanorum. In: Ciudades Romanas de Hispania II, Hispania Antigua, Serie Arqueológica. Roma – Bristol: Trinidad Nogales.*
*Mantas V. Cidadania e Estatuto Urbano na civitas Igaeditanorum (Idanha-a-Velha). Biblos (2006) vol. IV, 2.ª série, 49-92. Basarrate: Museo Nacional de Arte Romano, L'erma di Bretschneider; 2022. p. 393–412.*

#### *1.6. Idanha-a-Velha - Porta Norte (Roman)*

Small necropolis (4 adult and 1 non-adult individuals) located near the northern gate of the fortified wall of Idanha-a-Velha.

Samples (ACAD ID, Library ID, Archaeological ID, Bone):
- ID_25510; LP129_5; Ind.5 (Porta Norte); Petrosal
- ID_25538; LP127_12; Ind.4 (Lado Nascente); Petrosal
- ID_25537; LP127_11; Ind.1 (Chão da muralha) Casa do Átrio; Petrosal

According to the Anthropological analysis:
- Ind.1 is a 40–44-year-old male inhumed directly into the soil, in dorsal decubitus, and orientated W (head) to E (feet) without any grave goods. A rock was placed next to the right side of the face. The skeletal preservation is reasonable (31.9%), with no pathological signs recorded.
- Ind. 4 is a poorly preserved non-adult individual (3.1%). However, the absence of fusion between the occipital scale and the left pars lateralis allowed for an estimation of the age-at-death of <3 years. The biological sex could not be assigned, and no pathology was recorded.
- Ind.5 is a very well-preserved skeleton (79.8%) of an adult female (30-34 years) inhumed and orientated in the exact same manner as Ind.1 (see above). The mean stature was estimated at 154,7±2,62 cm. This individual had a relatively important set of bone alterations, which have been described in Monge Calleja et al., in prep.

#### *1.7. Idanha-a-Velha - Porta Sul (Roman)*

Small necropolis located near the southern gate of the fortified wall of Idanha-a-Velha.

Samples (ACAD ID, Library ID, Archaeological ID, Bone, Age):
- ID_25536; LP127_10; Ind.2; Petrosal
- ID_25522; LP126_14; Ind.1; Petrosal

#### *1.8. Castro de Avelãs - Torre Velha (6th - 12th century)*

The archaeological site is located in north-eastern Portugal. Archaeological excavations at the Torre Velha necropolis (2012-2015) have so far revealed 59 graves directly excavated in the schist rock, which is typical of Medieval sites in Iberia, with a minimum number of individuals estimated at 57, of which 39 individuals were exhumed from individual graves (33 adults; 6 non-adults) and 18 from ossuaries (17 adults; 1 non-adult). Radiocarbon dating on a total of 16 individuals demonstrated these individuals were buried between the 6$^{th}$ and 13$^{th}$ century CE (Cal AD ~500-1300).

Samples (ACAD ID, Library ID, Archaeological ID, Bone, Age):

- PT_23002; LP112_13; TVCA/12 UE [02] R9; Petrosal; 7th-9th century
- PT_22994; LP117_3; TVCA/12 UE [15] R9; Petrosal; 11th-12th century
- PT_22992; LP117_2; TVCA/12 - UE [50]; Petrosal; 11th-12th century
- PT_22997; LP111_2; O0/O1 UE [46]; Petrosal; 6th-7th century
- PT_22989; LP117_1; TVCA/12 ossário 54 crânio 38; Petrosal; 11th-12th century
- PT_23001; LP117_4; TVCA/12 AC7 Ind EU [16]; Petrosal; 7th-8th century

#### *1.9. Guarda - Prazo (Freixo de Numão) (7th - 13th century)*

The necropolis of Prazo is located near the locality of Freixo de Numão, in the district of Guarda, in northern Portugal. Anthropological and archaeological excavations were carried out between 1995 and 1997, and according to archeological and funerary findings, it was in use at least between the 5th and the 13th centuries AD, associated to Christian burial practices. The skeletal remains excavated from 22 graves and ossuaries represent at least 75 individuals of both sexes, including adults and non-adults.

Samples (ACAD ID, Library ID, Archaeological ID, Bone, Age):

- PT_23235; LP123_4; PZ/S.21/1; Petrosal; 7th - 9th century
- PT_23237; LP123_16; PZ/S.12/94; Petrosal; 12th - 13th century
- PT_23238; LP117_6; PZ/S.15/B01; Petrosal; 9th - 12th century
- PT_23236; LP123_12; PZ/S.4/101; Petrosal; 9th - 11th century

Detailed information can be found in:

*Cunha, E.; Matos, V. Dados bioarqueológicos para o conhecimento dos habitantes do sítio do Prazo (Freixo de Numão) durante a Idade Média. In: Sá Coixão, A. N. (ed.) Rituais e cultos da morte na região de Entre Douro e Côa. Freixo de Numão: A.C.D.R de Freixo de Numão (1999), pp. 101-128. https://hdl.handle.net/10316/80422*

*Matos, V.; Cunha, E. A Necrópole do Prazo no contexto das necrópoles medievais Portuguesas. Côavisão: Cultura e Ciência (1999), 1: 45-51. http://hdl.handle.net/10400.26/25510*

#### *1.10. Santarém - Islamic Necropolis (8th - 10th century)*

The necropolis of Avenida 5 de Outubro #2-8 was excavated during 2007 and 2008. It was used by the Romans, the Visigoths and by the earliest Muslim inhabitants of Santarém after the Islamic conquest (8th and 10th century). The remains of 58 adults and 32 non-adults were recovered from the Islamic necropolis. They were buried on their right side in simple pits, with no grave goods. More information about the site was published in Liberato (2012) and MacRoberts et al. (2024).

Samples (ACAD ID, Library ID, Archaeological ID, Bone, Age):
- PT_24170; LP114_10; 5 Out ue2334; Petrosal
- PT_24173; LP114_13; 5 Out ue931; Petrosal
- PT_24169; LP123_9; 5 Out ue1584; Petrosal

- PT_24171; LP114_11; 5 Out ue1092; Petrosal
- PT_24172; LP114_12; 5 Out ue1647; Petrosal
- PT_24174; LP114_14; 5 Out ue955; Petrosal

*Liberato M. Novos dados sobre a paisagem urbana da Santarém medieval ( séculos V-XII ): a necrópole visigoda e islâmica de Alporão. Medievalista (2012) ;11. https://doi.org/10.4000/medievalista.803*

*MacRoberts, R. A., Liberato, M., Roca-Rada, X., Valente, M. J., Relvado, C., Matos Fernandes, T., Barrocas Dias, C., Llamas, B., Vasconcelos Vilar, H., Schöne, B. R., Ribeiro, S., Santos, J. F., Teixeira, J. C., & Maurer, A.-F. Shrouded in history: Unveiling the ways of life of an early Muslim population in Santarém, Portugal (8th– 10th century AD). PLOS ONE (2024), 19(3), e0299958-. https://doi.org/10.1371/journal.pone.0299958*

#### *1.11. Loulé - Hospital da Misericordia (11th - 13th century)*

The Muslim cemetery of Hospital da Misericórdia was identified at the eastern edge of Loulé and its excavation in 2009 revealed that it was functioning in the 10$^{th}$ to 13$^{th}$ century. The excavation of thirty-seven graves allowed the identification of 11 non-adults and 25 adults. All individuals were laying in right lateral decubitus. Five secondary inhumations were also identified. Archaeological and anthropological information were published in (Pires & Benisse, 2010; Pires & Luzia, 2017).

Samples (ACAD ID, Library ID, Archaeological ID, Bone, Age):

- PT_24175; LP114_15; LHM 17; Petrosal; 11th - 12th century (Pires & Luzia, 2017)
- PT_24176; LP115_3; LHM 31; Petrosal
- PT_24177; LP114_16; LHM 12; Petrosal

*Pires, A., & Benisse, V. A intervençao no Hospital da Misericórdia de Loulé: contributo para a percepção da organização espacial da cidade medieval. Xelb (2010), 10, 437–454.*

*Pires, A., & Luzia, I. As Necrópoles Islâmicas de Loulé. In Loulé. Territórios, Memórias, Identidades (pp. 494–503). Museu Nacional de Arqueologia (2017).*

#### *1.12. Loulé - Quinta da Boavista (12th - 13th century)*

The Muslim cemetery of Quinta da Boavista was excavated in 1999. It is located North of Loulé and was used during the 11$^{th}$ to the 13$^{th}$ century. The excavation uncovered 41 human graves where 43 individuals were buried, including 36 adults and 7 non-adults. No grave goods were found and some variations in body depositions were observed. Archaeological and anthropological information were published in (Cunha et al., 2000, 2001; Luzia, 1999, 2002; Pires & Benisse, 2010; Pires & Luzia, 2017).

Samples (ACAD ID, Library ID, Archaeological ID, Bone, Age):
- PT_24179; LP115_5; LQB 1; Petrosal
- PT_24181; LP115_6; LQB 31; Petrosal
- PT_24180; LP115_1; LQB 17; Petrosal
- PT_24178; LP115_7; LQB 9; Petrosal

*Cunha, E., Marques, A., & Silva, A.. O passado em Al'-Ulyã. Estudo Antropológico de um cemitério muçulmano. Relatório técnico-científico (2000).*
*Cunha, E., Marques, C., & Silva, A. O passado em Al'-Ulyã: Estudo Antropológico de uma população muçulmana. Al-Ulya (2001), 8, 35–49.*
*Luzia, I. A excavação arqueológica de emergência do cemitério muçulmano da "Quinta da Boavista"/Loulé. Al-Ulya . Al-Ulya (1999), 7, 129–185.*
*Luzia, I. O passado de Al-Ulyã: a escavação arqueológica do cemitério muçulmano. In Património islâmico dos centros urbanos do Algarve: contributos para o futuro (2002) (pp. 151–156).*
*Pires, A., & Benisse, V. A intervençao no Hospital da Misericórdia de Loulé: contributo para a percepção da organização espacial da cidade medieval. Xelb (2010), 10, 437–454.*
*Pires, A., & Luzia, I. As Necrópoles Islâmicas de Loulé. In Loulé. Territórios, Memórias, Identidades (pp. 494–503). Museu Nacional de Arqueologia (2017).*

#### *1.13. Santarém - Rua dos Barcos (12-14th century)*

The archaeological and anthropological intervention carried out between April and June of 2003, allowed the exhumation of 147 individuals, 118 adults and 29 non-adults. Associated with the human remains, ceramics from various periods, coins and two earrings were recovered. In addition to the aforementioned material, a large ossuary associated with the burials was also recovered. The selected samples include adult and non-adult individuals.

Samples (ACAD ID, Library ID, Archaeological ID, Bone, Age):
- PT_23226; LP113_4; RB83; Petrosal; 13-14th century
- PT_23229; LP123_2; RB93; Petrosal; 13-14th century
- PT_23227; LP123_6; RB39; Petrosal; 13-14th century
- PT_23230; LP112_16; RB20; Petrosal; 13-14th century
- PT_23223; LP117_7; RB59; Tooth; 13-14th century
- PT_22219-23224; LP117_12+LP123_5; Esq 18; Petrosal; 13-14th century
- PT_22220; LP111_3; Esq 52; Petrosal; 13-14th century
- PT_23221; LP117_10; RB Ent 95; Tooth; 13-14th century
- PT_22218; LP113_1; Esq 29; Petrosal; 13-14th century
- PT_23234; LP116_7; RB142; Petrosal; 12th century
- PT_23220; LP117_8; RB E68; Tooth; 13-14th century
- PT_23222; LP117_9; RB 86; Tooth; 13-14th century
- PT_23228; LP116_4; RB143; Petrosal; 13-14th century
- PT_23231; LP115_13; RB130; Petrosal; 13-14th century
- PT_23225; LP123_15; RB139; Petrosal; 13-14th century
- PT_23232; LP116_5; RB144; Petrosal; 12th century

#### *1.14. São Miguel de Odrinhas (12-15th century)*

S. Miguel de Odrinhas is a Christian necropolis located in close proximity to the chapel of São Miguel, near the village of Odrinhas, in the municipality of Sintra (central Portugal). This extensive necropolis was partially excavated during four archaeological campaigns: one in the 1940s and others in 1988, 1997, and 2005. Over the last three campaigns, 40 graves were uncovered, yielding osteological material from 157 individuals. In the earlier excavation, 50 graves were opened, though the location of the human remains is currently unknown.

Anthropological analysis of the 157 individuals revealed the presence of 73 adults (36 males and 26 females) and 84 non-adults.

Samples (ACAD ID, Library ID, Archaeological ID, Bone):
- PT_24164; LP114_4; SMO99 Sep1 Oss2; Petrosal
- PT_24165; LP114_5; SMO2005 Sep9 Ent; Petrosal
- PT_24163; LP114_3; SMO99 Sep1 Oss4; Petrosal
- PT_24166; LP123_8; SMO88 Sep6 Oss1; Petrosal
- PT_24167; LP114_7; SMO97 Sep1 Ent; Petrosal
- PT_24168; LP114_8; SMO88 Sep6 ENT; Petrosal

Fernandes, Teresa. The Medieval Population of S. Miguel de Odrinhas (Sintra). Biological Characterisation. PhD Dissertation (2008). University of Évora.

#### *1.15. Castelo Branco - Church of São Miguel (Late-Medieval/Post-Medieval)*

The Church of São Miguel, now known as "Sé Catedral de Castelo Branco", is a significant Christian site in Castelo Branco, in eastern central Portugal. Documented since the 13th century, it became a cathedral in 1771. An archaeological intervention in 2004 during rehabilitation works uncovered a necropolis with 20 anthropomorphic granite rock-cut tombs in the East church square. Ten individuals (seven adults and three non-adults) were exhumed from nine graves. Although the individual sampled for this study (skeleton 5, grave 6) lacks direct dating, radiocarbon dating of skeleton 8 from grave 4 (Beta-524725) indicates a period between the late $15^{th}$ and $16^{th}$ centuries. Additional information can be found in:

Samples (ACAD ID, Library ID, Archaeological ID, Bone):

- PT_22187; LP128_1; Sep 6, Ent 5; Petrosal

Matos, V.; Marques, C.; Lopes, C. Severe vertebral collapse in a juvenile from the graveyard (13th/14th–19th centuries) of the São Miguel church (Castelo Branco, Portugal): differential palaeopathological diagnosis. *International Journal of Osteoarchaeology (*2011*),* 21(2)*:* 208-217. https://doi.org/10.1002/oa.1125

#### *1.16. Igreja de Santa Maria da Alcáçova, Castelo de Montemor-o-Velho (18th century)*

The first excavation campaign of the Necropolis of the Church of Santa Maria da Alcáçova of the Castle of Montemor-o-Velho occurred in May 2018, under a collaboration between the Department of Life Sciences of the University of Coimbra and the Municipality of Montemor-o-Velho. According to historical documentation, this necropolis was used until 1811, and served as the resting place of people residing north of Montemor-o-Velho, namely in Moinho da Mata and surrounding areas. At present, the archaeological excavations recovered a total of 17 non-adult individuals with estimated ages-at-death varying between a few months and 8 years-old.

Samples (ACAD ID, Library ID, Archaeological ID, Bone):

- PT_23215; LP112_12; MMV Esq 7; Petrosal
- PT_23216; LP112_11; MMV Esq 10; Petrosal
- PT_23242; LP112_9; MMV 1 Esq 5; Petrosal
- PT_23241; LP111_8; MMV 1 Esq 3; Petrosal

- PT_23214; LP113_9; MMV Esq 4; Petrosal
- PT_23217; LP115_8; MMV Esq 16; Petrosal
- PT_23213; LP112_14; MMV Esq 2; Petrosal

#### *1.17. Aveiro - Travanca (18th - 19th century)*

The archaeological intervention of São Mamede in the church of Travanca, a village that belongs to the municipality of Santa Maria da Feira, in Aveiro, was carried out between 2016 and 2017. Despite the use of this Christian space, from the Medieval period (5$^{th}$-15$^{th}$ century CE) to the beginning of the 20th century, only post-mediaeval graves preserved human bone remains. A total of 266 primary burials and 47 ossuaries were recovered from the 412 graves excavated. Among these, two individuals stand out, and were genetically analyzed here:

Individual 403 (PT_22190), a poorly preserved and very fragmented skeleton belonging to an adult male, presented several leprosies related bony lesions, namely in the rhinomaxillary area and feet (Melo et al., 2021). This skeleton, radiocarbon dated from the 17$^{th}$-19$^{th}$ century AD (Beta 514831), was buried in the churchyard, within a wooden coffin and oriented West-East. Forty-seven rosary beads and a cross with a crucified Jesus Christ were found in the abdominal region close to the left forearm.

Individual 255 (PT_22217) stands out by the presence of dental treatment, the only case from this necropolis. This grave was located on the west side of the churchyard, with a confirmed chronology dating to the early 1900s (Beta - 515041; 140 +/- 30 BP; 95.4% probability: 43.1%: 1669 - 1780 cal AD; 36.8%: 1798 - 1891 cal AD; 15.5%: 1908 - 1944 cal AD). The oral health of this individual was compromised by a high number of cariogenic lesions and severe periodontal disease. The occlusal surfaces of three lower teeth with cariogenic lesions were treated with black filling, and gold was used in six anterior and upper teeth.

Samples (ACAD ID, Library ID, Archaeological ID, Bone):

- PT_22190; LP116_10; 403; Petrosal
- PT_22217; LP111_1; 255; Petrosal

Melo, L., Matos, V. M. J., Santos, A. L., Ferreira, C., & Silva, A. M. The first probable evidence of leprosy in a male individual (17th-19th century AD) unearthed in Northern Portugal (Travanca, Santa Maria da Feira). *International Journal of Paleopathology (*2021*)*, 32: 80-86. doi: https://doi.org/10.1016/j.ijpp.2020.12.001

#### *1.18. Santarém - Castelo de Abrantes (18th-19th centuries)*

The church of Santa Maria do Castelo is located within the castle/fortification of Abrantes, a city on the right bank of the Tagus river in central Portugal. In 2015, during an archaeological excavation under the CAST.AB project, an anthropological intervention was conducted to excavate two skeletons (adult males) accidently discovered in the north side of the church's external area. Radiocarbon dating (Beta-514829) indicates these individuals lived between the late $18^{th}$ and the early $19^{th}$ centuries.

Samples (ACAD ID, Library ID, Archaeological ID, Bone):

- PT_22183; LP117_11; Cast. AB 2015, Sond 7, Ent 1; Petrosal
- PT_22179; LP116_9; Cast. AB 2015, Ent 2, Ind 2; Petrosal

### 2. SI Figures

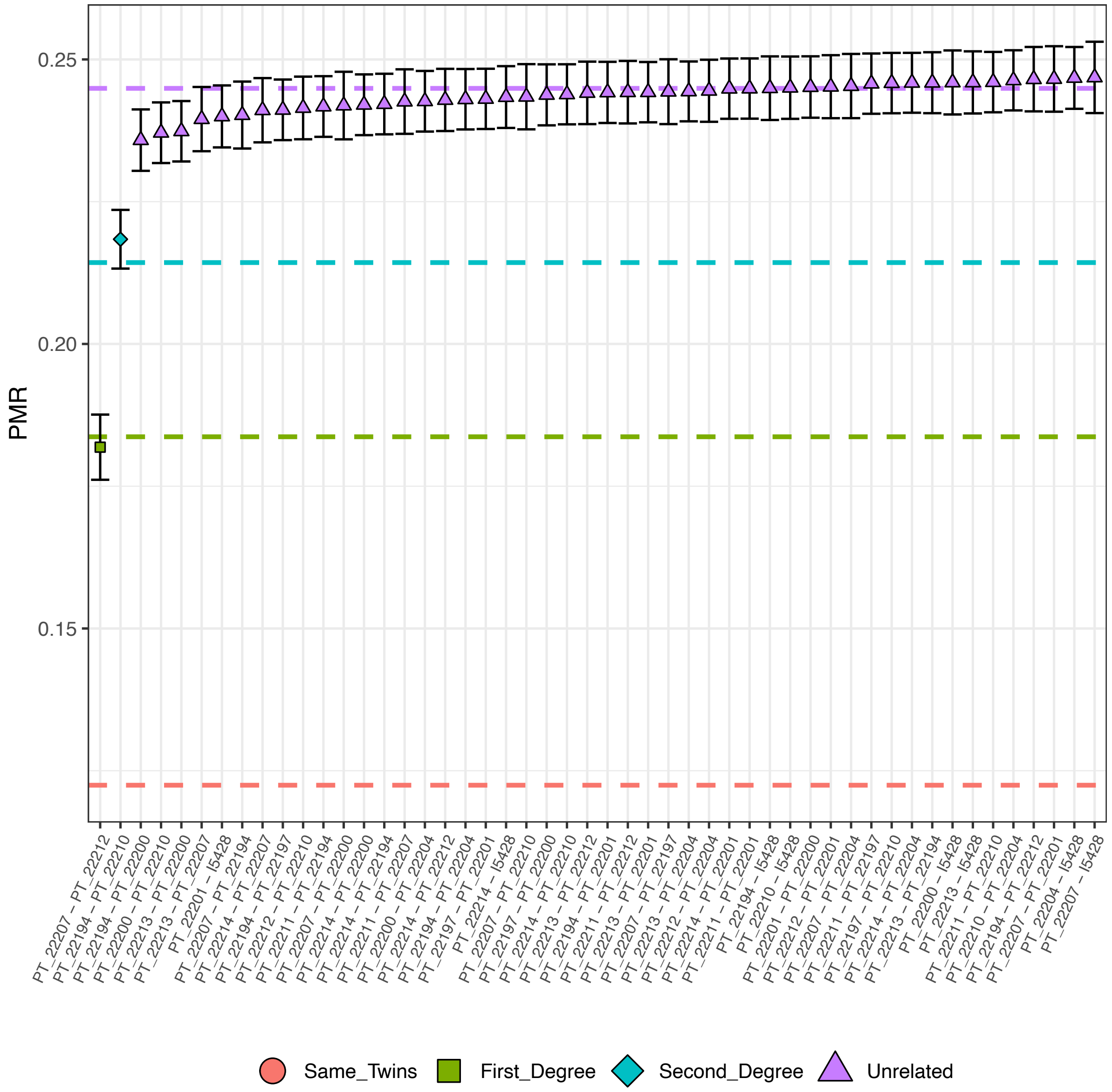

Figure S1. Kinship analysis from Cova_das_Lapas_N/C using BREADR.

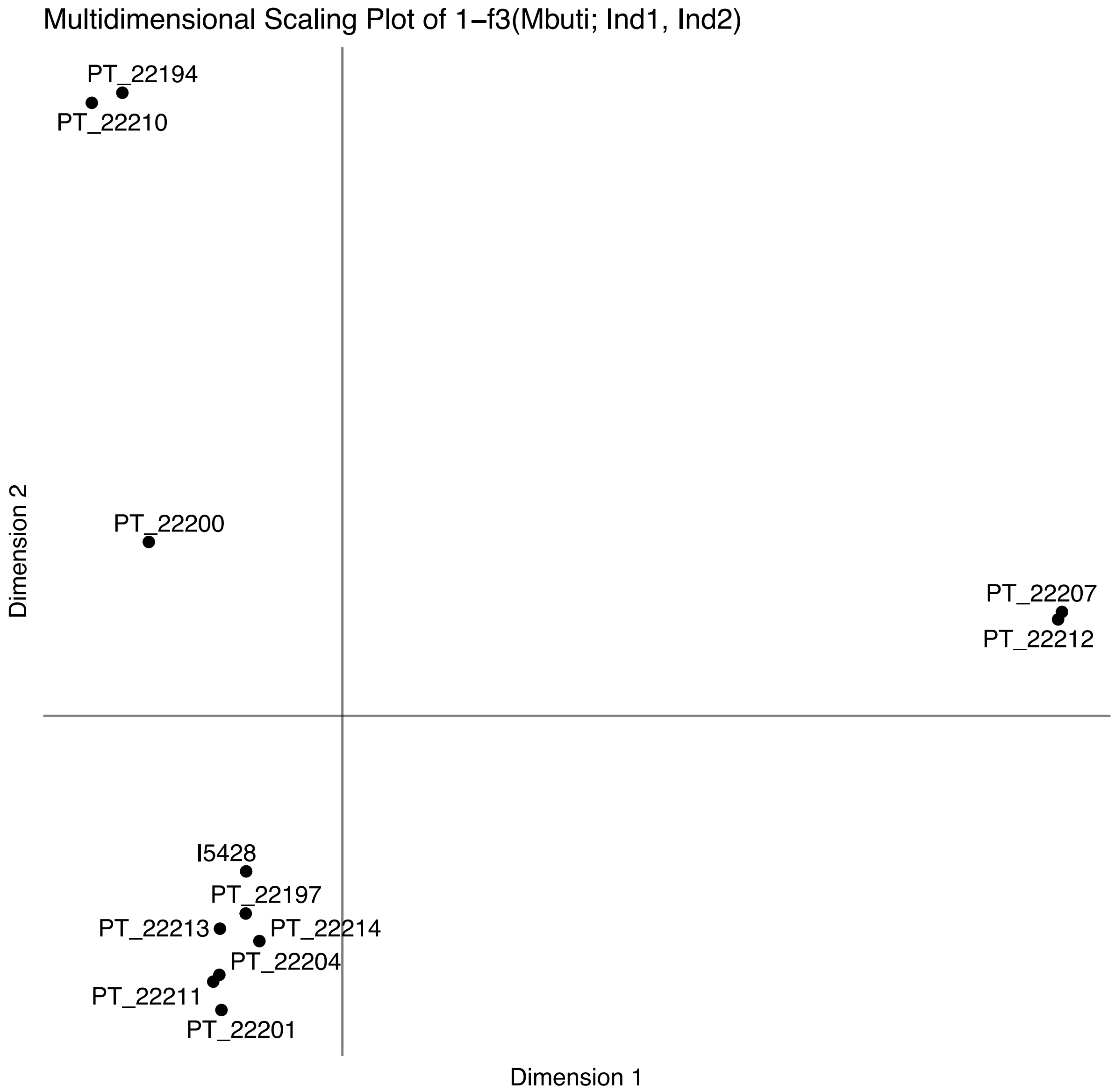

Figure S2. Multidimensional scaling (1-$f_3$) for the Cova_das_Lapas_N/C individuals. Outgroup pairwise $f_3$ computed with Mbuti as outgroup.

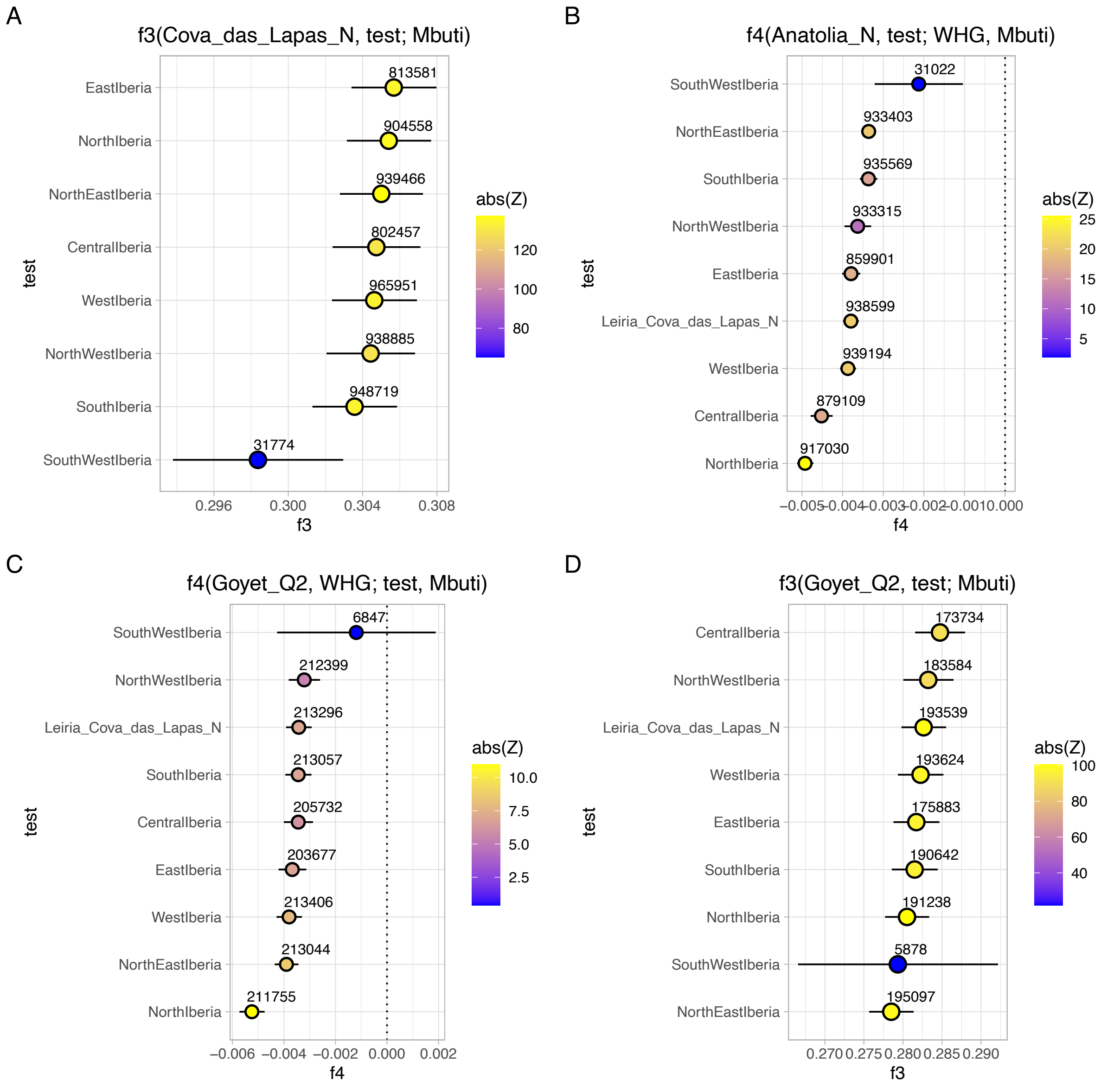

Figure S3: The x-axes represent the $f_3$- or $f_4$-statistic values, with results displayed as the mean ± 1-SD, and colors representing Z-scores. The numbers above each dot indicate the number of SNPs used for each calculation. (A) Outgroup $f_3$-statistics of the form $f_3$(Cova_das_Lapas, Test; Mbuti) where *Test* includes various Iberian Neolithic geographical areas. (B) $f_4$-statistics in the form $f_4$(Anatolia_N, Test; WHG, Mbuti) with *Test* including Cova_das_Lapas_N and different Iberian Neolithic geographical areas. (C) $f_4$-statistics of the form $f_4$(Goyet_Q2, WHG; Test, Mbuti) with *Test* including Cova_das_Lapas_N and different Iberian Neolithic geographical areas. (D) Outgroup $f_3$-statistics of the form $f_3$(Goyet_Q2, Test; Mbuti), where *Test* includes Cova_das_Lapas_N and various Iberian Neolithic geographical areas.

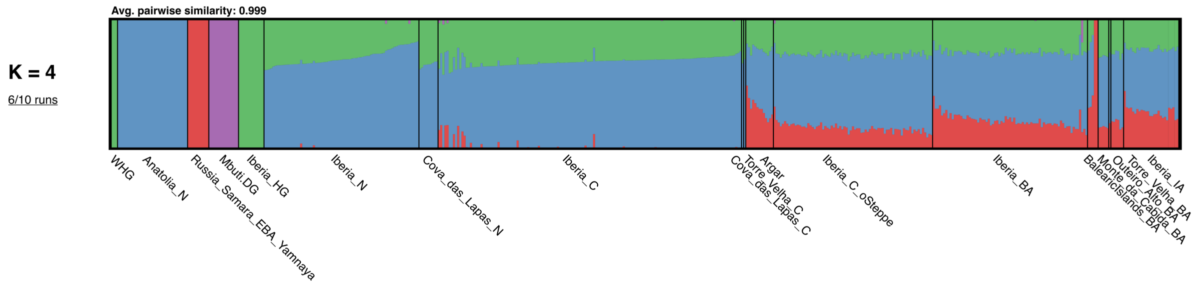

Figure S4. Ancestry proportions estimated using supervised ADMIXTURE (k=4), with the four predefined ancestries being Western Hunter-Gatherers (WHG; green), Neolithic Southwestern Asian farmers (Anatolia_N; blue), Bronze Age Western Steppe herders (Yamnaya_BA; red) and present-day Sub-Saharan Africans (Mbuti; purple). Ancient Iberians whose ancestry proportions were estimated, include individuals from the pre-Neolithic (HG), Neolithic (N), Copper Age (C), Bronze Age (BA) and Iron Age (IA), as well as the individuals from Cova_das_Lapas_N/C, Torre_Velha_C/BA, Outeiro_Alto_BA, and Monte_da_Cabida_BA.

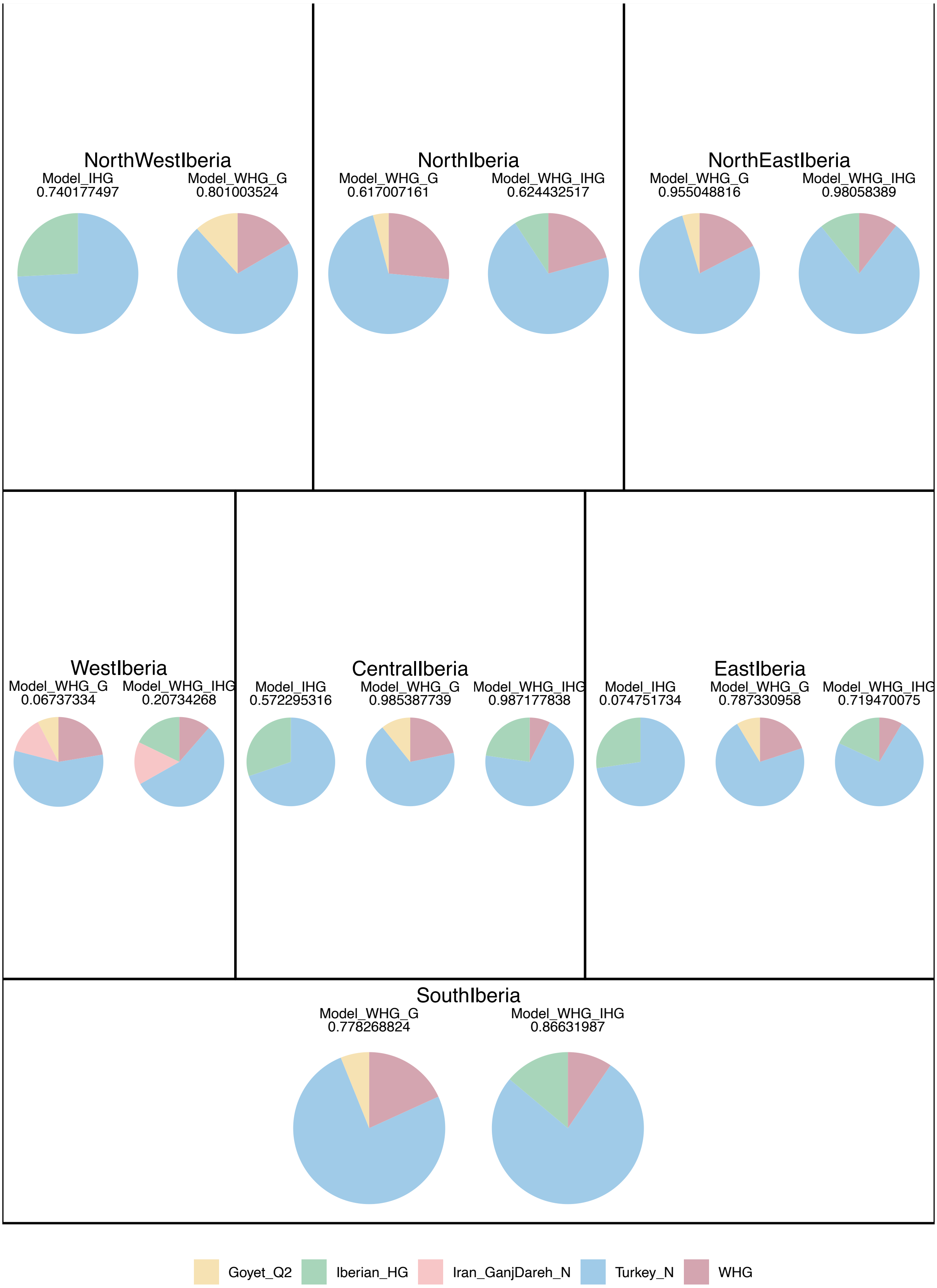

Figure S5. Ancestry proportions for different geographically defined Neolithic Iberian groups using different admixture modeling frameworks. The p-values are provided below the model label.

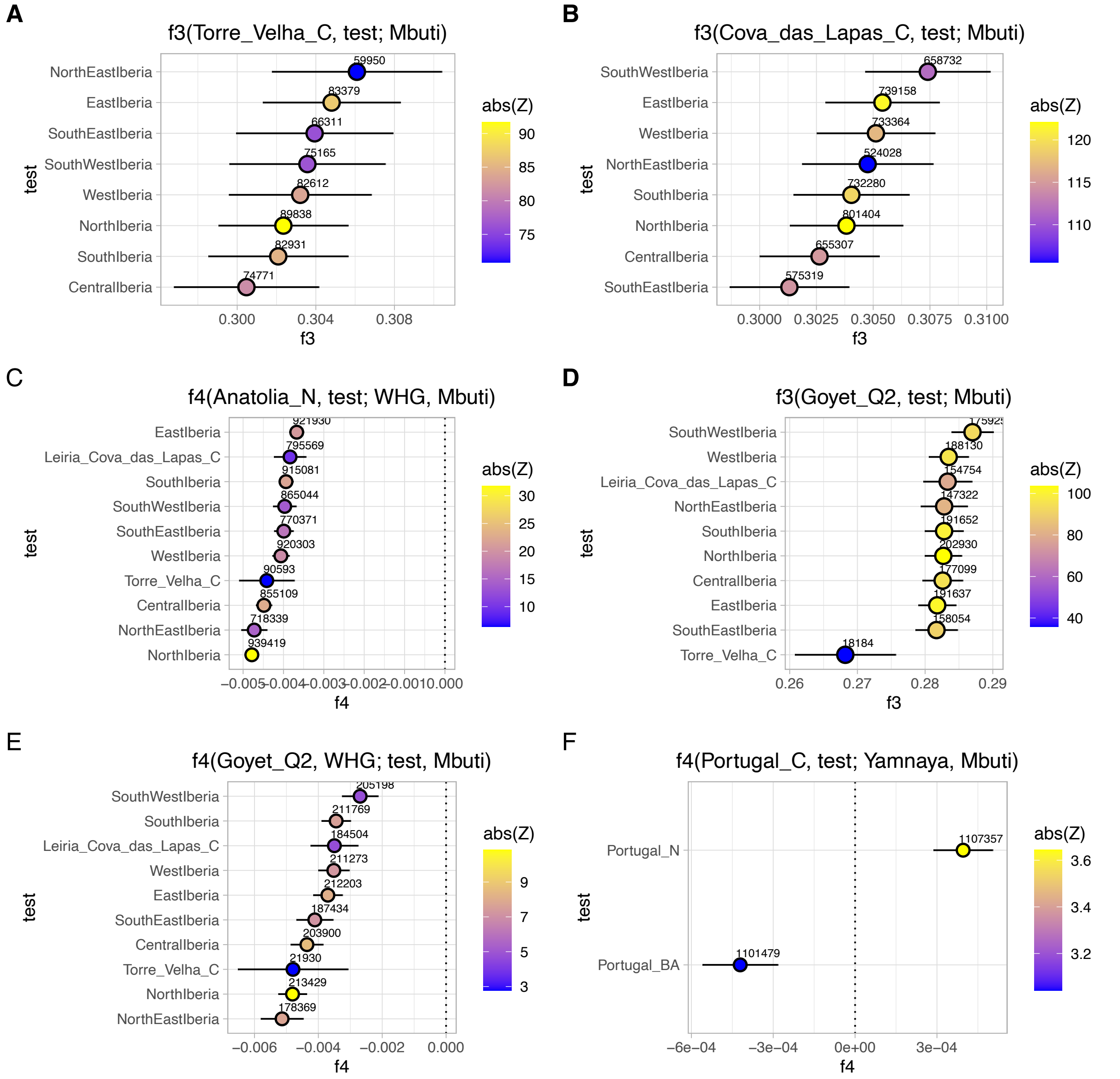

Figure S6: The x-axes represent the $f_3$- or $f_4$-statistic values, with results displayed as the mean ± 1-SD, and colors representing Z-scores. The numbers above each dot indicate the number of SNPs used for each calculation. Outgroup $f_3$-statistics of the form (A) $f_3$ (Torre_Velha_C, Test; Mbuti) and (B) $f_3$(Cova_das_Lapas_C, Test; Mbuti), where *Test* includes various Iberian Chalcolithic geographical areas. (C) $f_4$-statistics in the form $f_4$ (Anatolia_N, Test; WHG, Mbuti) with *Test* including Cova_das_Lapas_C, Torre_Velha_C and different Iberian Chalcolithic geographical areas. (D) Outgroup $f_3$-statistics of the form $f_3$(GoyetQ2, Test; Mbuti), where *Test* includes Cova_das_Lapas_C, Torre_Velha_C and various Iberian Chalcolithic geographical areas. (E) $f_4$-statistics in the form $f_4$ (GoyetQ2, WHG; WHG, Mbuti) with *Test* including Cova_das_Lapas_C, Torre_Velha_C and different Chalcolithic Iberian geographical areas. (F) $f_4$-statistic in the form $f_4$(Portugal_C, Test; Yamnaya, Mbuti), where *Test* includes Portugal_N and Portugal_BA.

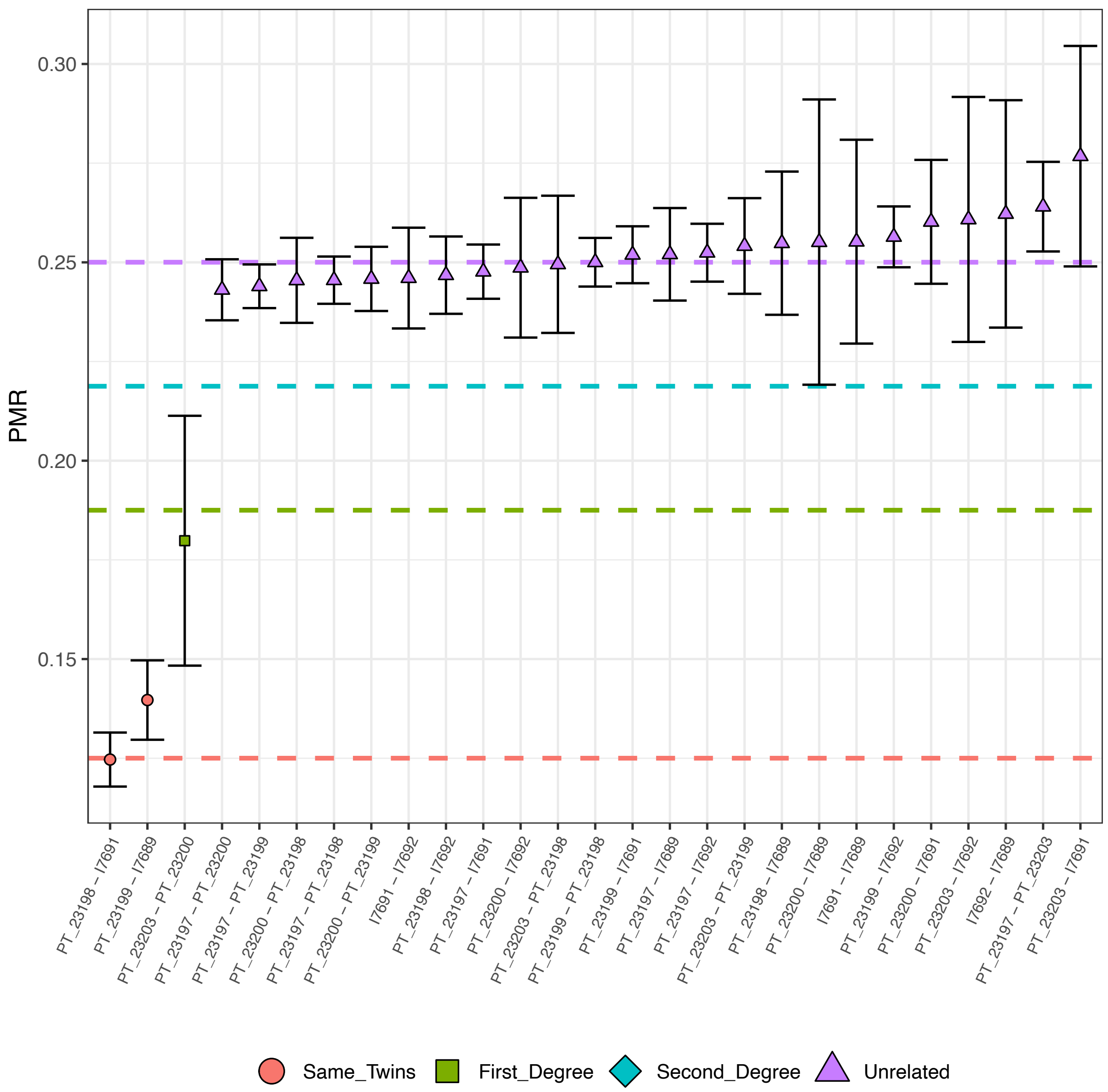

Figure S7. Kinship analysis from Monte_da_Cabida_BA using BREADR.

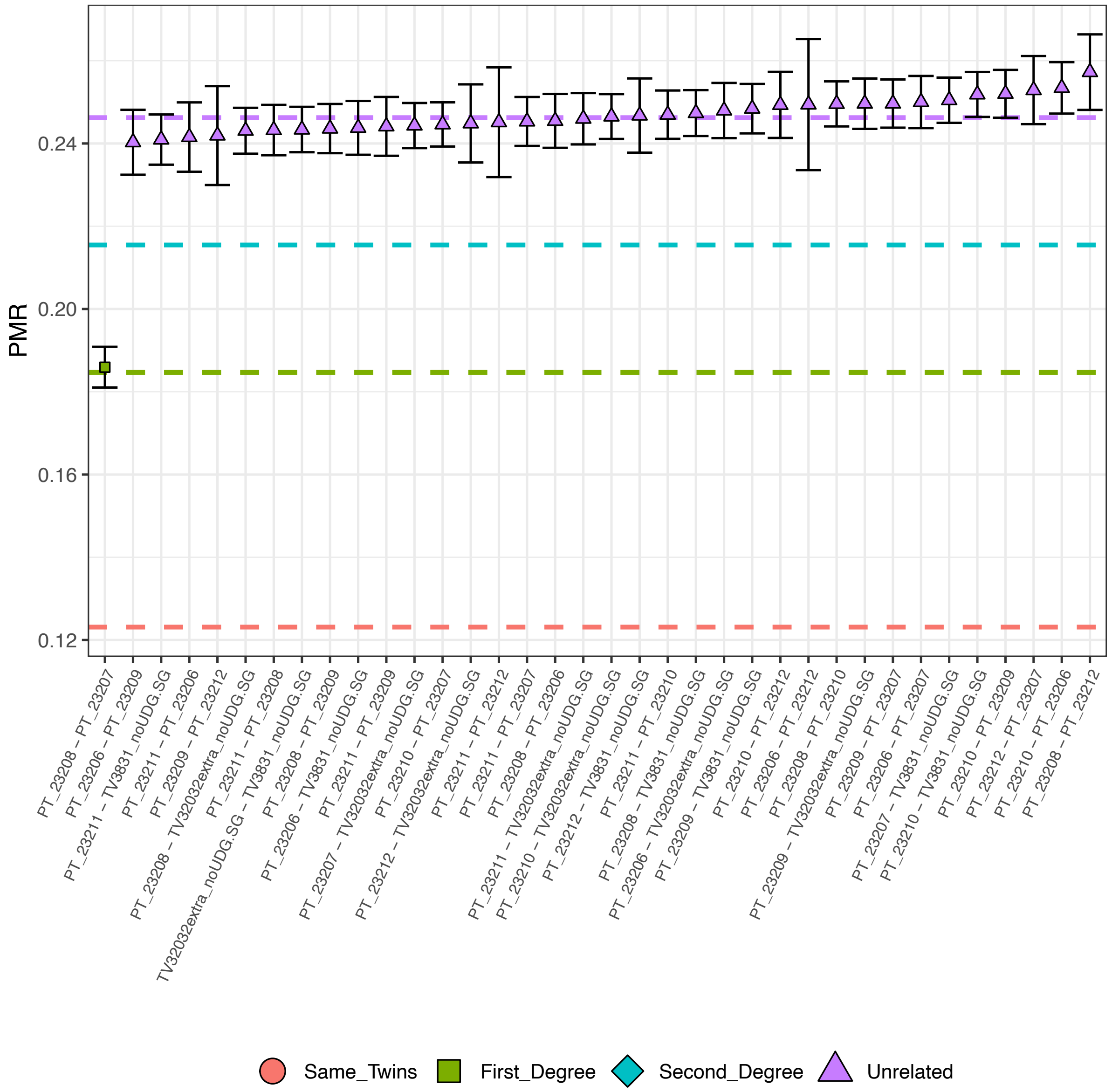

Figure S8. Kinship analysis from Torre_Velha_C/BA using BREADR.

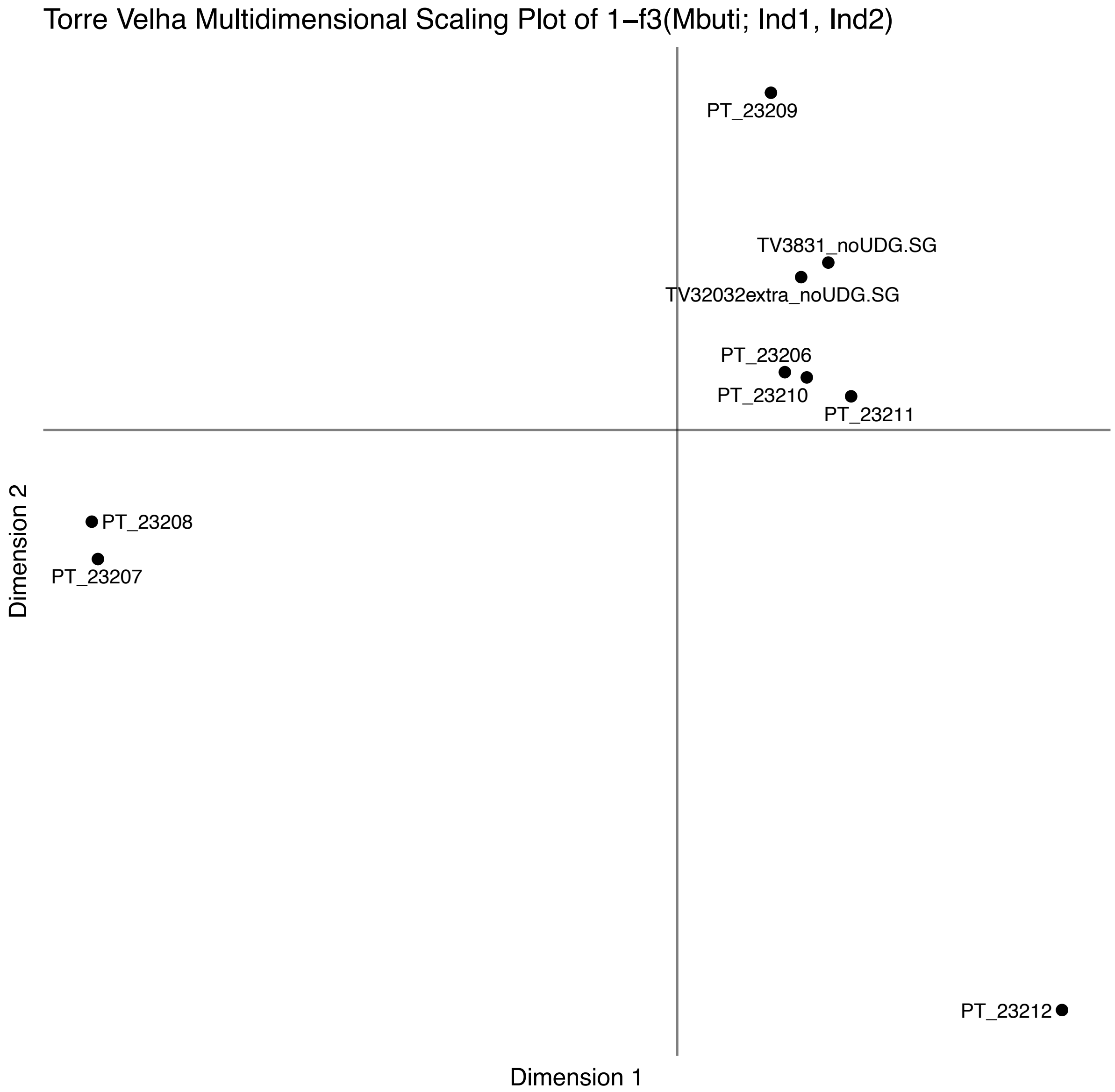

Figure S9. Multidimensional scaling (1- $f_3$) for the Torre_Velha_C/BA individuals. Outgroup pairwise $f_3$ computed with Mbuti as outgroup.

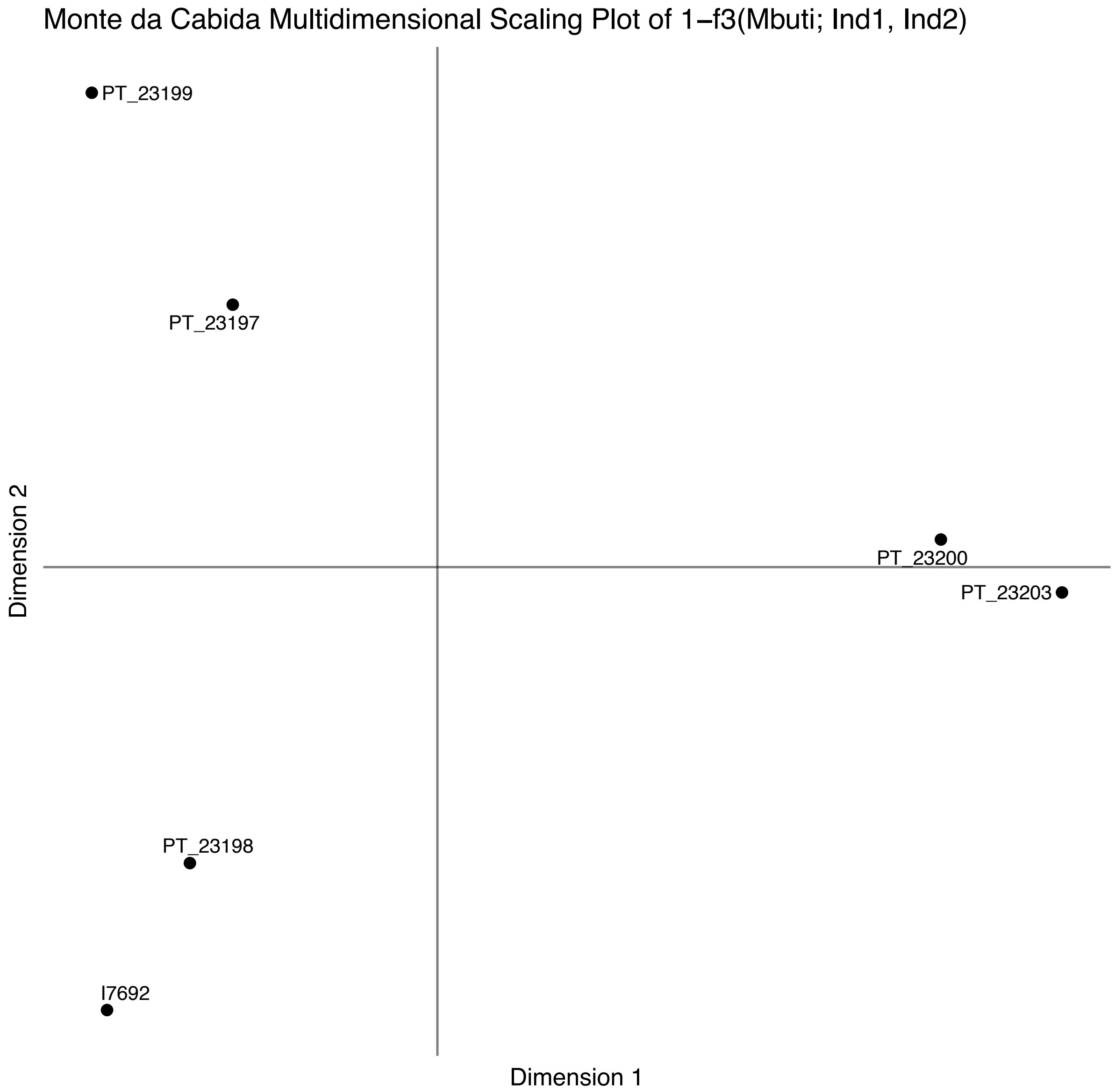

Figure S10. Multidimensional scaling (1- $f_3$) for the Monte_da_Cabida_BA individuals. Outgroup pairwise $f_3$ computed with Mbuti as outgroup.

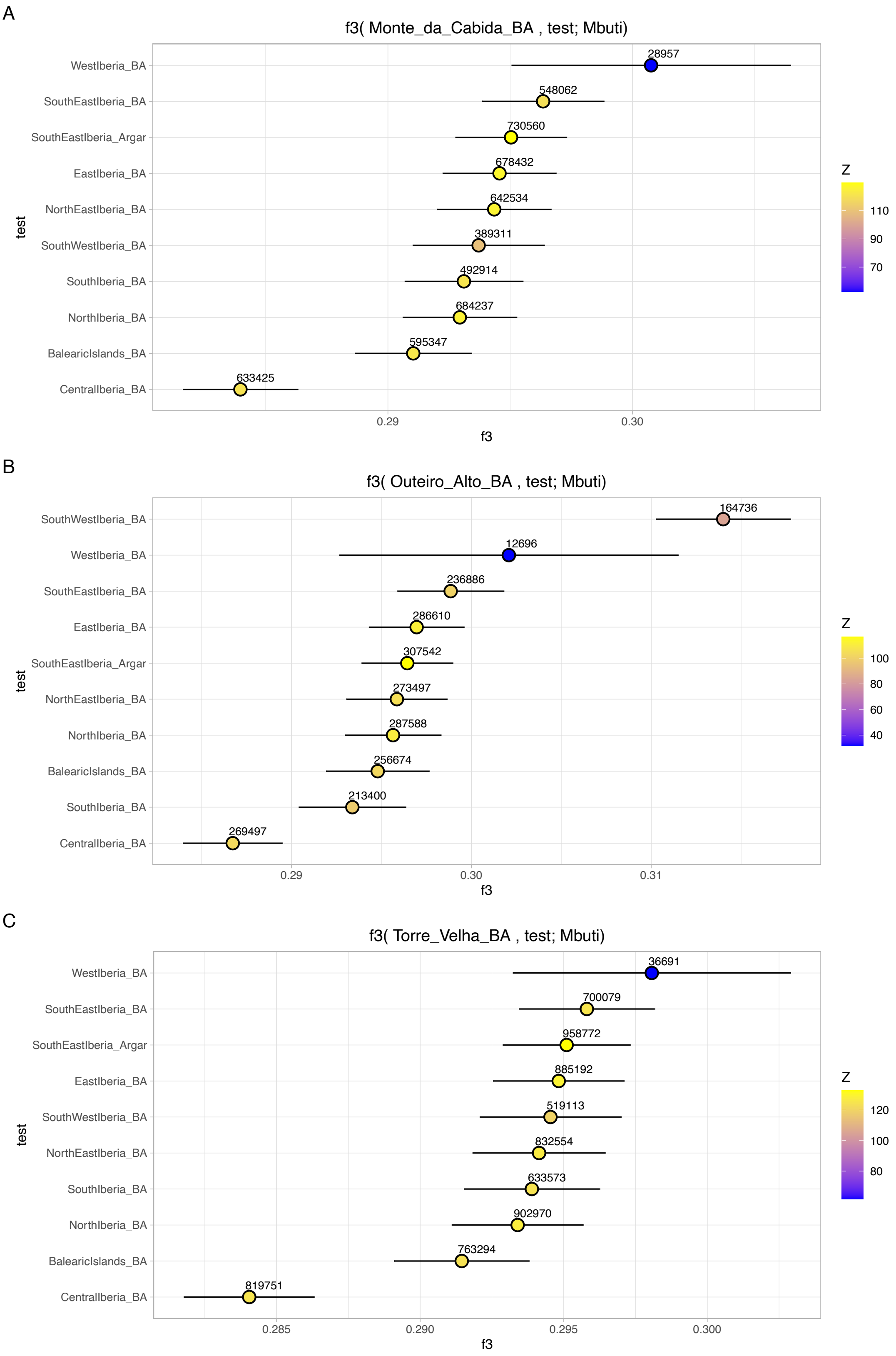

Figure S11. The x-axes represent the $f_3$-statistic values, with results displayed as the mean ± 1-SD, and colors representing Z-scores. The numbers above each dot indicate the number of SNPs used for each calculation. Outgroup $f_3$-statistics of the form (A) $f_3$(Monte_da_Cabida_BA, Test; Mbuti), (B) $f_3$(Outeiro_Alto_BA, Test; Mbuti), and (C) $f_3$(Torre_Velha_BA, Test; Mbuti), where *Test* includes various Iberian Bronze Age geographical areas.

A

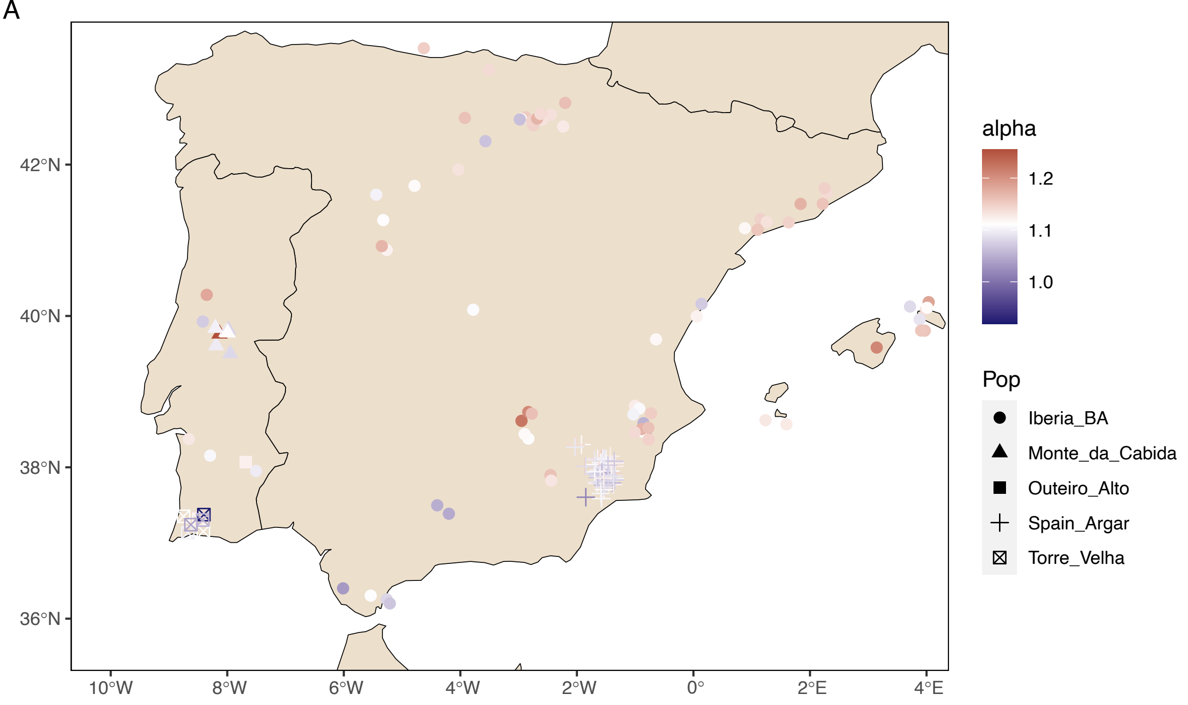

B

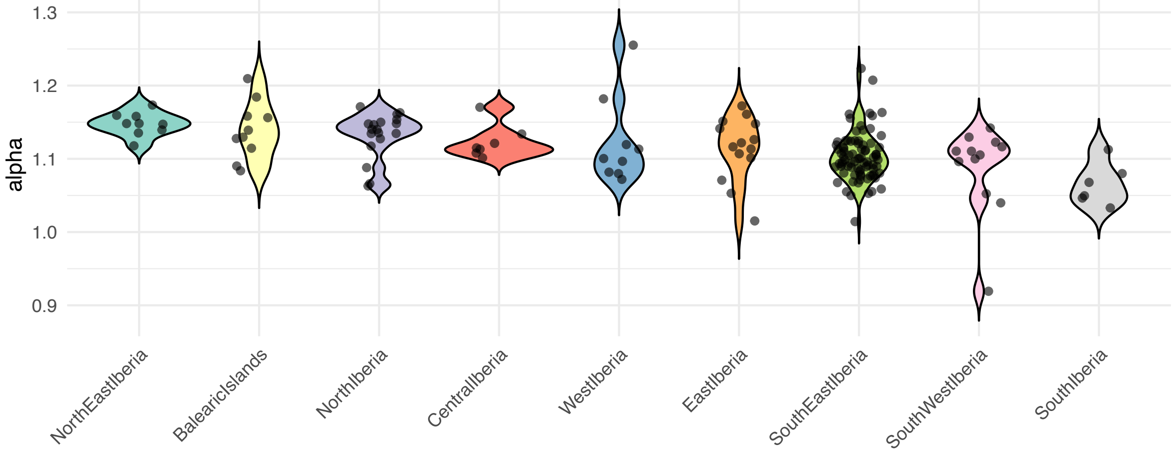

Figure S12. $f_4$-ratios ($f_4$(Yamnaya_Samara, Mbuti; Test, Morocco_Iberomaurusian) / $f_4$(Yamnaya_Samara, Mbuti; Anatolia_N, Morocco_Iberomaurusian)), where *Test* are (A) Iberian Bronze Age individuals as well as Torre_Velha_BA, Outeiro_Alto_BA and Monte_da_Cabida_BA, or (B) different geographically defined Bronze Age Iberian groups. Individuals with absolute Z-scores higher than twice their standard error were excluded. Most $f_4$-ratios exceed 1, indicating, as expected, higher Yamnaya_Samara ancestry than Anatolia_N relative to Morocco_Iberomaurusian, but a slight decrease towards the southwest is found.

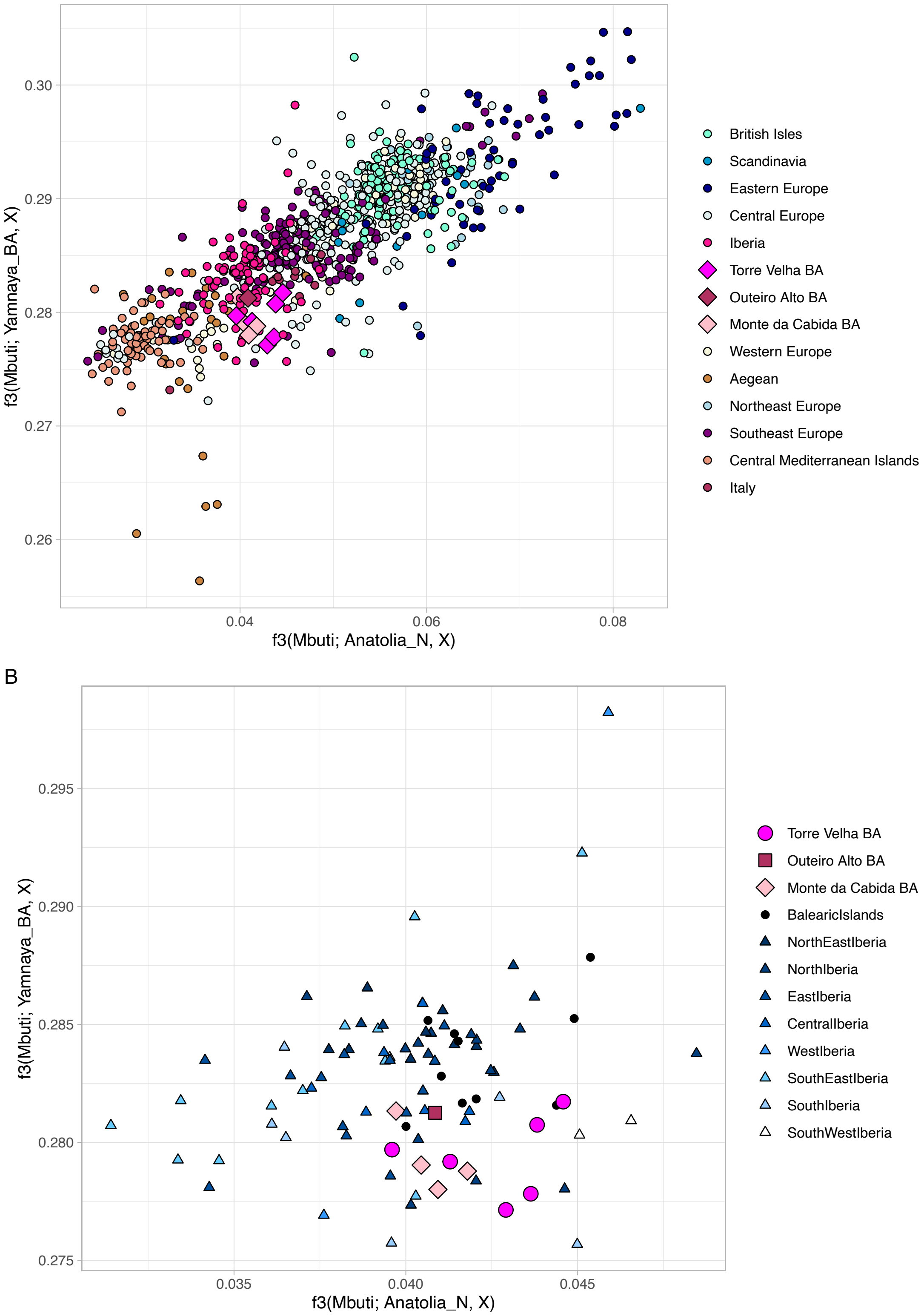

Figure S13. Outgroup *f3*-statistics of the form $f_3$(Mbuti; Pop, X) showing the amount of shared drift between the ancient individuals (X) with Neolithic Anatolian Farmers (Anatolia_N; x-axis) and Steppe-related Bronze Age Eurasians (Yamnaya_BA; y-axis). (A) Clusters of Eurasian individuals are represented in shades of blue, pink, and amber from the top-left to the bottom-right. The Torre_Velha_BA, Outeiro_Alto_BA and Monte_da_Cabida_BA individuals are represented as colored diamonds. (B) Clusters of Iberian individuals are represented in shades of blue on a north-south gradient and the Torre_Velha_BA, Outeiro_Alto_BA and Monte_da_Cabida_BA individuals are represented as colored diamonds as in (A).

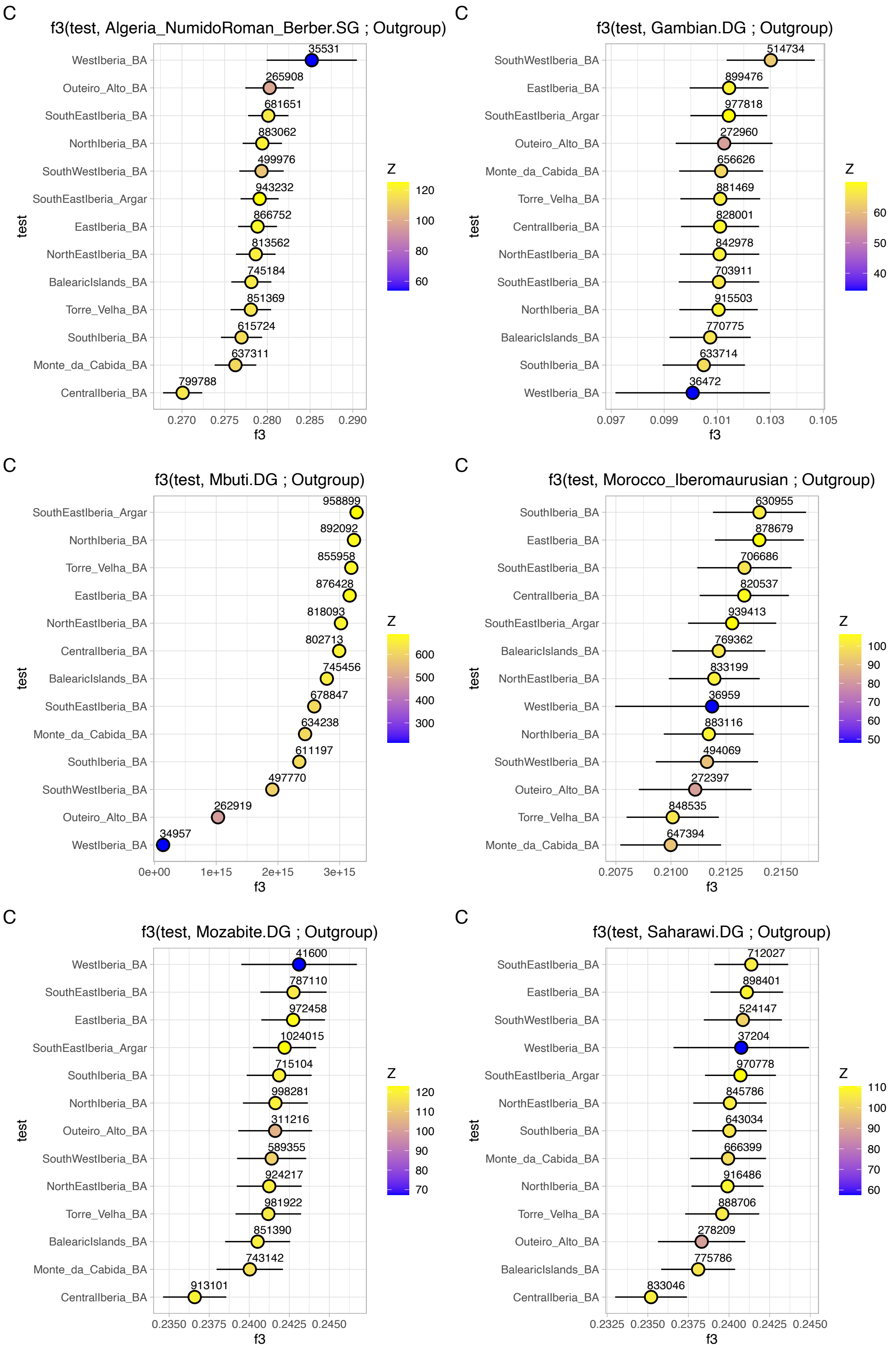

Figure S14. The x-axes represent the *f3*-statistic values, with results displayed as the mean ± 1-SD, and colors representing Z-scores. The numbers above each dot indicate the number of SNPs used for each calculation. Outgroup *f*3-statistics of the form (Test, African_Pop; Outgroup), where *Test* are Torre_Velha_BA, Outeiro_Alto_BA, Monte_da_Cabida_BA and Iberian Bronze Age geographical areas. All outgroups are Mbuti, except when Mbuti is used as *African_Pop*, then the Outgroup is Chimp.

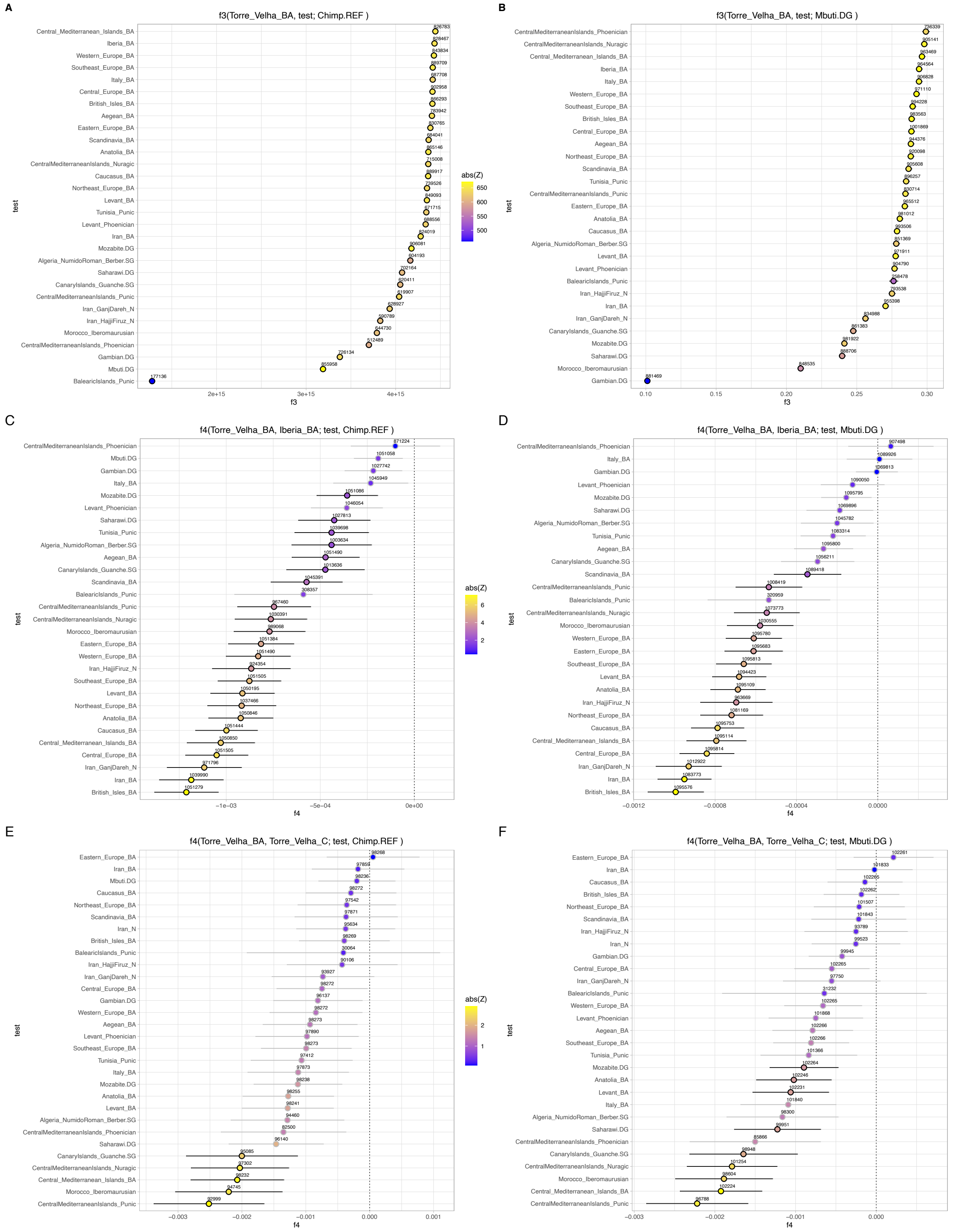

Figure S15. The x-axes represent the $f_3$- and $f_4$-statistic values, with results displayed as the mean ± 1-SD, and colors representing Z-scores. The numbers above each dot indicate the number of SNPs used for each calculation. Outgroup $f_3$-statistics are presented as: (A) $f_3$(Torre_Velha_BA, Test; Mbuti) and (B) $f_3$(Torre_Velha_BA, Test; Chimp), with *Test* including Eurasian and African populations from the Bronze Age or proxies. $f_4$-statistics are shown as: (C) $f_4$(Torre_Velha_BA, Iberia_BA; Test, Mbuti) and (D) $f_4$(Torre_Velha_BA, Iberia_BA; Test, Chimp), and (E) $f_4$(Torre_Velha_BA, Torre_Velha_C; Test, Mbuti) and (F) $f_4$(Torre_Velha_BA, Torre_Velha_C; Test, Chimp), with *Test* including Eurasian and African populations from the Bronze Age or proxies.

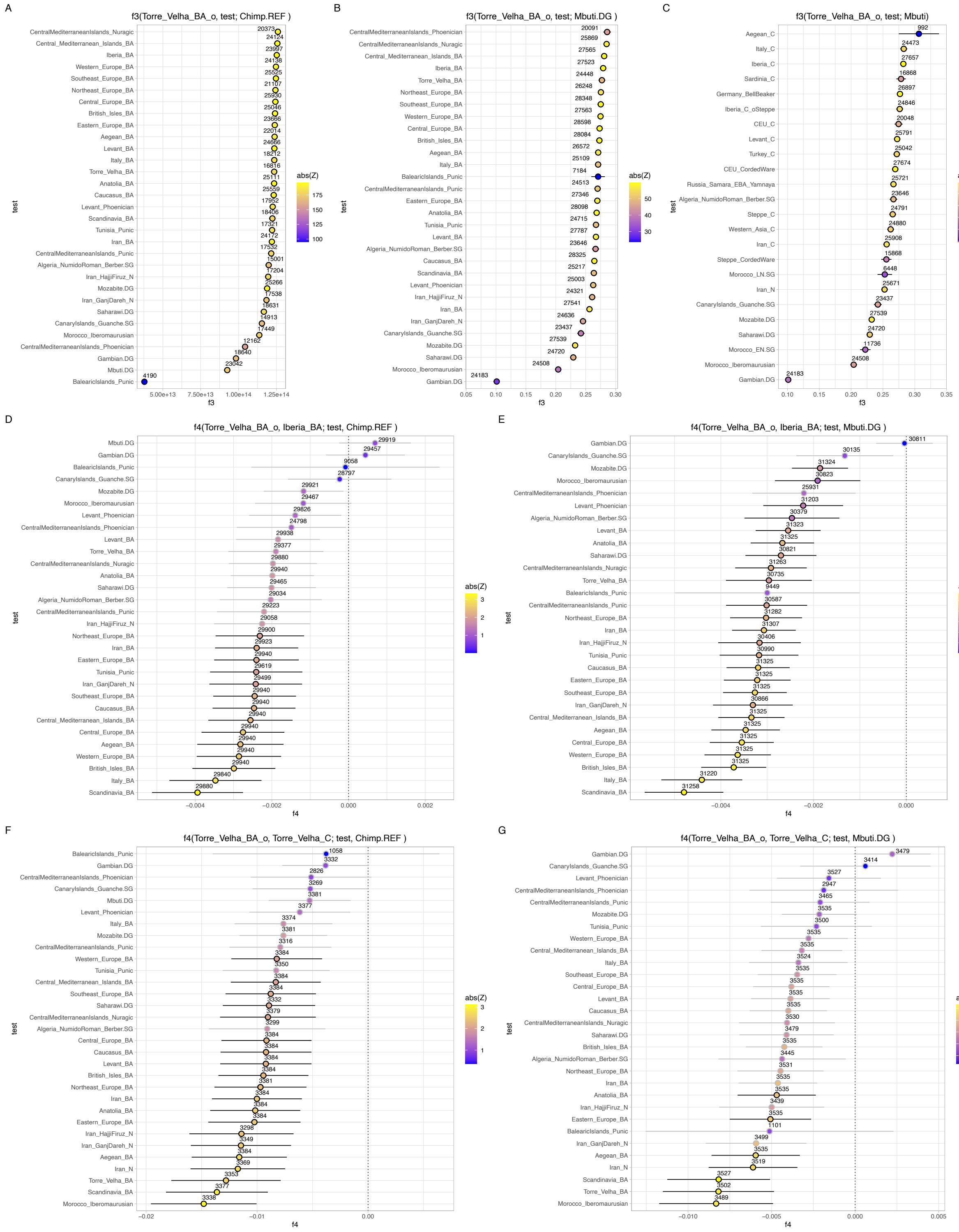

Figure S16. The x-axes represent the $f_3$- and $f_4$-statistic values, with results displayed as the mean ± 1-SD, colors representing Z-scores and black strokes Z-scores > 2. The numbers above each dot indicate the number of SNPs used for each calculation. Outgroup $f_3$-statistics are presented as: (A) $f_3$ (Torre_Velha_BA_o, Test; Chimp) and (B) $f_3$ (Torre_Velha_BA_o, Test; Mbuti), with *Test* including Eurasian and African populations from the Bronze Age or proxies; and (C) $f_3$(Torre_Velha_BA_o, Test; Mbuti), with *Test* including Eurasian and African populations from the Chalcolithic or proxies. $f_4$-statistics are shown as: (D) $f_4$(Torre_Velha_BA_o, Iberia_BA; Test, Chimp) and (E) $f_4$(Torre_Velha_BA_o, Iberia_BA;

Test, Mbuti), and (F) $f_4$(Torre_Velha_BA_o, Torre_Velha_C; Test, Chimp) and (G) $f_4$(Torre_Velha_BA_o, Torre_Velha_C; Test, Mbuti), with *Test* including Eurasian and African populations from the Bronze Age or proxies.

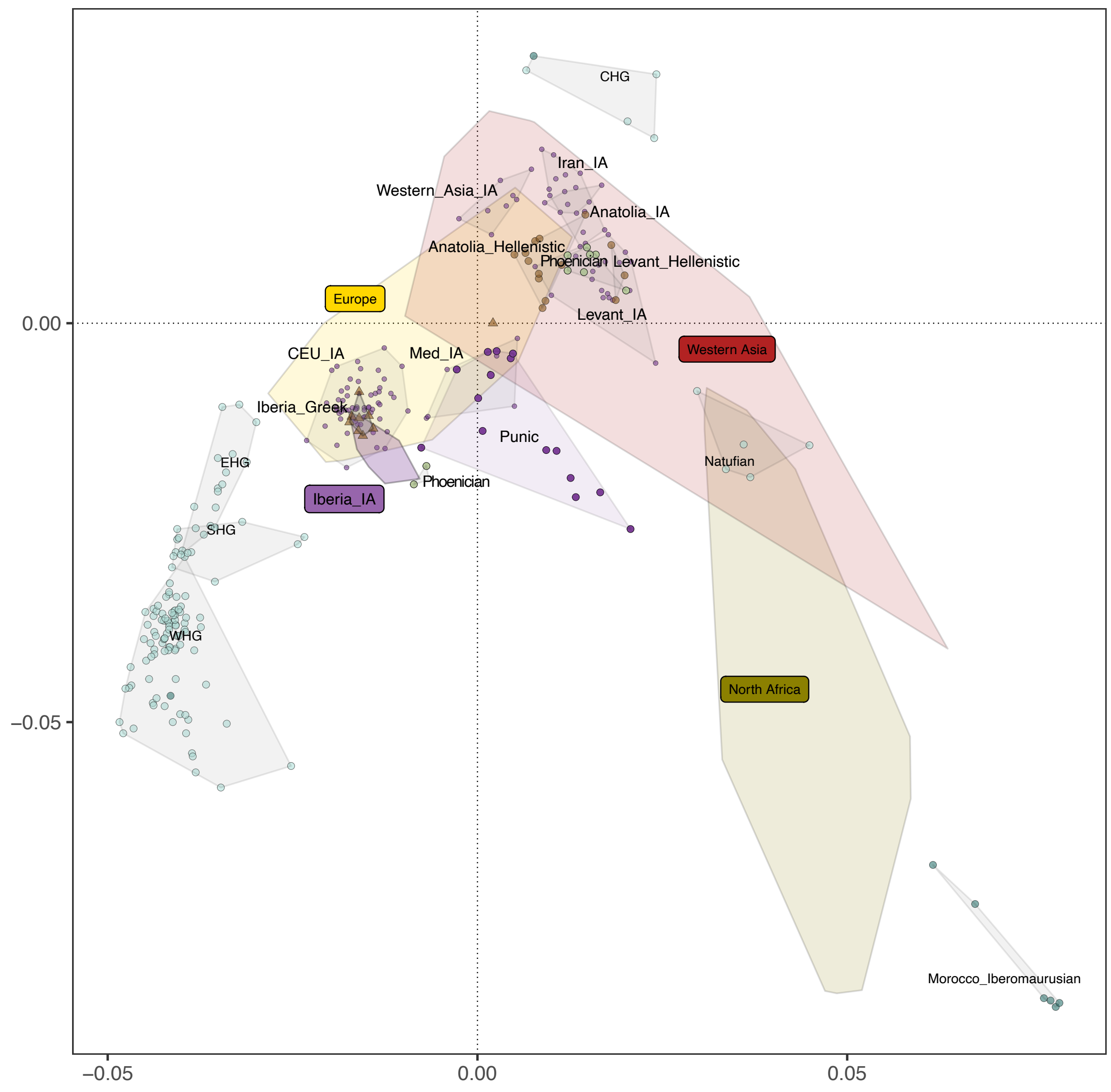

Figure S17. PCA of present-day West Eurasians and North Africans (overlaid colored polygons represent geographical clusters) with ancient individuals from Iberia and other regions projected onto the first two principal components, focusing on the Iron Age. Colors correspond to different temporal periods, as shown in 1B as well as Punic populations in purple and Phoenician in light green.

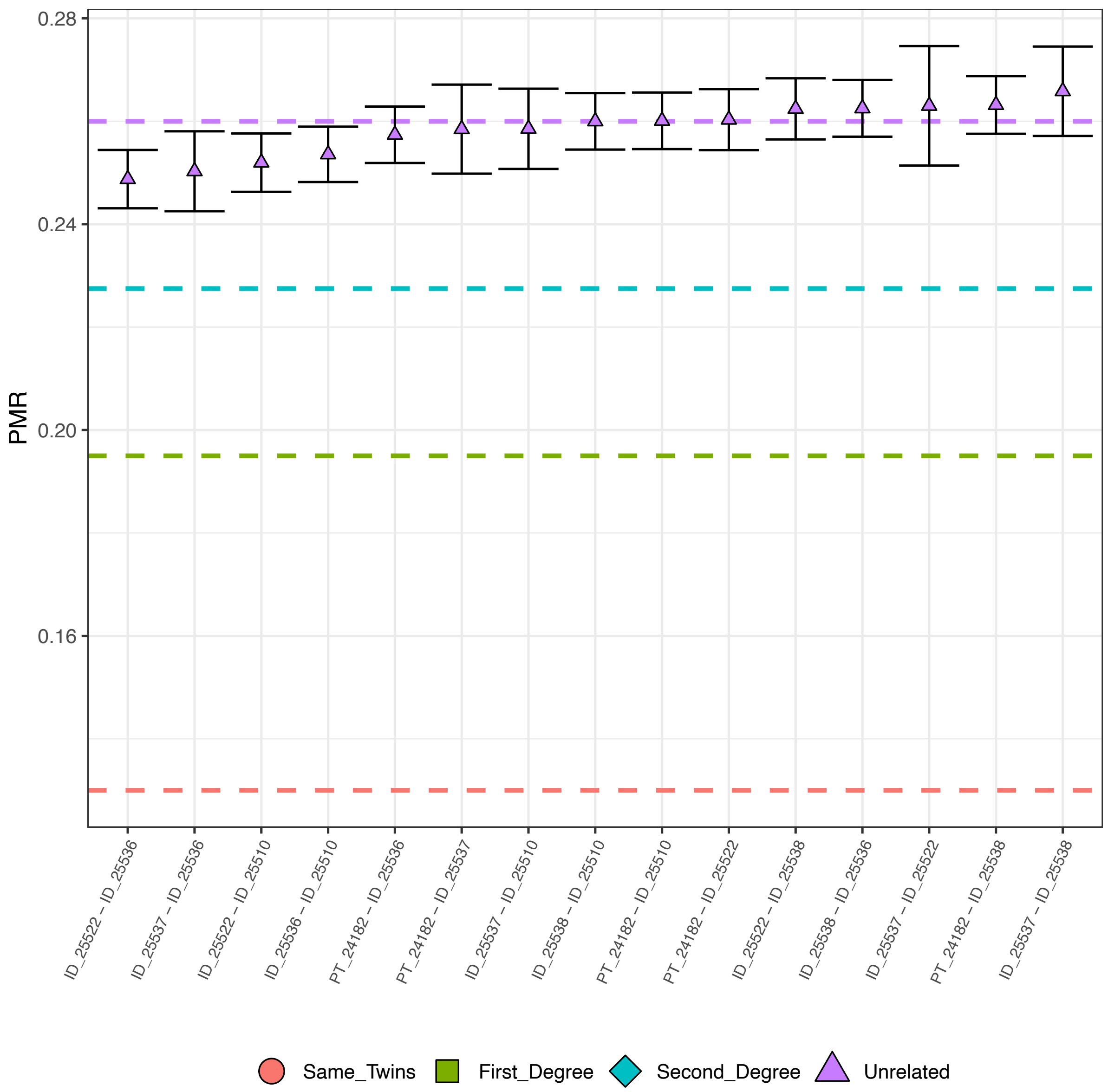

Figure S18. Kinship analysis from Idanha_a_Velha_Roman/EarlyMedieval using BREADR.

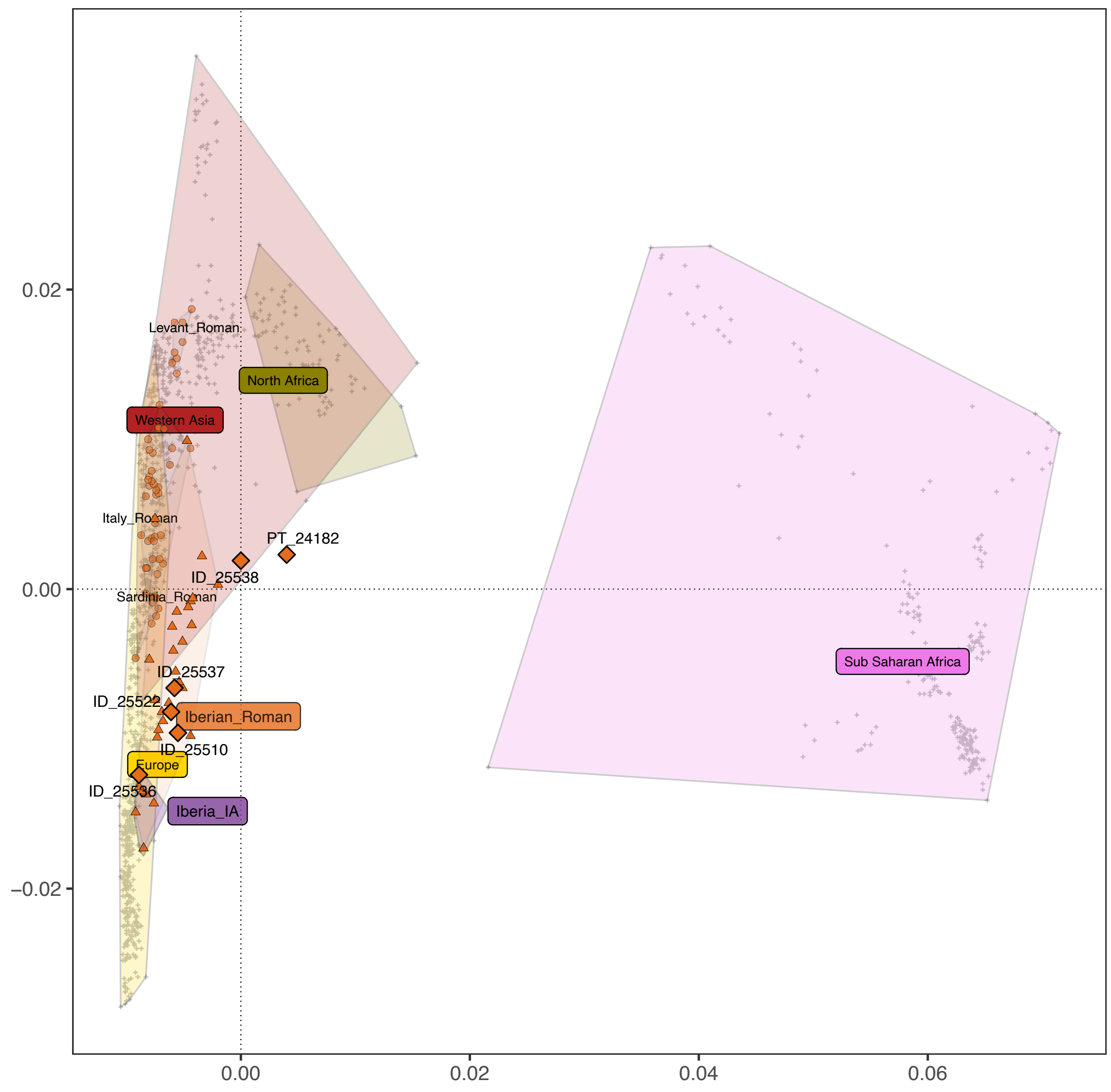

Figure S19. PCA of present-day West Eurasians, North Africans and Sub-Saharan Africans (overlaid colored polygons represent geographical clusters) with ancient individuals from Iberia and other regions projected onto the first two principal components, focusing on the Roman period. Colors correspond to different temporal periods, as shown in 1B.

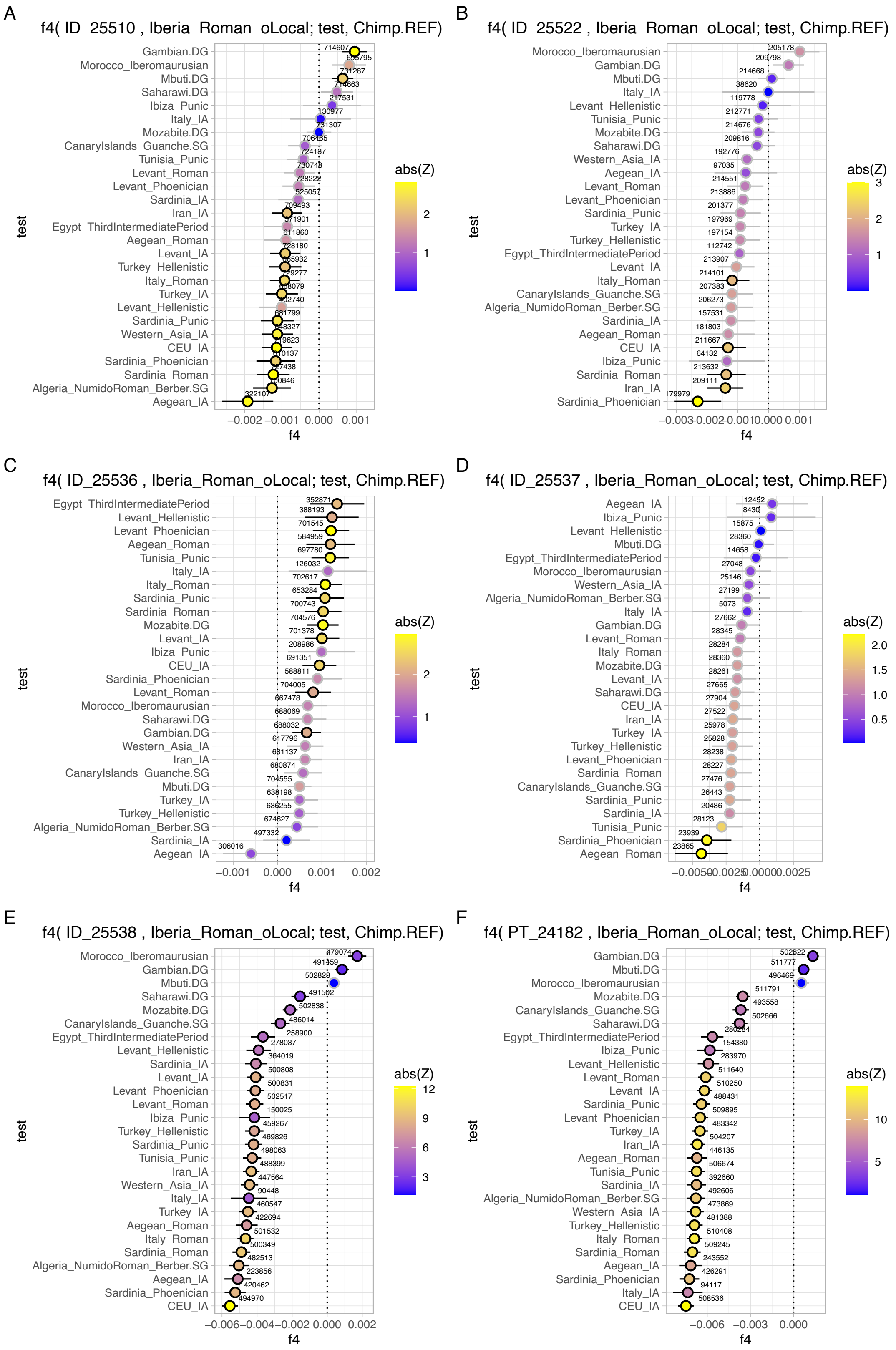

Figure S20. The x-axes represent the $f_4$-statistic values, with results displayed as the mean ± 1-SD, colors representing Z-scores and black strokes Z-scores > 2. The numbers above each dot indicate the number of SNPs used for each calculation. $f_4$-statistics are shown as $f_4$(X, Iberia_Roman_oLocal; Test, Chimp) with *Test* including Eurasian and African populations from the Roman period or proxies and *X* being (A) ID_25510, (B) ID_25522, (C) ID_25536, (D) ID_25537, (E) ID_25538 and (F) PT_24182.

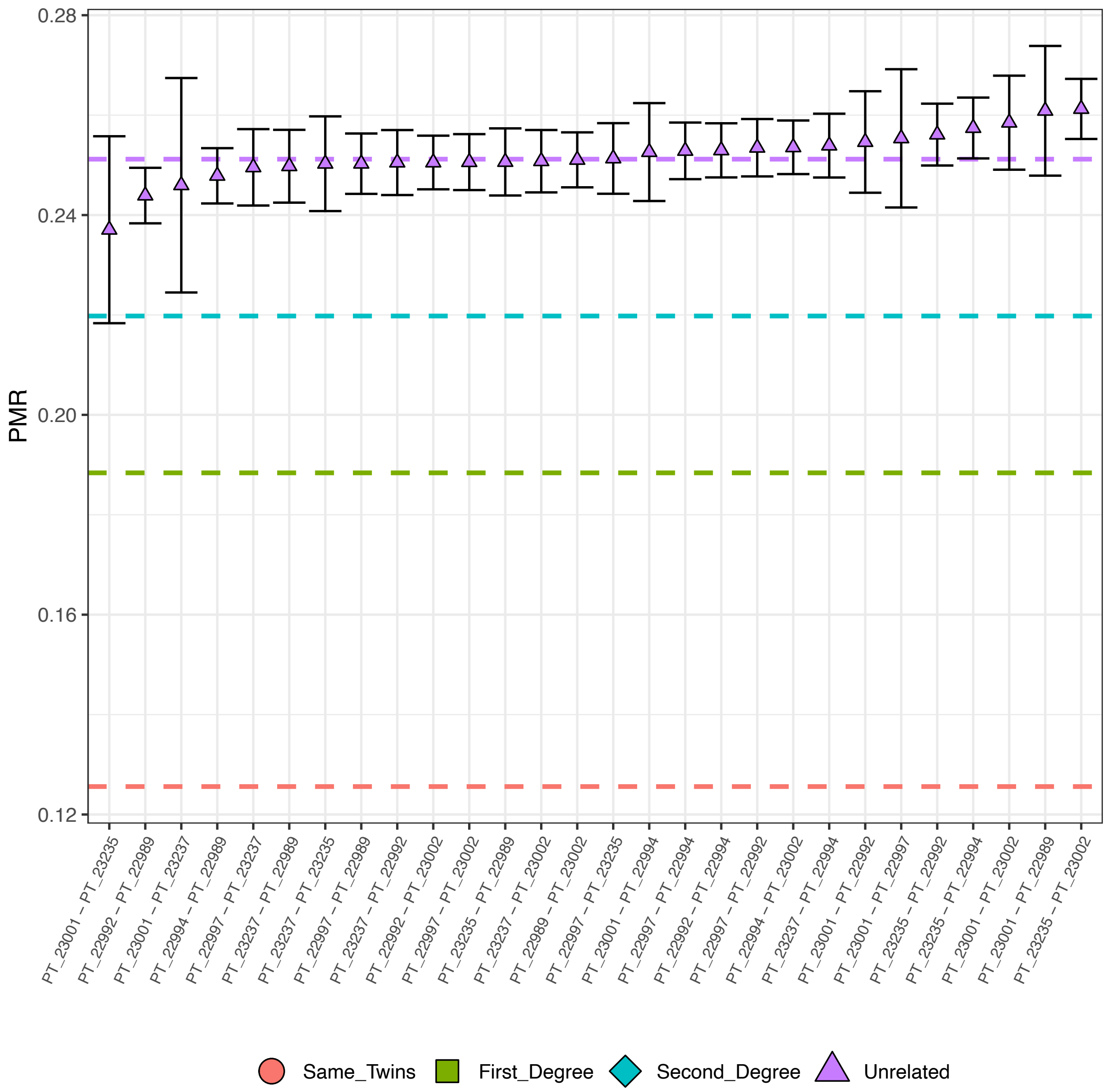

Figure S21. Kinship analysis from Guarda_Prazo_Freixo_de_Numão (EarlyMedieval and Conquest) and Castro_de_Avelãs_Torre_Velha (EarlyMedieval and Conquest) using BREADR.

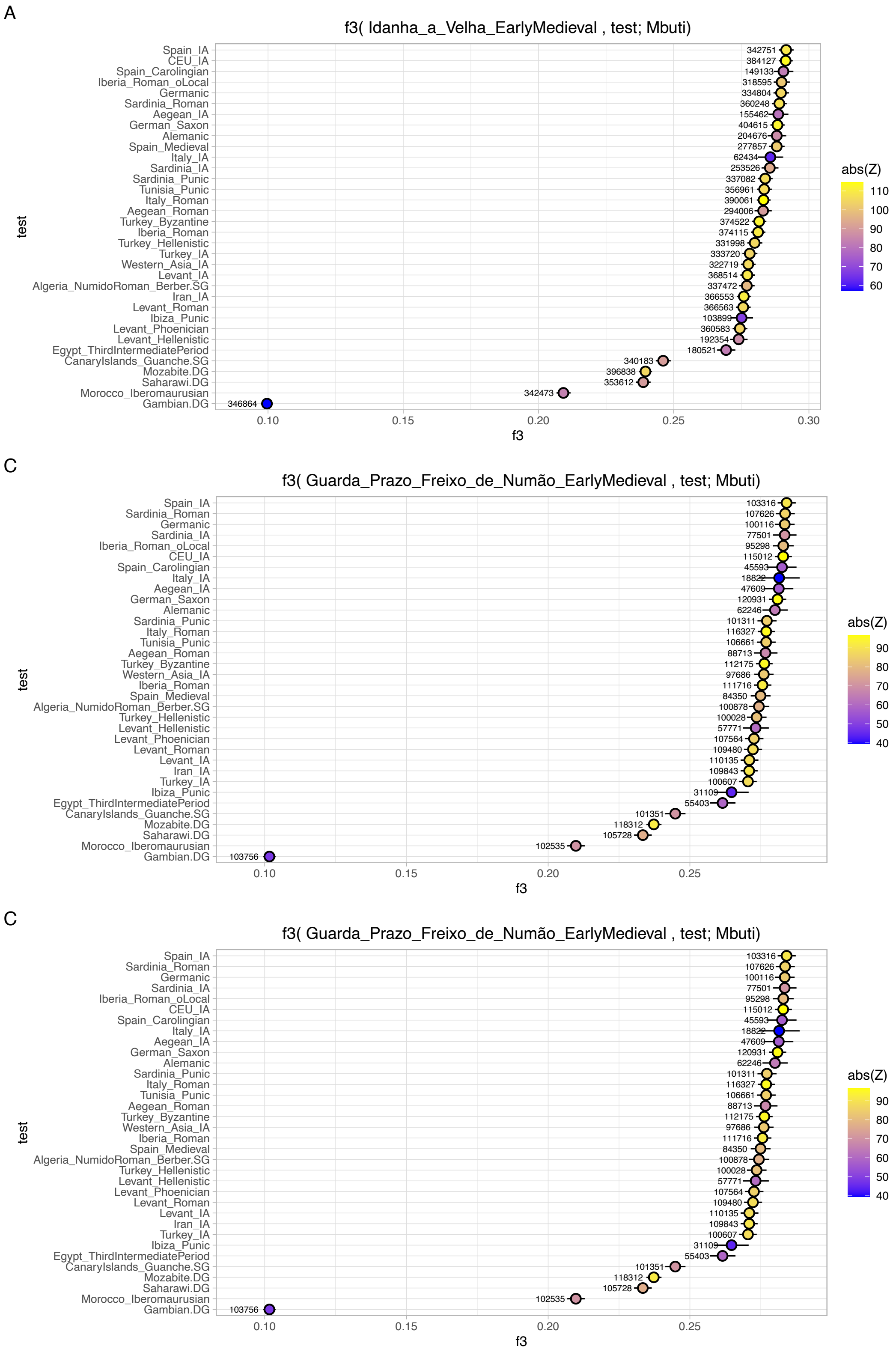

Figure S22. The x-axes represent the $f_3$-statistic values, with results displayed as the mean ± 1-SD, and colors representing Z-scores. The numbers above each dot indicate the number of SNPs used for each calculation. Outgroup $f_3$-statistics are presented as $f_3$(X, Test; Chimp) with *Test* including Eurasian and African populations from the Early Medieval period or proxies and *X* being (A) Idanha_a_Velha_EarlyMedieval, (B) Guarda_Prazo_Freixo_de_Numão_EarlyMedieval and (C) Castro_de_Avelãs_Torre_Velha_EarlyMedieval.

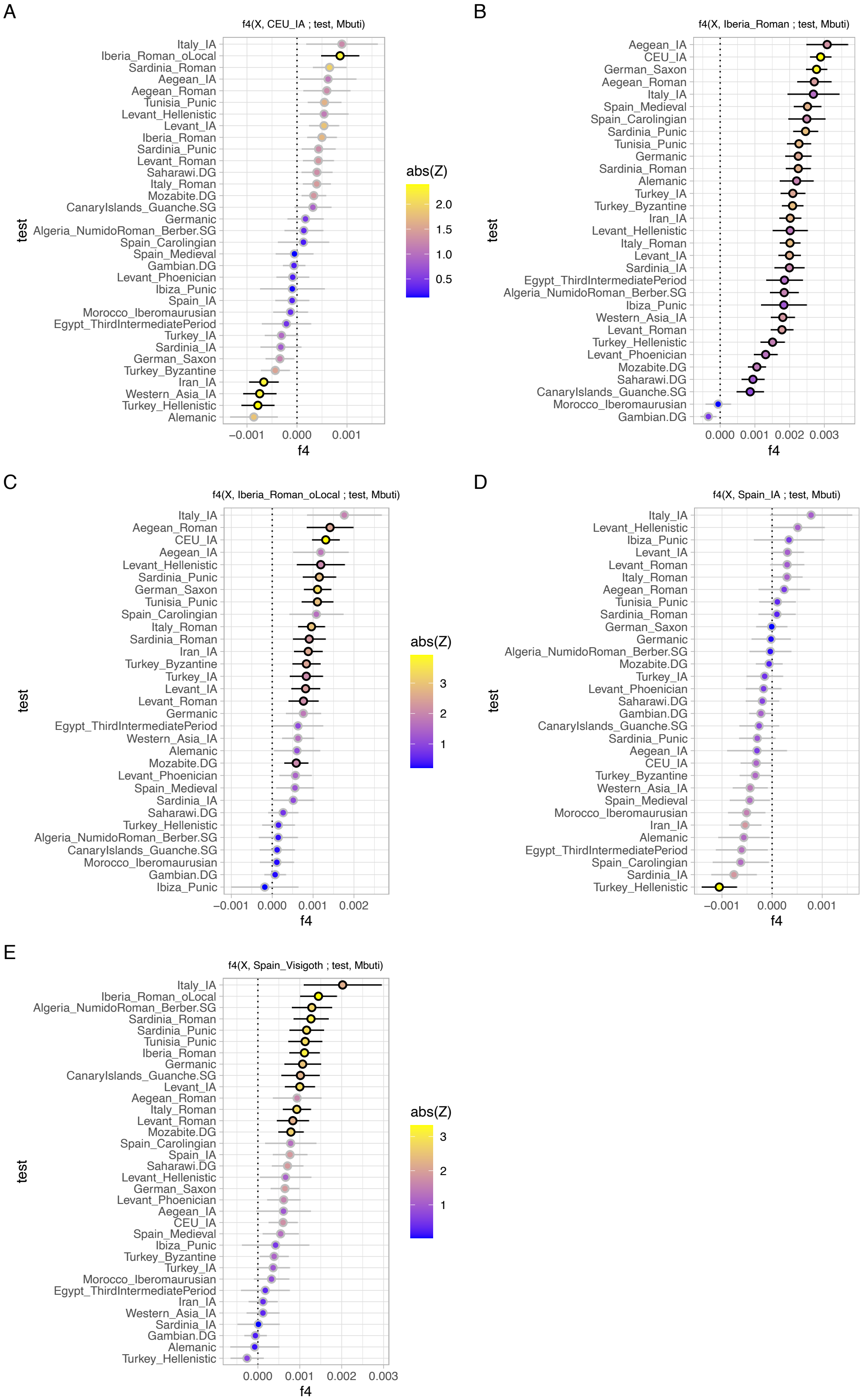

Figure S23. The x-axes represent the $f_4$-statistic values, with results displayed as the mean ± 1-SD, colors representing Z-scores and black strokes Z-scores > 2. $f_4$-statistics are shown as $f_4$(Idanha_a_Velha_EarlyMedieval, X; Test, Mbuti) with *Test* including Eurasian and African populations from the Visigoth period or proxies and *X* being (A) CEU_IA, (B) Iberia_Roman, (C) Iberia_Roman_oLocal, (D) Spain_IA, (E) Spain_Visigoth.

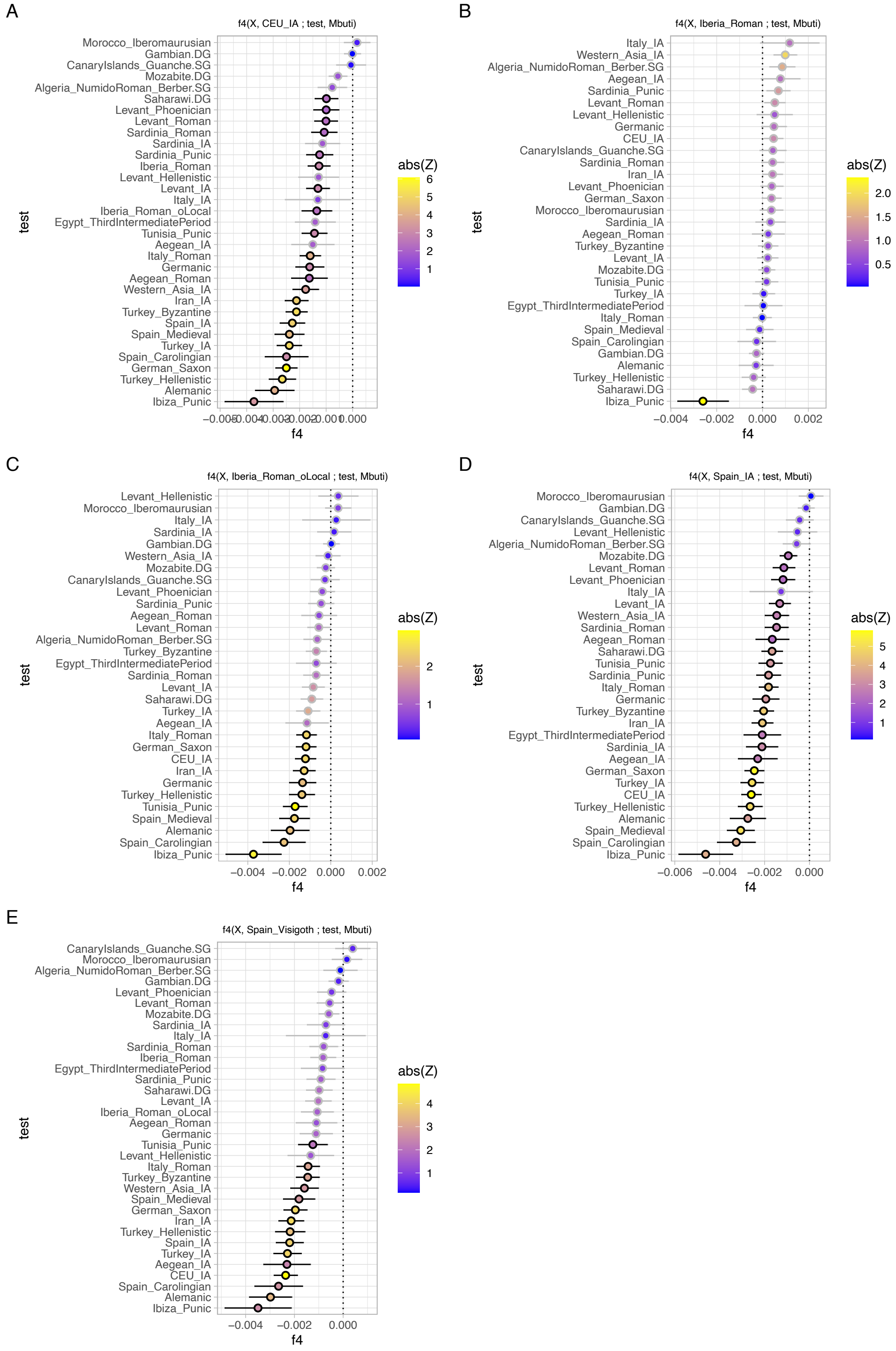

Figure S24. The x-axes represent the $f_4$-statistic values, with results displayed as the mean ± 1-SD, colors representing Z-scores and black strokes Z-scores > 2. $f_4$-statistics are shown as $f_4$(X, Y; Test, Mbuti) with *X* being Guarda_Prazo_Freixo_de_Numão_EarlyMedieval, *Test* including Eurasian and African populations from the Visigoth period or proxies and *Y* being (A) CEU_IA, (B) Iberia_Roman, (C) Iberia_Roman_oLocal, (D) Spain_IA, (E) Spain_Visigoth.

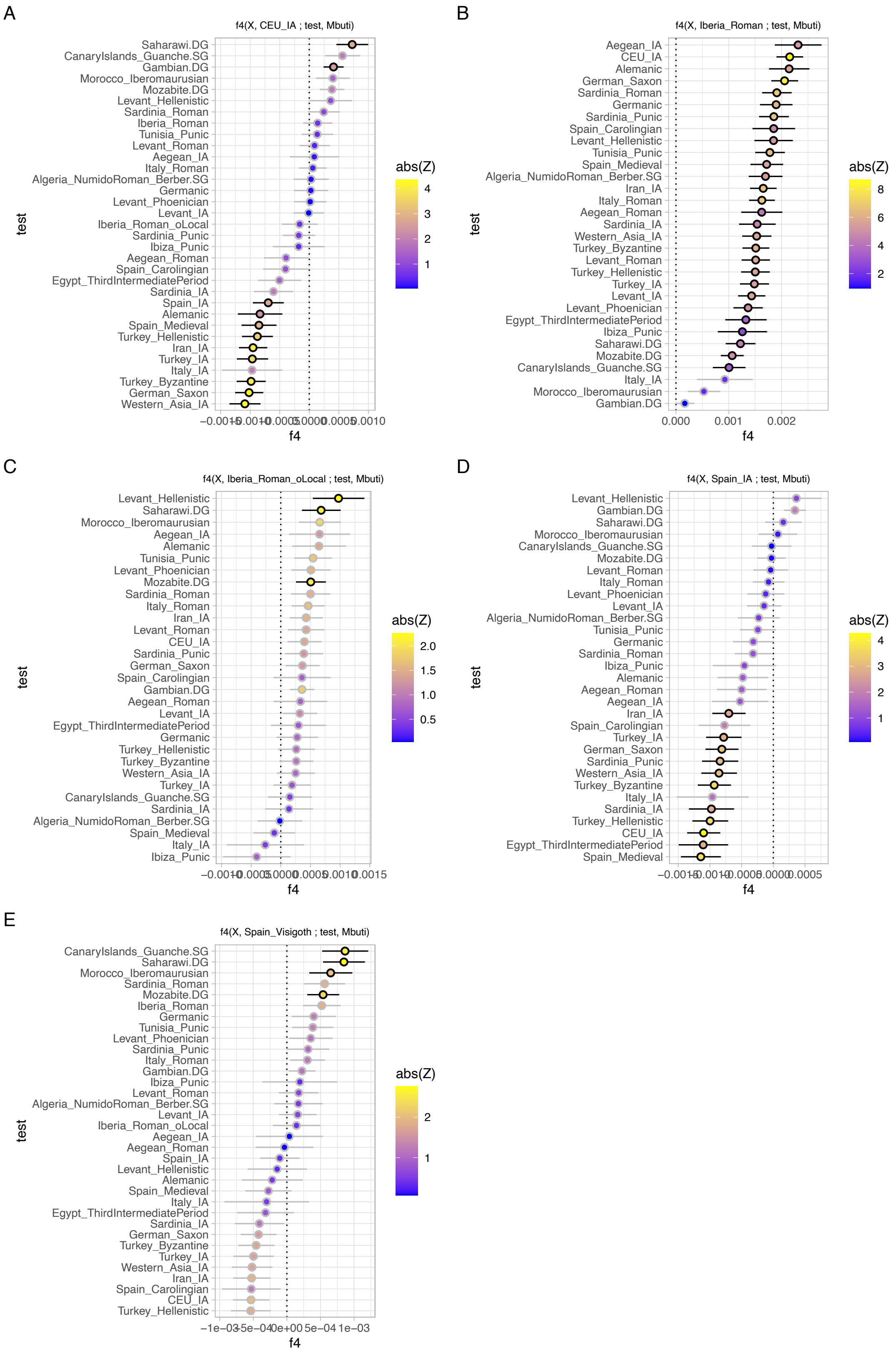

Figure S25. The x-axes represent the $f_4$-statistic values, with results displayed as the mean ± 1-SD, colors representing Z-scores and black strokes Z-scores > 2. $f_4$-statistics are shown as $f_4$(X, Y; Test, Mbuti) with *X* being Castro_de_Avelãs_Torre_Velha_EarlyMedieval, *Test* including Eurasian and African populations from the Visigoth period or proxies and *Y* being (A) CEU_IA, (B) Iberia_Roman, (C) Iberia_Roman_oLocal, (D) Spain_IA, (E) Spain_Visigoth.

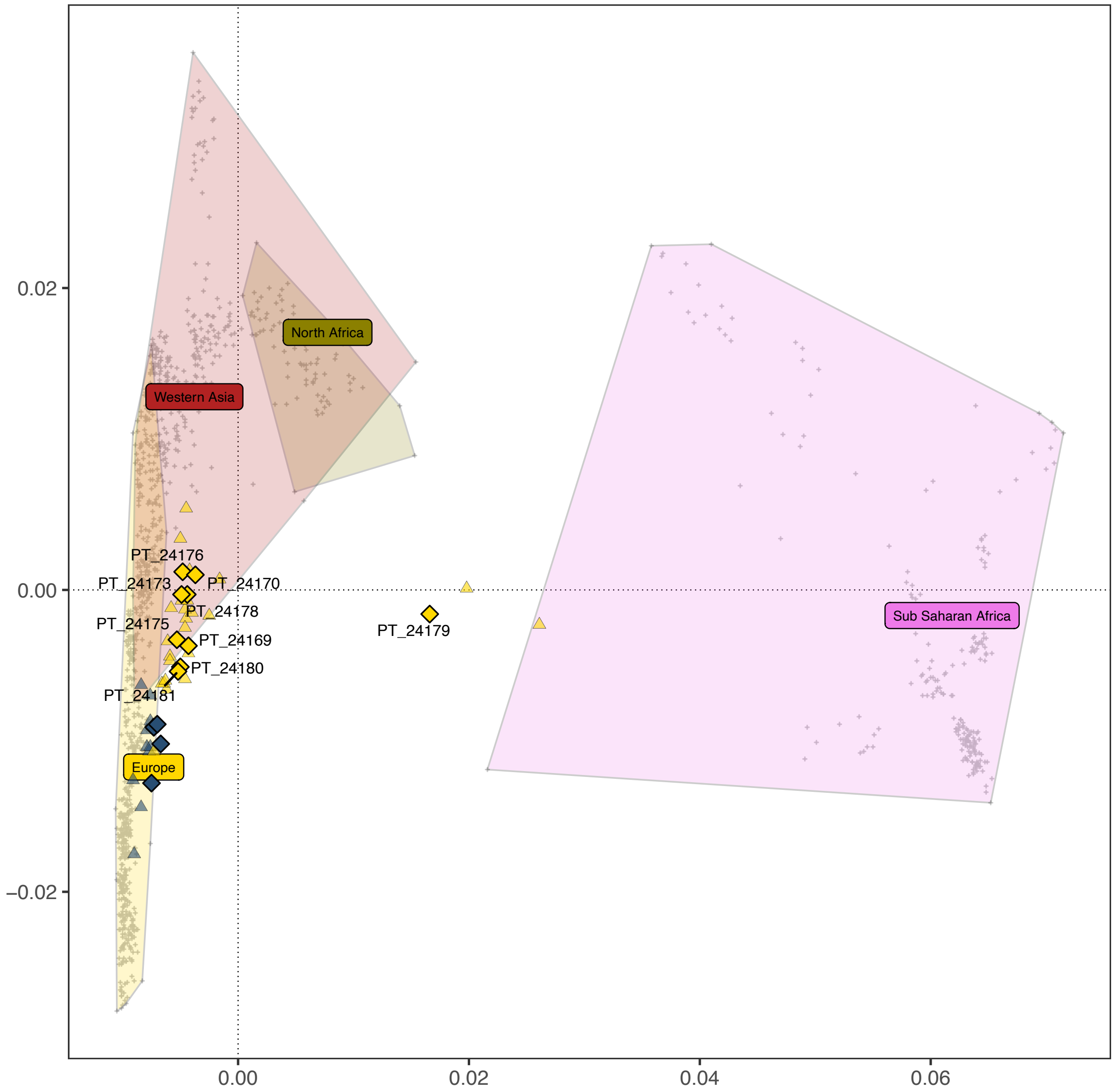

Figure S26. PCA of present-day West Eurasians, North Africans and Sub-Saharan Africans (overlaid colored polygons represent geographical clusters) with ancient individuals from Iberia and other regions projected onto the first two principal components, focusing on the Islamic period. Colors correspond to different temporal periods, as shown in 1B.

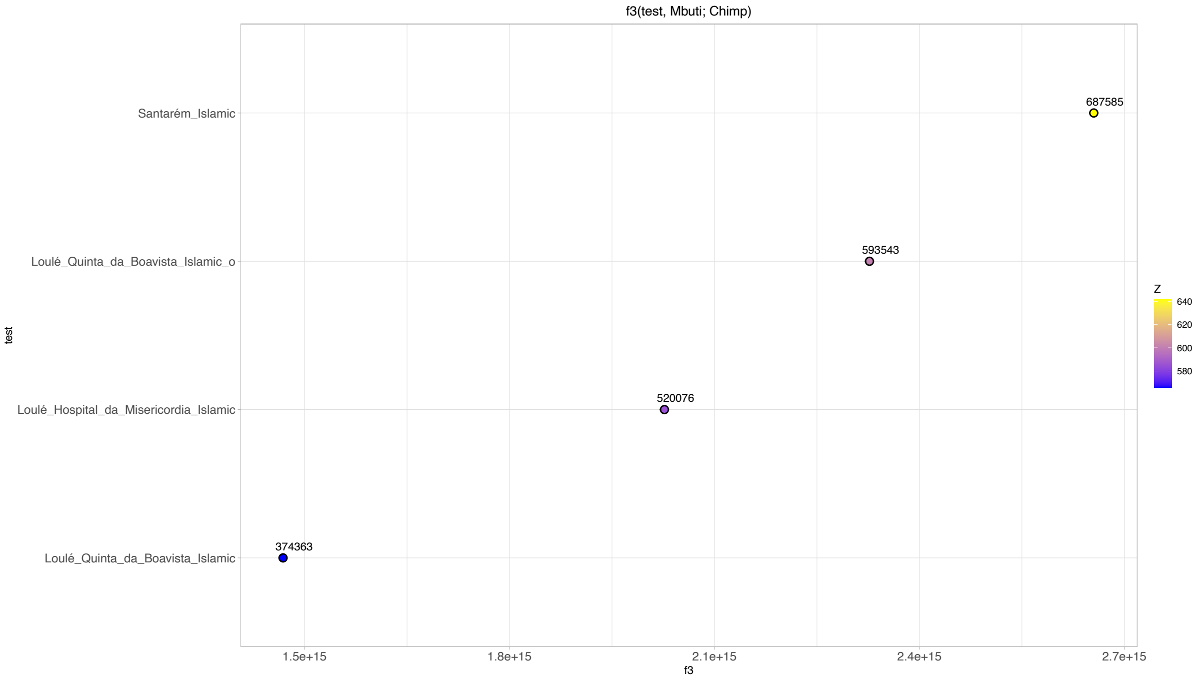

Figure S27. The x-axes represent the ff-statistic values, with results displayed as the mean ± 1-SD, and colors representing Z-scores. The numbers above each dot indicate the number of SNPs used for each calculation. Outgroup $f_3$-statistics are presented as $f_3$(Test, Mbuti; Chimp) with *Test* being Santarém_Islamic and Loulé_Islamic.

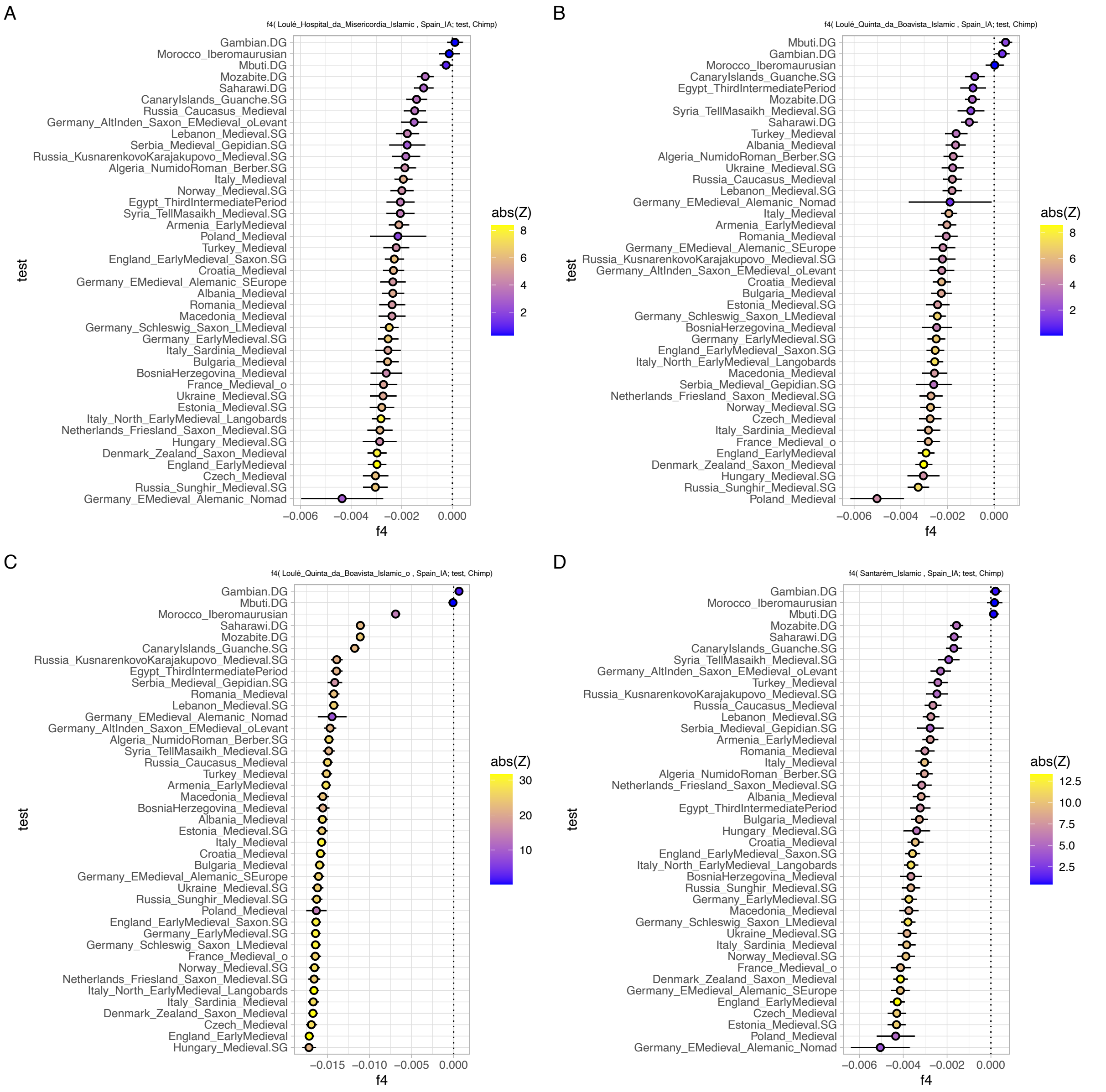

Figure S28. The x-axes represent the $f_4$-statistic values, with results displayed as the mean ± 1-SD, and colors representing Z-scores. The numbers above each dot indicate the number of SNPs used for each calculation. $f_4$-statistics are shown as $f_4$(X, Spain_IA; Test, Chimp) with *Test* including Eurasian and African populations from the Islamic period or proxies and *X* being (A) Loulé_Hospital_da_Misericordia_Islamic, (B) Loulé_Quinta_da_Boavista_Islamic, (C) Loulé_Quinta_da_Boavista_Islamic_o and (D) Spain_Visigoth.

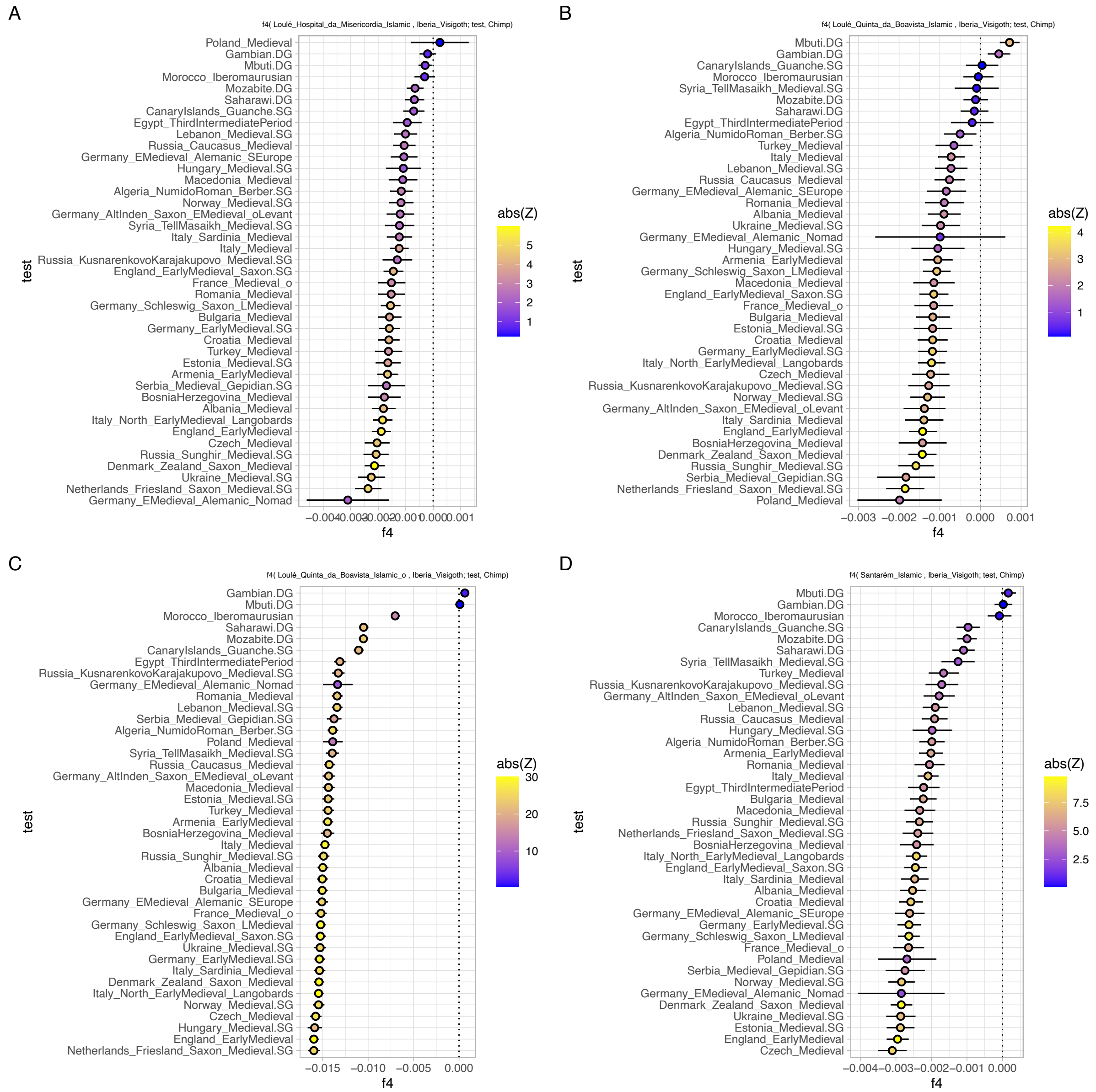

Figure S29. The x-axes represent the $f_4$-statistic values, with results displayed as the mean ± 1-SD, and colors representing Z-scores. The numbers above each dot indicate the number of SNPs used for each calculation. $f_4$-statistics are shown as $f_4$(X, Iberia_Visigoth; Test, Chimp) with *Test* including Eurasian and African populations from the Islamic period or proxies and *X* being (A) Loulé_Hospital_da_Misericordia_Islamic, (B) Loulé_Quinta_da_Boavista_Islamic, (C) Loulé_Quinta_da_Boavista_Islamic_o and (D) Spain_Visigoth.

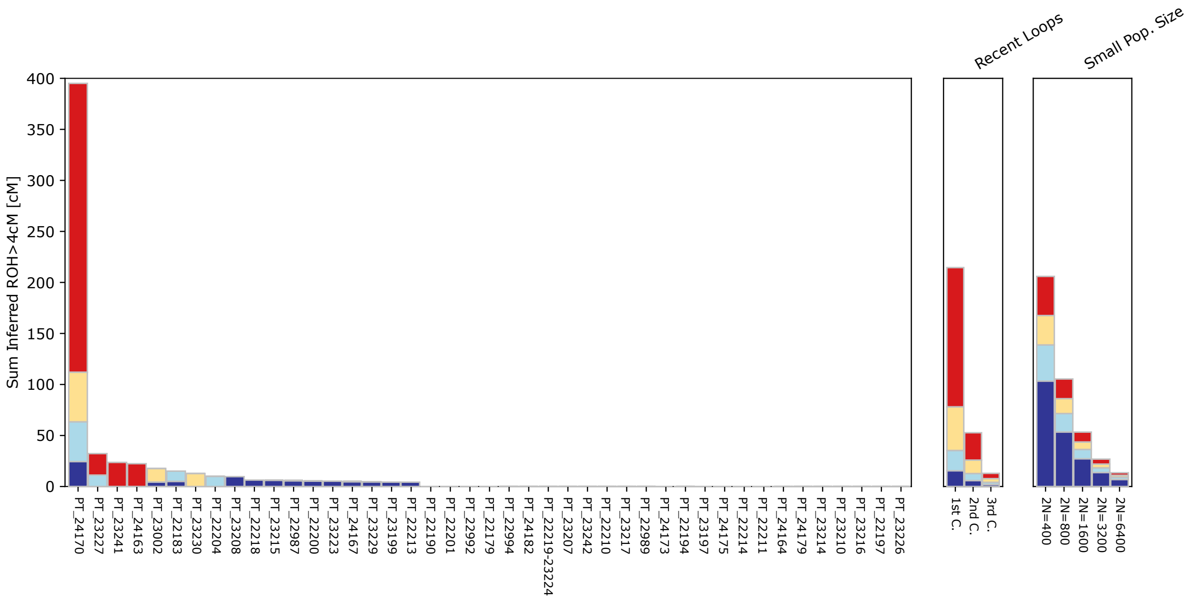

Figure S30. Runs of Homozygosity (ROH).

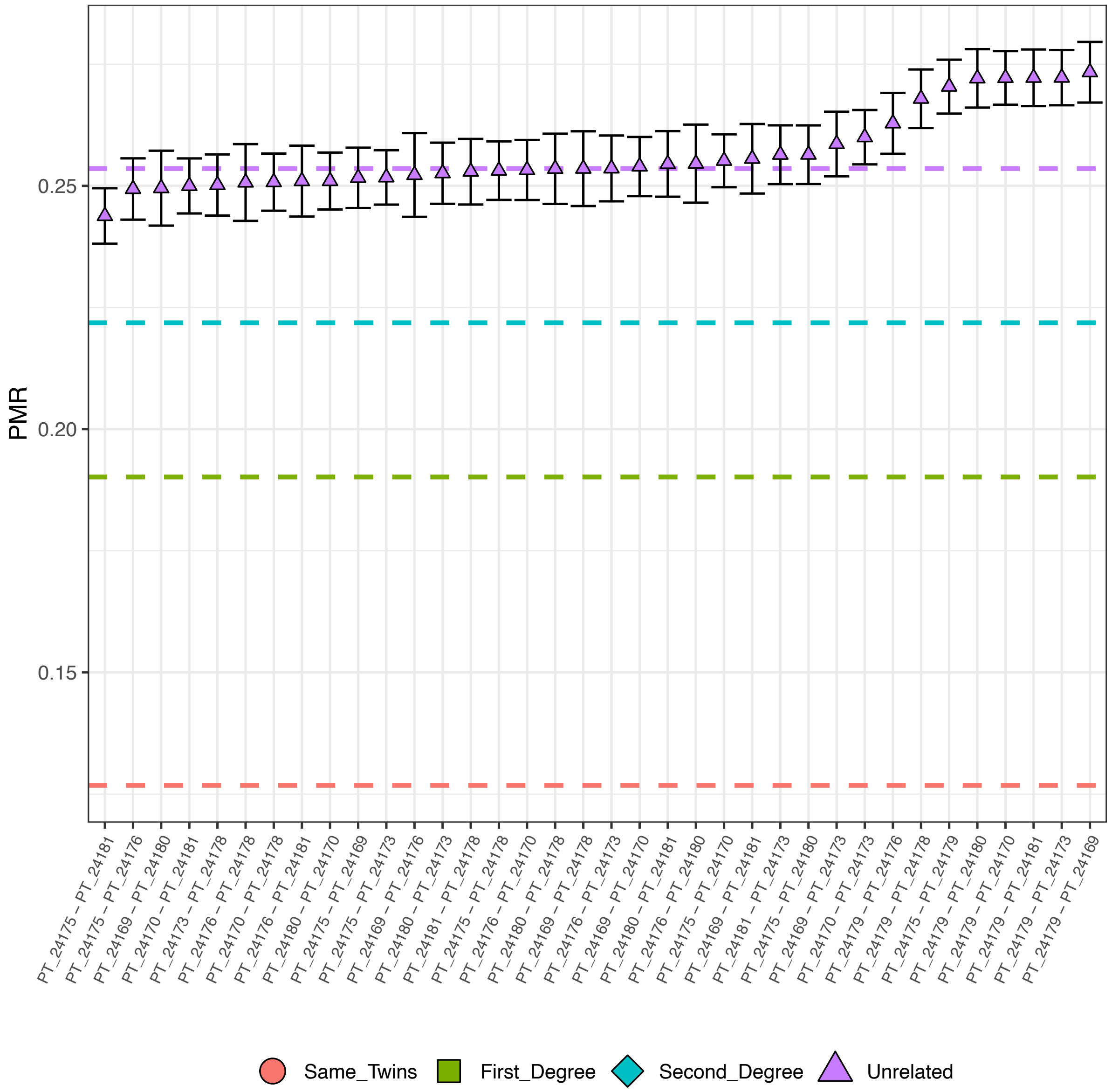

Figure S31. Kinship analysis from Loulé_Islamic and Santarém_Islamic using BREADR.

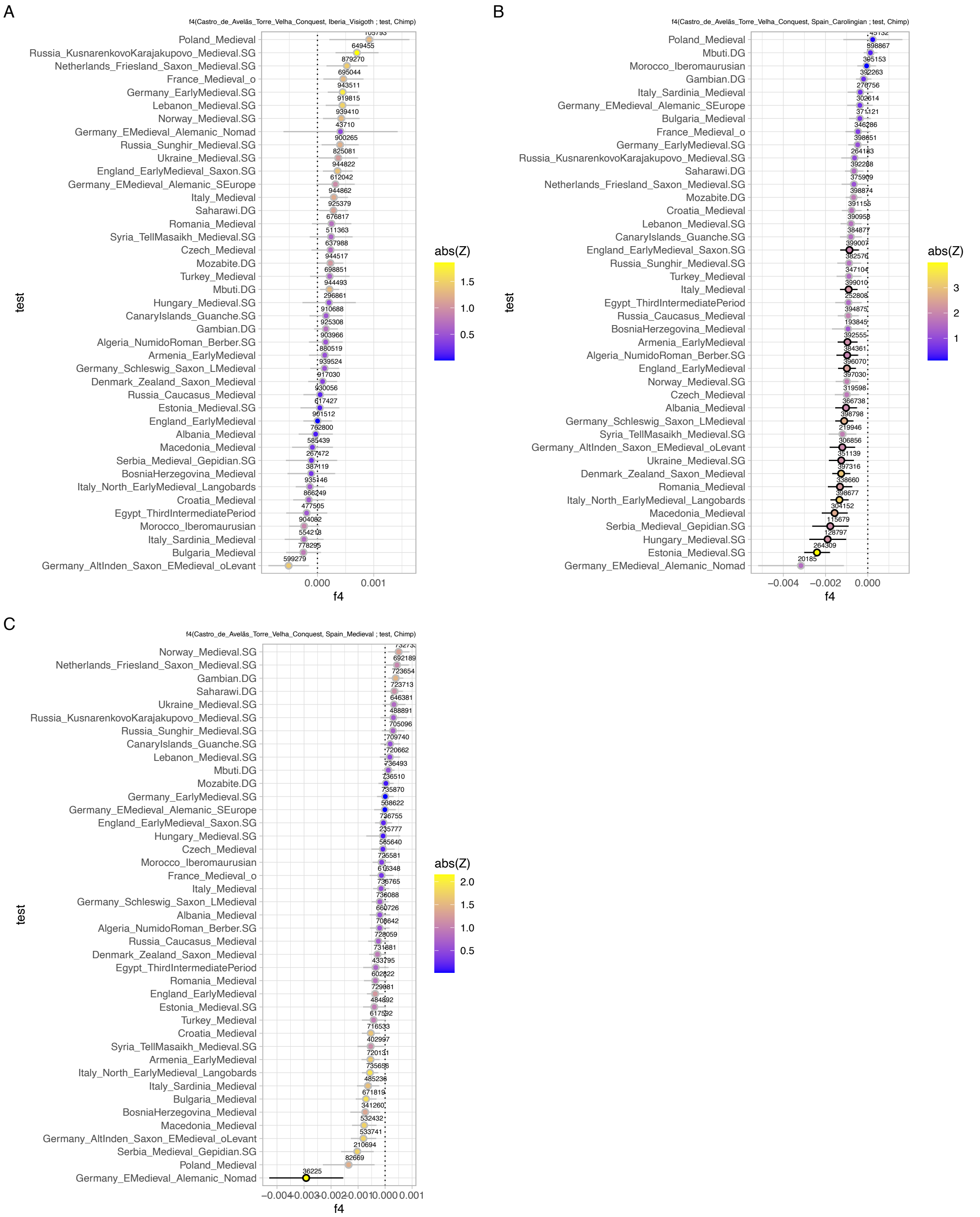

Figure S32. The x-axes display $f_4$-statistic values as mean ± 1 SD, with colors indicating Z-scores and black strokes for Z-scores > 2. Numbers above each dot show the number of SNPs used. $f_4$-statistics are shown as $f_4$(Castro_de_Avelãs_Torre_Velha_Conquest, X; Test, Mbuti) with *Test* including Eurasian and African populations from the Medieval period or proxies and *X* being (A) Iberia_Visigoth, (B) Spain_Carolingian, (C) Spain_Medieval.

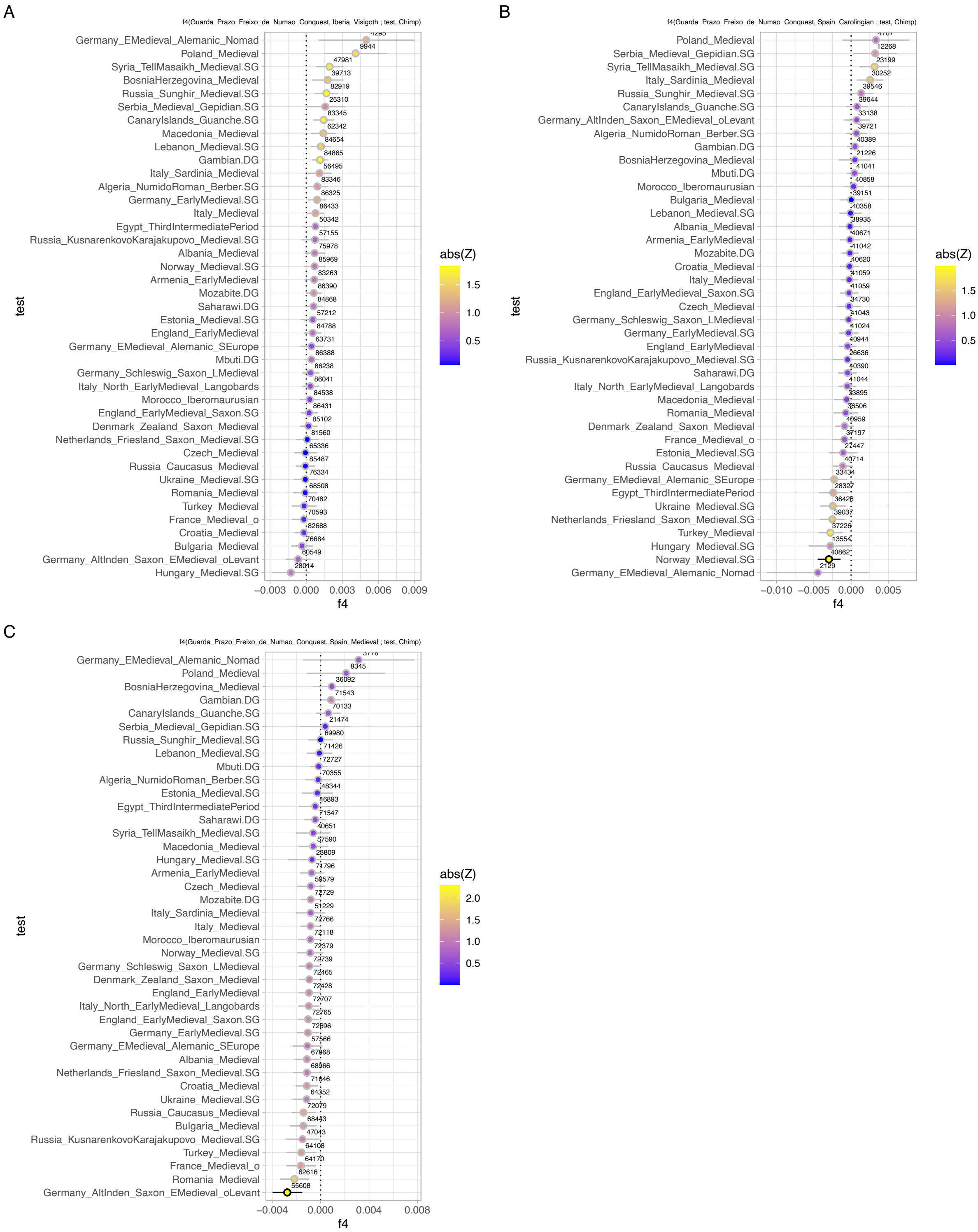

Figure S33. The x-axes represent the $f_4$-statistic values, with results displayed as the mean ± 1-SD, colors representing Z-scores and black strokes Z-scores > 2. The numbers above each dot indicate the number of SNPs used for each calculation. $f_4$-statistics are shown as $f_4$(Guarda_Prazo_Freixo_de_Numão_Conquista, X; Test, Mbuti) with *Test* including Eurasian and African populations from the Medieval period or proxies and *X* being (A) Iberia_Visigoth, (B) Spain_Carolingian, (C) Spain_Medieval.

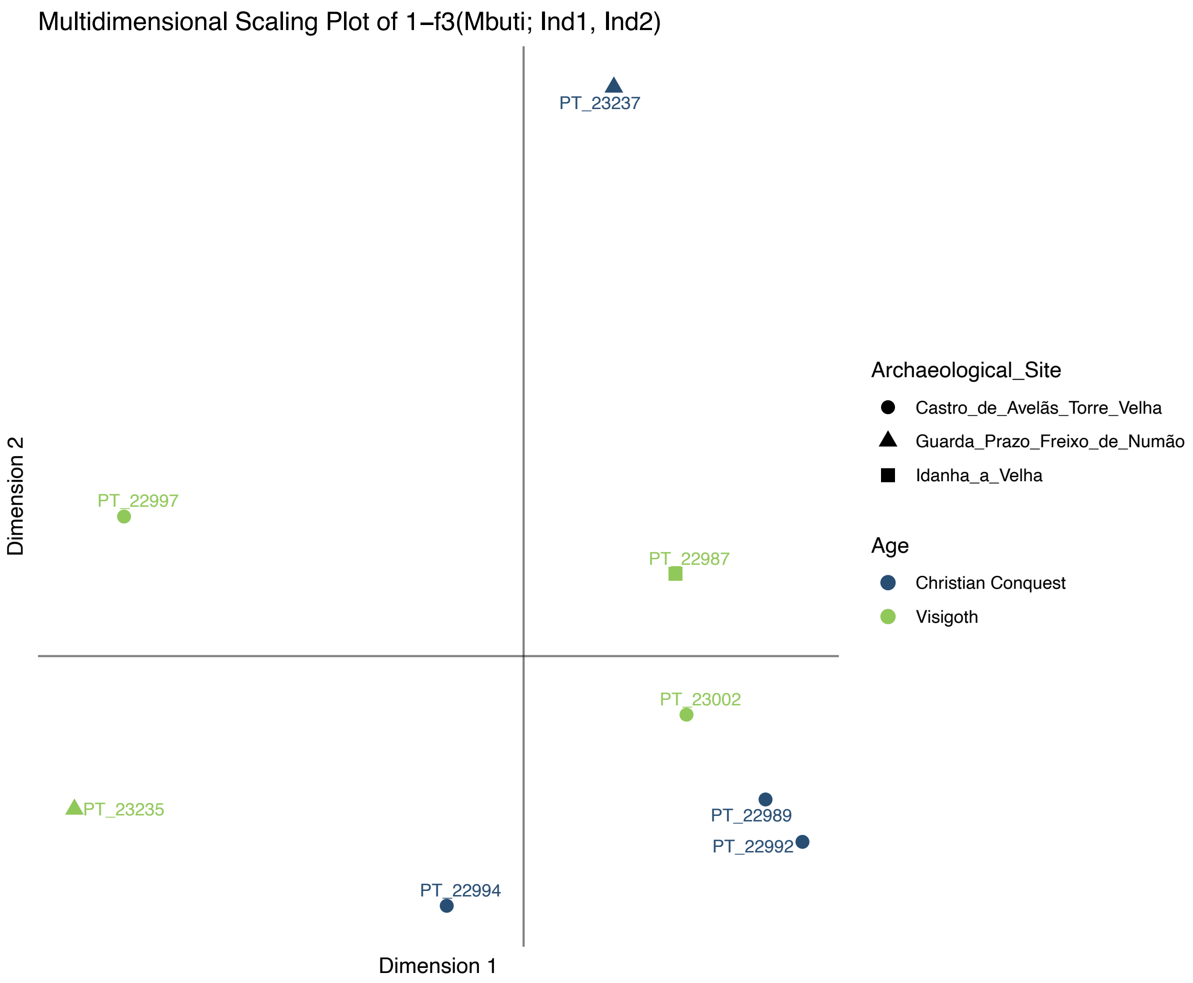

Figure S34. Multidimensional scaling (1- $f_3$) for the Visigoth and Christian Conquest individuals. Outgroup pairwise $f_3$ computed with Mbuti as outgroup.

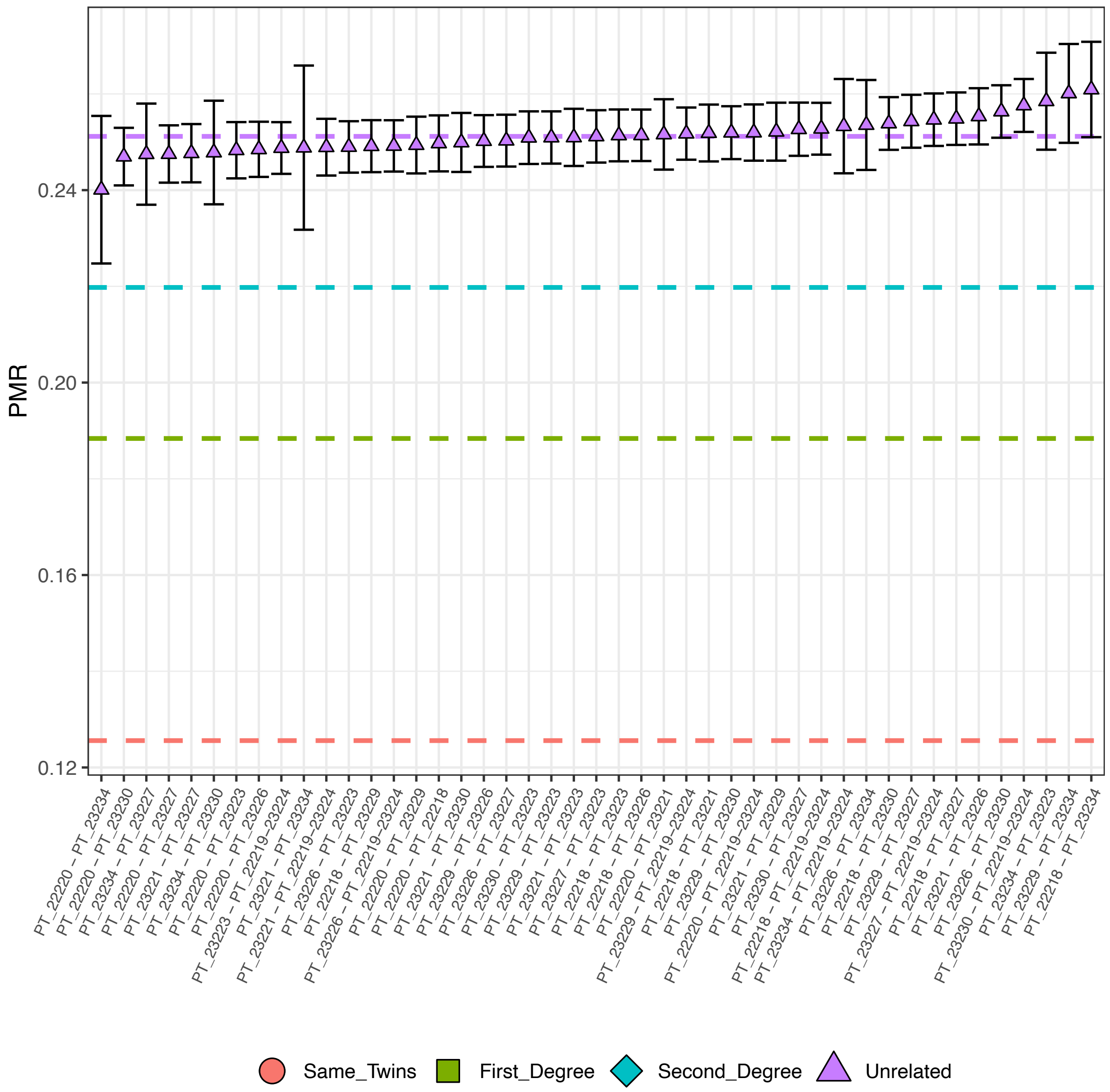

Figure S35. Kinship analysis from Santarém_Rua_dos_Barcos_13th-c using BREADR.

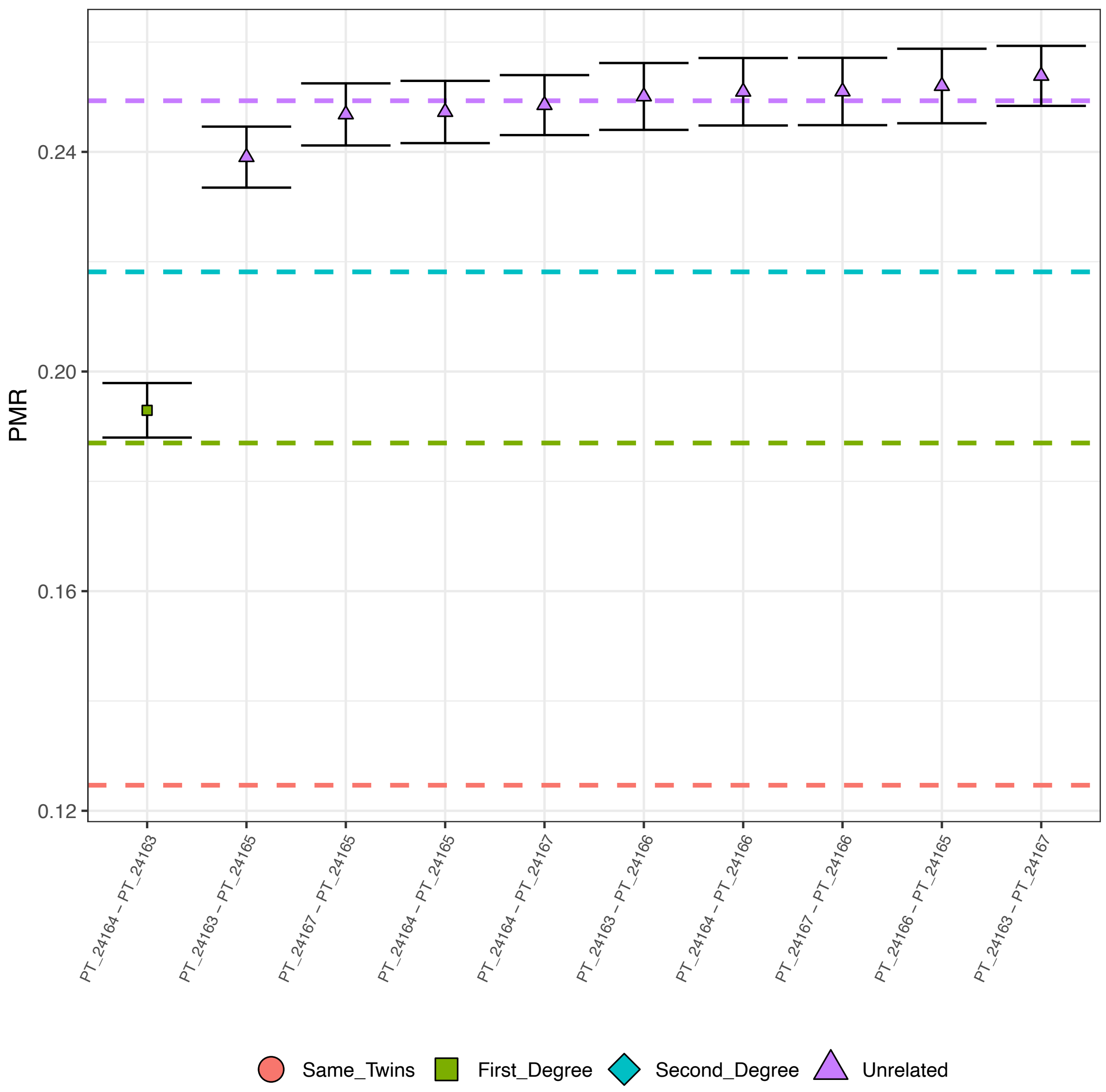

Figure S36. Kinship analysis from São_Miguel_de_Odrinhas_13th-c using BREADR.

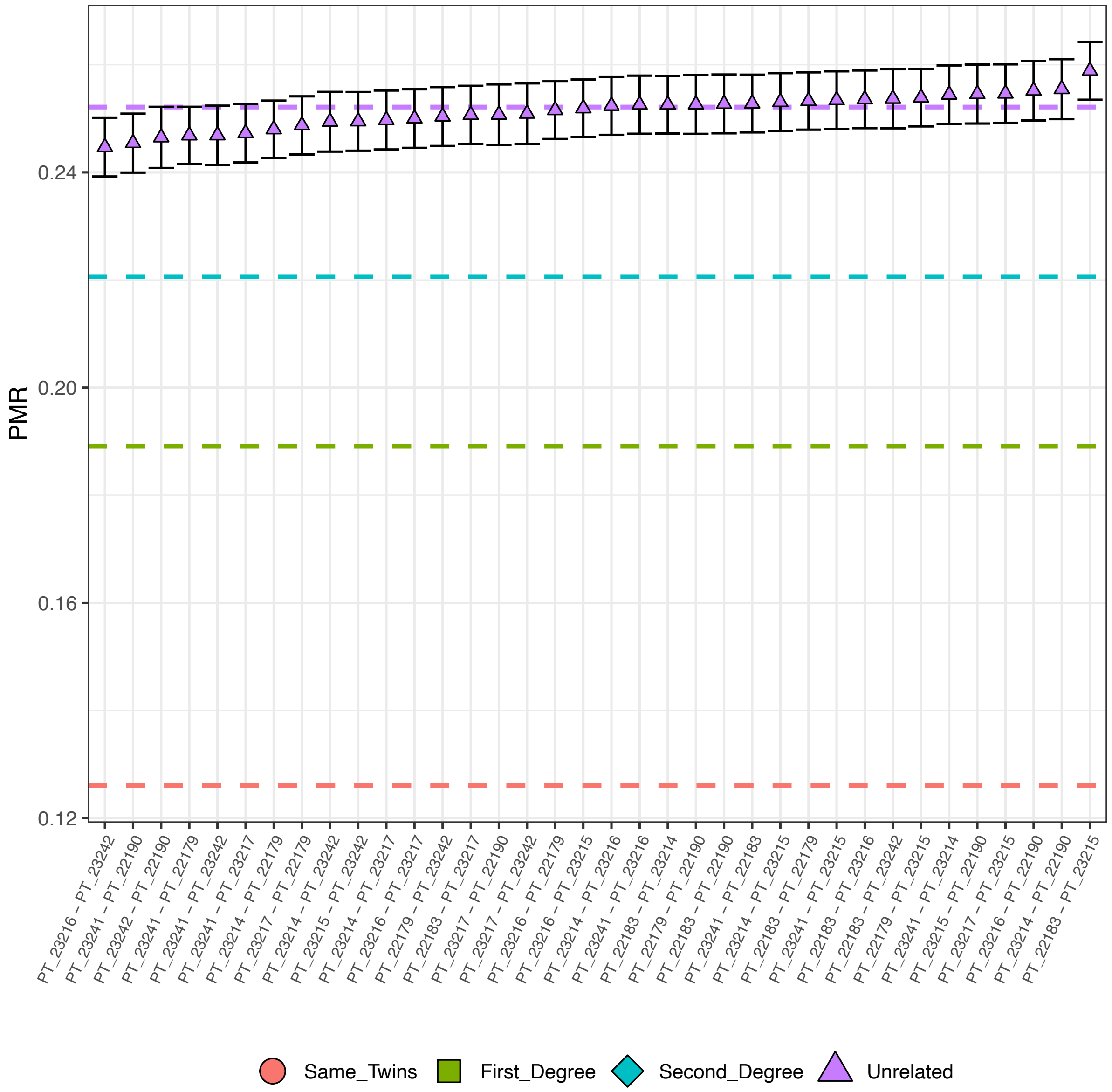

Figure S37. Kinship analysis from Castelo_de_Montemor_o_Velho_18th-c, Aveiro_Travanca_18th-c, and Castelo_de_Abrantes_19th-c using BREADR.

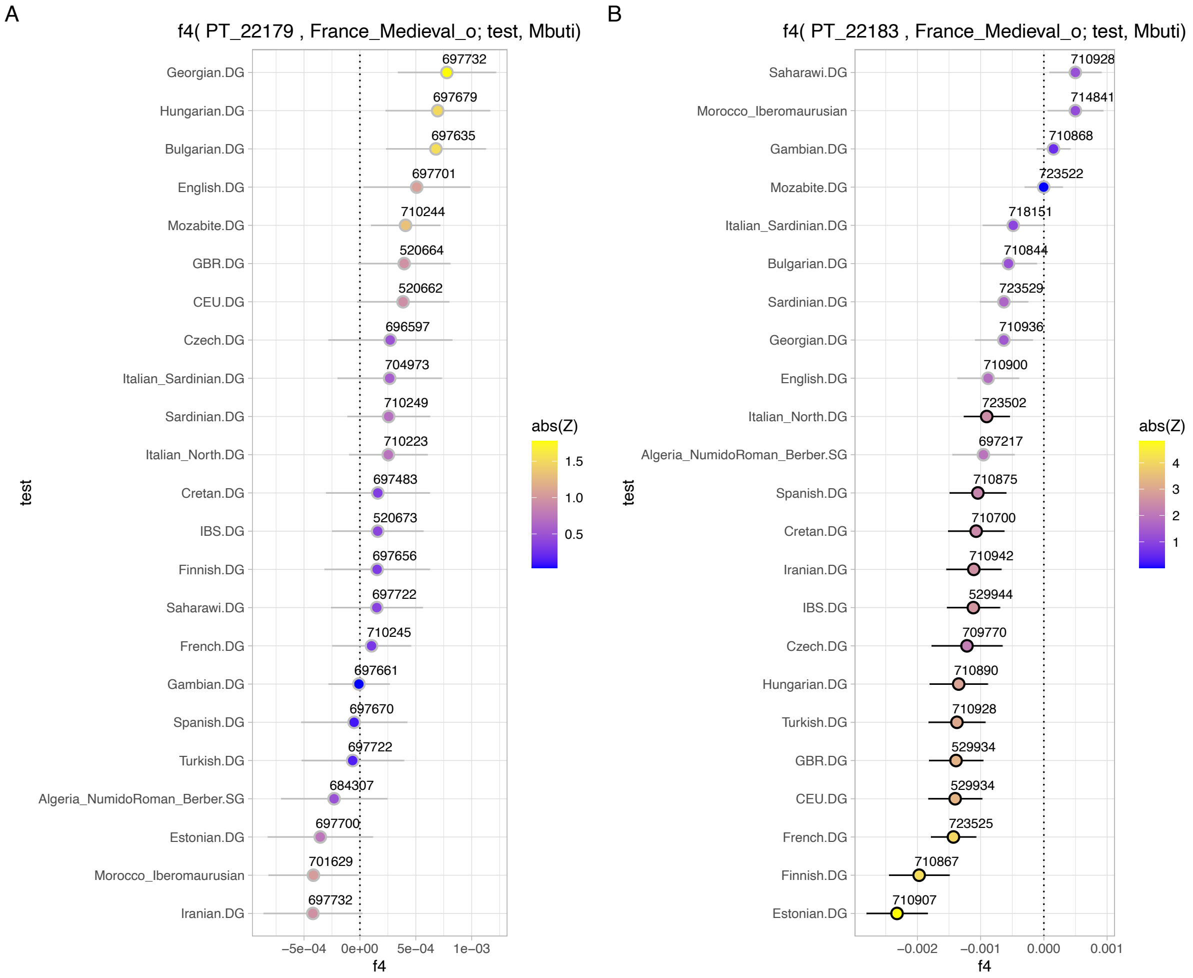

Figure S38. The x-axes represent the $f_4$-statistic values, with results displayed as the mean ± 1-SD, colors representing Z-scores and black strokes Z-scores > 2. The numbers above each dot indicate the number of SNPs used for each calculation. $f_4$-statistics are shown as $f_4$(X, France_Medieval_o; Test, Mbuti) with *Test* including Eurasian and African populations from the Medieval period or proxies and *X* being (A) PT_22179 and (B) PT_22183.
